# Single-cell lineages reveal the rates, routes, and drivers of metastasis in cancer xenografts

**DOI:** 10.1101/2020.04.16.045245

**Authors:** Jeffrey J. Quinn, Matthew G. Jones, Ross A. Okimoto, Shigeki Nanjo, Michelle M. Chan, Nir Yosef, Trever G. Bivona, Jonathan S. Weissman

## Abstract

Cancer progression is characterized by rare, transient events which are nonetheless highly consequential to disease etiology and mortality. Detailed cell phylogenies can recount the history and chronology of these critical events – including metastatic seeding. Here, we applied our Cas9-based lineage tracer to study the subclonal dynamics of metastasis in a lung cancer xenograft mouse model, revealing the underlying rates, routes, and drivers of metastasis. We report deeply resolved phylogenies for tens of thousands of metastatically disseminated cancer cells. We observe surprisingly diverse metastatic phenotypes, ranging from metastasis-incompetent to aggressive populations. These phenotypic distinctions result from pre-existing, heritable, and characteristic differences in gene expression, and we demonstrate that these differentially expressed genes can drive invasiveness. Furthermore, metastases transit via diverse, multidirectional tissue routes and seeding topologies. Our work demonstrates the power of tracing cancer progression at unprecedented resolution and scale.

**One Sentence Summary:** Single-cell lineage tracing and RNA-seq capture diverse metastatic behaviors and drivers in lung cancer xenografts in mice.

## Main Text

Cancer progression is governed by evolutionary principles (reviewed in (*1*)), which leave clear phylogenetic signatures upon every step of this process (*2, 3*), from early acquisition of oncogenic mutations (i.e., the relationships between normal and malignantly transformed cells (*4*)), to metastatic colonization of distant tissues (i.e., the relationship between a primary tumor and metastases (*5*)), and finally adaptation to therapeutic challenges (i.e., the relationship between sensitive and resistant clones (*6*)). Metastasis is a particularly critical step in cancer progression to study because it is chiefly responsible for cancer-related mortality (*7*). Yet because metastatic events are intrinsically rare, transient, and stochastic (*8, 9*), they are typically impossible to monitor in real time. Analogous to the cell fate maps that have played an essential role in deepening our understanding of organismal development and cell type differentiation (*10, 11*), accurately reconstructed phylogenetic trees of tumors and metastases can reveal key features of this process, such as the clonality, timing, frequency, origins, and destinations of metastatic seeding (*12*).

Lineage tracing techniques allow one to map the genealogy of related cells, providing a crucial tool for exploring the phylogenetic principles of biological processes like cancer progression and metastasis. Classical lineage tracing strategies can infer tumor ancestry from the pattern of shared sequence variations across tumor subpopulations (e.g., naturally occurring mutations, like single-nucleotide polymorphisms or copy-number variations) (*13, 14*). These “retrospective” tracing approaches are particularly valuable for studying the subclonal dynamics of cancer in patient-derived samples, such as elucidating which mutations contribute to metastasis and when they occur (*15* – *18*). However, the resolution of these approaches is typically low because of the limited number of distinguishing natural mutations, and the conclusions can be confounded by incomplete or impure bulk tumor sampling (*19*), sequencing artifacts (*20*), varying levels of intratumor heterogeneity, and non-neutral mutations (*1, 5*). Alternatively, so-called “prospective” lineage tracing approaches – wherein cells are marked with a static label (e.g. genetic barcode or fluorescent tag) – can measure gross population dynamics at *clonal* resolution (*21*), but cannot resolve important and fine *subclonal* features of cancer biology, like evolution and the rate, timing, and directionality of metastatic events.

The recent development of Cas9-enabled lineage tracing techniques with single-cell RNA-sequencing readouts (*22−26*) provides the potential to explore cancer progression at vastly larger scales and finer resolution than has been previously possible with classical prospective or retrospective tracing approaches. These new methods most commonly rely on similar technical principles (reviewed in (*27, 28*)). Briefly, Cas9 targets and cuts a defined genomic locus (hereafter “Target Site”), resulting in a stable insertion/deletion (indel) “allele” that is inherited over subsequent generations; as the cells divide, they accrue more Cas9-induced indels at additional sites that further distinguish successive clades of cells (**Fig. 1A, Fig. S1**). At the end of the lineage tracing experiment, the indel alleles are collected from individual cells by sequencing and paired with single-cell expression profiles of the cell state (*22, 23*). Then, as in retrospective tracing approaches, various computational approaches ((*29 – 34*)) can reconstruct a phylogenetic tree that best models subclonal cellular relationships (e.g., by maximum-parsimony) from the observed shared or distinguishing alleles. Thus far, Cas9-enabled tracing has been successfully applied to study important aspects of metazoan biology, like the cellular progenitor landscape in early mammalian embryogenesis (*23, 35*), hematopoiesis (*36*), and neural development in zebrafish (*22*). Additionally, resources now exist for studying other phylogenetic processes in mouse (*23, 35*), and analytical tools are available for computationally reconstructing and benchmarking trees from large lineage tracing datasets (*34, 37*).

**Fig. 1.**
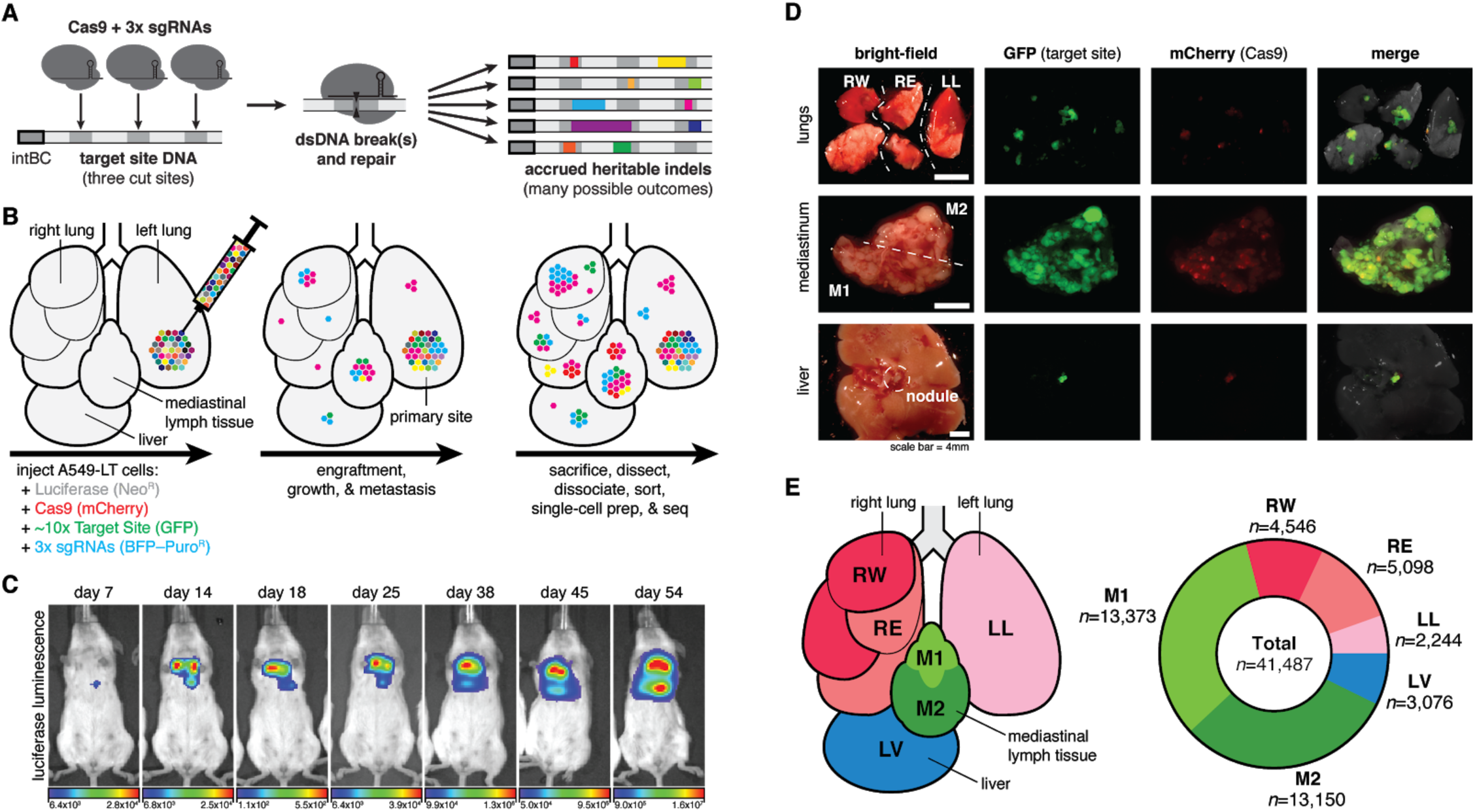
Lineage tracing in a lung cancer xenograft model in mice. (**A**) Our Cas9-enabled lineage tracing technology. Cas9 and three sgRNAs bind and cut cognate sequences on genomically integrated Target Sites, resulting in diverse indel outcomes (multicolored rectangles), which act as heritable markers of lineage. (**B**) Xenograft model of lung cancer metastasis. Approximately 5,000 A549-LT cells were surgically implanted into the left lung of immunodeficient mice. The cells engrafted at the primary site, proliferated, and metastasized within the five lung lobes, mediastinal lymph, and liver. (**C**) *In vivo* bioluminescence imaging of tumor progression over 54 days of lineage recording, from early engraftment to widespread growth and metastasis. (**D**) Fluorescent imaging of collected tumorous tissues. (**E**) Anatomical representation of the six tumorous tissue samples (**left**), and the number of cells collected with paired single-cell transcriptional and lineage datasets (**right**).

Here we apply lineage tracing to explore the subclonal dynamics of metastatic dissemination in an orthotopic xenograft model of lung cancer in mice (*38*). Specifically, we have modified our previously described “molecular recorder” for lineage tracing (*23*), now enabling the capture of highly detailed, single-cell-resolution phylogenies across tens of thousands of cells with continuous tracing *in vivo* over several months. Additionally, we have expanded on our analytical toolkit, Cassiopeia (*34*), with algorithms for inference of unobserved events from a phylogeny, which we applied here to resolve metastatic transitions between tissues. These and other advances allowed us to study the rates, transcriptional drivers, and tissue routes of metastasis at unprecedented scale and resolution.

### Tracing metastasis in a mouse xenograft model

We chose to study metastasis using a human *KRAS*-mutant lung adenocarcinoma line (A549 cells) in an orthotopic xenograft model in mice because this system is characterized by aggressive metastases (*38*) and orthotopic xenografting experiments such as this are useful for modeling cancer progression *in vivo (39)*. We engineered A549 cells with a refined version of our molecular recorder technology (*23*) (**Fig. S2**; Methods). Specifically, the engineered cells contained: (i) luciferase for live imaging; (ii) Cas9 for generating heritable indels; (iii) ∼10 uniquely barcoded copies of the Target Site for recording lineage information, which can be captured as expressed transcripts by single-cell RNA-sequencing; and finally (iv) triple-sgRNAs to direct Cas9 to the Target Sites, thereby initiating lineage recording (**Fig. 1A; Fig. S2A–C**). To enable tracing over months-long timescales, we carefully designed the sgRNAs with nucleotide mismatches to the Target Sites, thereby decreasing their affinity (*40, 41*) and tuning the lineage recording rate (*23, 42*). Approximately 5,000 engineered cells (“A549-LT”) were then embedded in matrigel and surgically implanted into the left lung of an immunodeficient (C.B-17 *SCID*) mouse (**Fig. 1B**). We followed bulk tumor progression by live luciferase-based imaging (**Fig. 1C**): early bioluminescent signal was modest and restricted to the primary site (left lung), consistent with engraftment; with time, the signal progressively increased and spread throughout the thoracic cavity, indicating tumor growth and metastasis. After 54 days, the mouse was sacrificed and tumors were identified in the five lung lobes, throughout the mediastinal lymph tissue, and on the liver (**Fig. 1D**), in a pattern that is consistent with previous studies in this model (*38*). From these tumorous tissues, we collected six samples, including one from the left lung (i.e., including the primary site; **Fig. 1E, left**). The tumor samples were dissociated, fluorescence-sorted to exclude normal mouse cells, and finally processed for single-cell RNA-sequencing. To simultaneously measure the transcriptional states and phylogenetic relationships of the cells, we prepared separate RNA expression and Target Site amplicon libraries, respectively, resulting in 41,487 paired single-cell profiles from six tissue samples (**Fig. 1E, right**; **Fig. S3**; Methods).

In addition to the mouse described above (hereafter “M5k”), we also performed lineage tracing in three other mice (called “M10k”, “M100k”, and “M30k”), using A549-LT cells engineered with slightly different versions of the lineage tracing technology (**Fig. S4;** Methods). Unless otherwise noted, we focus our primary discussion of the results on mouse M5k because it yielded the richest lineage tracing dataset with the most cells and distinct lineages.

### Distinguishing clonal cancer populations

Our lineage recorder “Target Site” (*23*) carries two orthogonal units of lineage information: (i) a static 14bp-randomer barcode (“intBC”) that is unique and distinguishes between multiple integrated Target Site copies within each cell; and (ii) three independently evolving Cas9 cut-sites per Target Site that record heritable indel alleles and are used for subclonal tree reconstruction (**Fig. 1A**). Each Target Site is expressed from a constitutive promoter allowing it to be captured by single-cell RNA-sequencing. After amplifying and sequencing the Target Site mRNAs, the reads were analyzed using the Cassiopeia processing pipeline (*34*). Briefly, this pipeline leverages unique molecular identifier (UMI) information and redundancy in sequencing reads to confidently call intBCs and indel alleles from the lineage data, which inform subsequent phylogenetic reconstruction (**Fig. S1**; Methods).

We first determined the number of clonal populations (that is, groups of related cells that descended from a single clonogen at the beginning of the xenograft experiment), which are each associated with a set of intBCs. Importantly, the A549-LT cells were prepared at high diversity such that clones carry distinct intBC sets. By sampling the A549-LT cells before implantation, we estimate that the implanted pool of 5,000 cells initially contained 2,150 distinguishable clones (**Fig. S2D**). Based on their intBC sets, we assigned the vast majority of cancer cells collected from the mouse (97.7%) to 100 clonal populations (**Figs. S5A-B**), ranging in size from >11,000 (Clone #1, “CP001”) to ∼30 cells (CP100) (**Fig. S5C**). Though there were some smaller clonal populations, we focused on these largest 100 because lineage tracing in few cells is less informative. Furthermore, despite initially implanting ∼2,150 distinct clones, only ∼100 clones successfully engrafted and proliferated; this indicates that only a small minority of cells may be competent for engraftment and survival *in vivo* (**Fig. S2D**). Moreover, we find no correlation between initial (pre-implantation) and final (post-sacrifice) clonal population size (Spearman’s ρ=-0.026; **Fig. S2E**), suggesting that clone-intrinsic characteristics that confer greater fitness *in vitro* do not necessarily confer greater fitness *in vivo* (*43, 44*).

Features that influence the lineage recording capacity and tree reconstructability differed between clonal populations, such as the copy-number of Target Sites, the percentage of recording sites bearing indel alleles, and allele diversity (**Fig. S6A-C; Fig. S7**). Though most clonal populations exceeded parametric standards for confident phylogenetic reconstruction, some had slow recording kinetics or low allele diversity and failed to pass quality-control filters (17 clones, 7.3% of total cells in mouse M5k, **Fig. S6D; Fig. S7B**); these clones were excluded from tree reconstruction and downstream analyses (Methods).

We observed that the clonal populations exhibited distinct distributions across the six tissues (**Fig. S8A-C**), ranging from being present exclusively in the primary site (e.g., CP029, CP046), to overrepresented in a tissue (CP003, CP020), or distributed broadly over all sampled tissues (CP002, CP013). The level of tissue dispersal is a consequence of metastatic dissemination and thus can inform on the frequency of past metastatic events as follows: clonal populations that reside exclusively in the primary site likely never metastasized; those that did not broadly colonize tissues likely metastasized rarely; and those with more broad dispersal across all tissues likely metastasized more frequently. To quantify the relationship between tissue distribution and metastatic phenotype, we defined a statistical measure of the observed-versus-expected tissue distributions of cells (termed “Tissue Dispersion Score”; Methods) to operate as a coarse, tissue-resolved approximation of the metastatic rate. Across the 100 clonal populations in this mouse, we observed a wide range of Tissue Dispersion Scores (**Fig. S8D**), suggesting broad metastatic heterogeneity across the tumor populations. We next explored this suggested metastatic heterogeneity more directly and at far greater resolution using the evolving lineage information.

### Single-cell-resolved cancer phylogenies

The key advantage of our lineage tracer is not in following *clonal* lineage dynamics (i.e. from cells’ static intBCs, as described in the section above) but rather in reconstructing *subclonal* lineage dynamics (i.e. from cells’ continuously evolving indel alleles). As such, we next reconstructed high-resolution phylogenetic trees using the Cassiopeia suite of phylogenetic inference algorithms (*34*) with modified parameters tailored to this dataset’s unprecedented complexity and scale (Methods). Each of the resulting trees comprehensively describes the phylogenetic relationships between all cells within the clonal population (**Fig. 2A**), thus summarizing their life histories from initial, clonogenic founding in the mouse to final dissemination across tissues and tumors. The trees are intricately complex (mean tree depth of 7.25; **Fig. S6E**) and highly resolved (consisting of 37,888 cells with 33,266 (87.8%) unique lineage states; **Fig. S6C**).

**Fig. 2.**
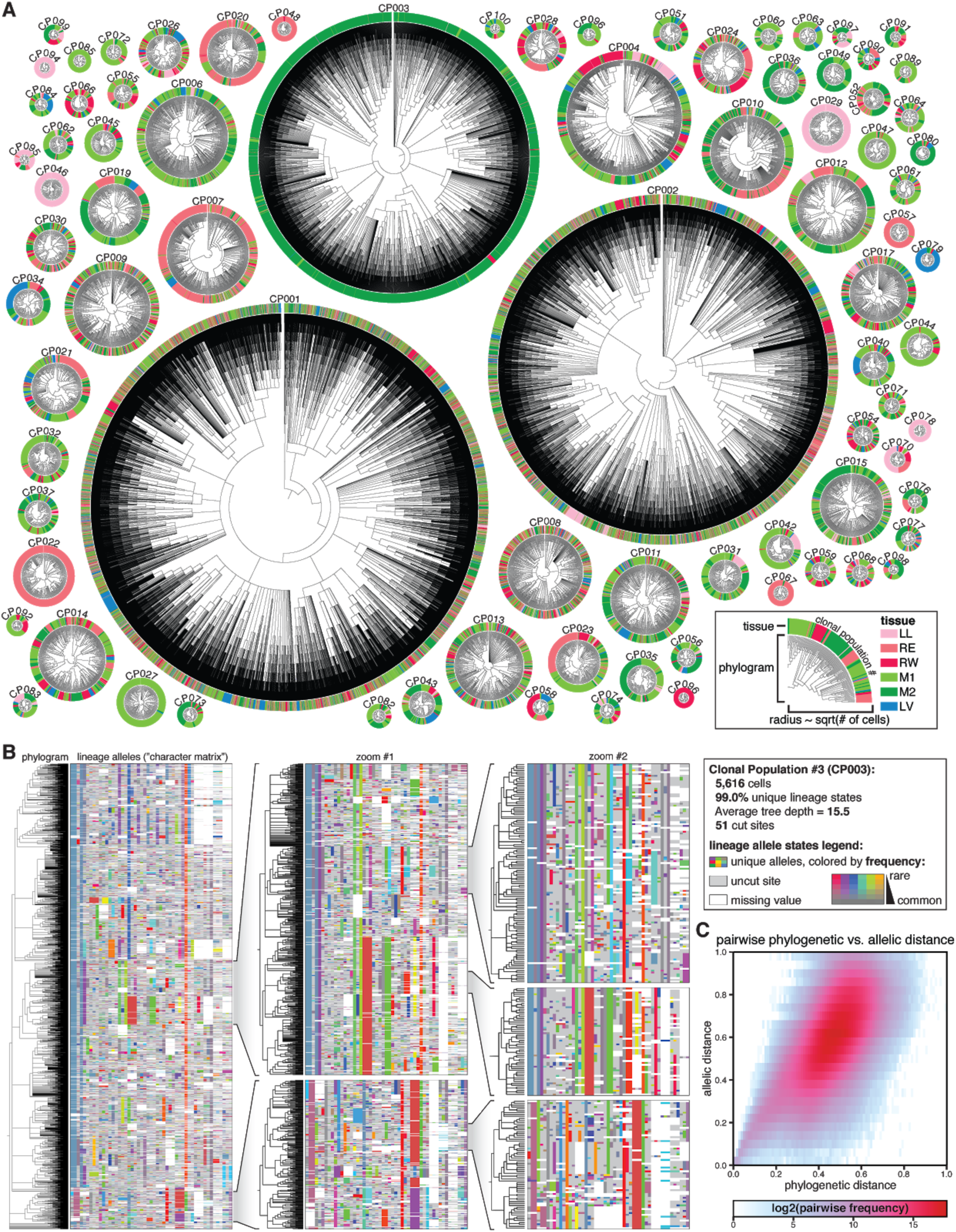
High-resolution phylogenetic trees capture the histories of clonal cancer populations. (**A**) Highly detailed phylogenetic reconstructions for each clonal population, represented as radial phylograms. Each cell is represented along the circumference and colored by tissue, as in **Fig. 1E** and legend. Trees differ in size, tissue distribution, and frequency of tissue transitions. Each tree is scaled by the square-root of the number of cells. (**B**) Phylogenetic tree and lineage alleles of one clonal population (CP003; *N*=5,616 cells). The phylogram (**left**) represents cell–cell relationships and the matrix (**right**) represents the lineage alleles for each cell. Alleles are uniquely colored, where saturation indicates allele rarity (legend). (**B, inlays**) Nested zooms of individual clades show the patterns of shared and distinguishing indel alleles, and highlight indel diversity, tree depth, and tree complexity. (**C**) Correspondence between phylogenetic distance (the normalized pairwise tree distance between two cells) and allelic distance (the normalized pairwise difference in alleles between two cells) for CP003, indicating that the tree accurately models phylogenetic relationships.

To illustrate the intricate complexity of the trees in this dataset, we present the reconstructed phylogram and lineage alleles for a representative clonal population of 5,616 cells (CP003; **Fig. 2B**) with 99.0% (5,560) unique cell lineage states, mean tree depth of 10.0, and maximum tree depth of 20. Intuitively, cells that are more closely related to one another ought to share more lineage alleles, which is evident from the patterns of shared alleles within clades and distinguishing alleles between clades (**Fig. 2B, zoomed inlays**). Indeed, we find systematic agreement between phylogenetic distance (i.e., the distance between two cells in the tree) and allelic distance (the difference between two cells’ lineage alleles) for this example (**Fig. 2C**) and across all other trees (**Fig. S10**), thus supporting their accuracy. The high diversity of distinguishable Cas9-induced indels (9,936 unique alleles across all M5k cells; evident in the array of unique allele colors in **Fig. 2B**) also reduces the probability of homoplasy, an issue which complicates tree reconstruction and impairs tree accuracy (*34, 45*). Altogether, these features indicate that the reconstructed trees accurately model the true phylogenetic relationships between cells.

### Inferring and quantifying past metastatic events from phylogenies

A striking feature revealed by the reconstructed phylogenies is the varying extent to which closely related cells reside in different tissues (**Fig. 2A**), patterns which directly result from ancestor cells having physically transited from one tissue to another in the past (i.e., metastatic seeding). Varying rates of metastasis produce different patterns of concordance between phylogeny and tissue (**Fig. 3A**). For example, non-metastatic populations result in all clades remaining within a single tissue (**Fig. 3A-B, left**); conversely, highly metastatic populations result in closely related cells residing in different tissues (**Fig. 3A-B, right**). Finally, intermediate levels of metastasis can similarly lead to a dispersed tissue distribution as in the highly metastatic regime, though with fewer metastatic transitions, thus supporting the need to reconstruct trees in order to distinguish such cases (**Fig. 3A-B, middle**).

**Fig. 3.**
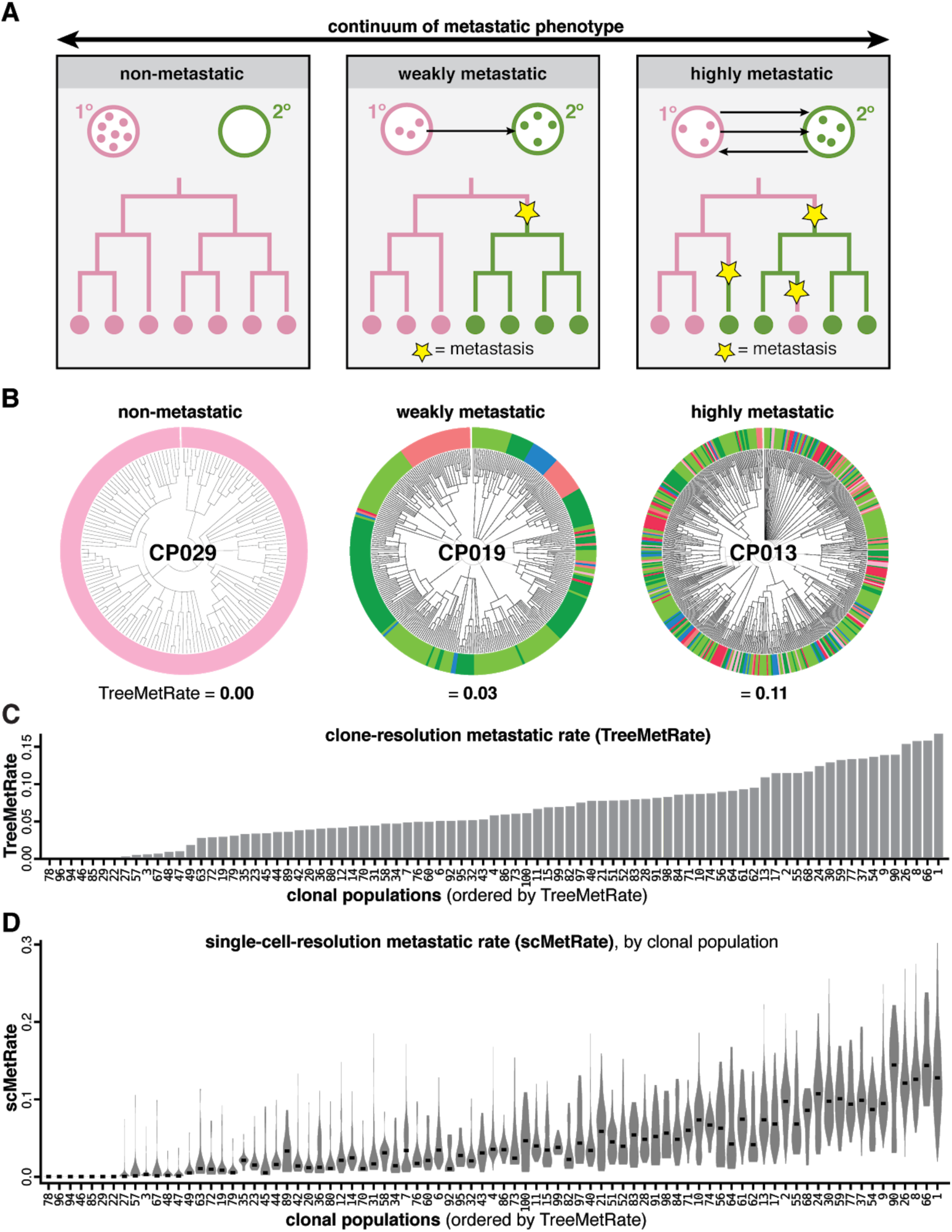
Quantifying the diverse metastatic phenotypes of clonal populations directly from cell lineages. (**A**) Theoretical continuum of metastatic phenotypes, spanning non-metastatic (never exiting the primary site) to highly metastatic (frequently transitioning between tumors; arrows). Ancestral metastatic events between tissues leave clear phylogenetic signatures (yellow stars). (**B**) Example clonal populations that illustrate the wide range of metastatic phenotypes observed: a non-metastatic population that never exits the primary site (CP029); a moderately metastatic population that infrequently transitions between different tissues (CP019); and a frequently metastasizing population with closely related cells residing in different tissues (CP013). Cells colored by tissue as in **Fig. 1E**; metastatic phenotypes scored by the TreeMetRate. (**C**) The distribution of TreeMetRates for each clonal population. (**D**) The distributions of single-cell-resolution metastatic phenotypes (scMetRates) for each clonal population, rank-ordered by TreeMetRate; median scMetRate indicated in black.

To quantitatively study the relationship between metastatic phenotype and phylogenetic topology, we used the Fitch-Hartigan maximum parsimony algorithm (*46, 47*). Our implementation of this algorithm provides the minimal number of ancestral (i.e., not directly observed) metastatic transitions that are needed to explain the final (i.e., observed) tissue location of each cell in a given tree. We defined a score of the metastatic potential (termed “TreeMetRate”) by dividing the inferred minimal number of metastatic transitions by the total number of possible transitions (i.e., edges in the tree). Empirically, we observe a distribution of clonal populations that spans the full spectrum of metastatic phenotypes between low (non-metastatic) and high (very metastatic) TreeMetRates (**Fig. 3B,C**). The TreeMetRate is stable across bootstrapping experiments in simulated trees (**Fig. S9E-F**) and when using an alternative phylogenetic reconstruction method (Neighbor-Joining (*29*)) on empirical data (**Fig. S11A**; Pearson’s *ρ*=0.94), indicating that the TreeMetRate is a robust measurement of metastatic behavior – though, notably, Cassiopeia trees are more parsimonious than those reconstructed by Neighbor-Joining (**Fig. S11B**). Empirically, the Tissue Dispersal Score agrees with the TreeMetRate at low metastatic rates (**Fig. S12A,C**), however, the TreeMetRate more accurately captures the underlying metastatic rate over a broad range of simulated metastatic rates because it can distinguish between moderate and high metastatic rates (**Fig. S9D**), which both result in broad dispersion across tissues (**Fig. 3A**), whereas the Tissue Dispersal Score saturates at intermediate metastatic rates (**Fig. S9B**). Furthermore, the TreeMetRate also agrees with the probability that a cell’s closest relative (by lineage allele similarity) resides in a different tissue for each clonal population (termed “AlleleMetRate”; **Fig. S12B, D**); importantly, the AlleleMetRate is an alternative metric of metastatic potential that exploits the evolving nature of our lineage tracer but is independent of tree reconstruction. Again, however, simulations indicate that the TreeMetRate is the superior measurement of the underlying metastatic rate (**Fig. S9A-D**), underscoring the value of the reconstructed phylogenies in helping identify aspects of metastatic behavior that would otherwise be invisible.

We further extended our parsimony-based approach to quantify the metastatic phenotype at the resolution of individual cells (termed the “scMetRate”) by averaging the TreeMetRate for all subclades containing a given cell (Methods). This measurement is sensitive to *subclonal* differences in metastatic behavior (**Fig. 3C**), and highlighted intriguing bimodal metastatic behavior for clone CP007 (**Fig. 5G;** discussed below). Additionally, we find that the scMetRate is uncorrelated to clonal population size, proliferation signatures (*48, 49*), or cell cycle stage (*50*) (**Fig. S13**), indicating that it can measure metastatic potential uncoupled from proliferative capacity. Overall, these results indicate that cancer cells in this dataset exhibit diverse metastatic phenotypes both between and within clonal populations, which can be meaningfully distinguished and quantified by virtue of the lineage tracer, but would have otherwise been hidden from classical barcoding approaches.

### Transcriptional drivers of differences in metastatic phenotype

A central question in cancer biology is the extent to which cellular properties (e.g., transcriptional state) underlie cancer phenomena (*51*), like metastatic capacity. By comparing the paired transcriptional and lineage datasets, we found that different metastatic behaviors corresponded to differential expression of genes, many with known roles in metastasis. First, after filtering and normalizing the scRNA-sequencing data, we applied *Vision (52)*, a tool for assessing the extent to which the variation in cell-level quantitative phenotypes can be explained by transcriptome-wide variation in gene expression. While we found little transcriptional effect attributable to clonal population assignment, we found a modest association between a cell’s transcriptional profile and its tissue sample or metastatic rate (**Fig. S14**). We next performed pairwise differential expression analyses comparing cells from completely non-metastatic clonal populations (i.e., four clones that never metastasized from the primary tissue in the left lung, like CP029) to metastatic clones in the same tissue (**Fig. S15**). This clone-resolution analysis identified several genes with significant expression changes which were also consistent across each non-metastatic clone (log2 fold-change > 1.5, FDR < 0.01), such as IFI6. These initial results suggested that differences in metastatic phenotype may manifest in characteristic differences in gene expression, and motivated deeper analysis.

Next, we sought to comprehensively identify genes that are associated with metastatic behavior by regressing single-cell gene expression against the scMetRates (over all observed cells, clonal populations, and tissues; **Fig. 4A**; Methods), thereby leveraging both the scRNA-seq dataset and the single-cell phylogenies. Many of the identified positive metastasis-associated candidates (i.e., genes with significantly higher expression in highly metastatic cells) have known roles in potentiating tumorigenicity (**Fig. 4B, top**). For example, IFI27 is an interferon-induced factor that is anti-apoptotic and promotes epithelial-mesenchymal transition (EMT), cell migration, and cancer stemness in various carcinomas *(53, 54*); REG4 enhances cell migration and invasion in colorectal carcinoma *(55*) and KRAS-driven lung adenocarcinoma (*56*); and TNNT1 has elevated expression in many cancers and may promote EMT and invasiveness (*57*). Similarly, many negative metastasis-associated candidates (i.e., genes with significantly lower expression in highly metastatic cells) have known roles in attenuating metastatic potential (**Fig. 4B, bottom**). For example, NFKBIA (IκBα) is a pan-cancer tumor suppressor via inhibition of pro-tumoral NFκB signaling (*58*); lower expression of ID3 enhances tumor cell migration and invasion *in vitro* and in lung adenocarcinoma xenograft models (*59*); and downregulation of ASS1 supports tumor metabolism and proliferation (*60*). (Interestingly, our most significant negative candidate was KRT17, which has previously been implicated in *promoting* invasiveness in lung adenocarcinoma (*61*) and its overexpression has been associated with poor prognosis in some cancers (*62*); we follow-up on this unexpected finding below.) Additionally, many of the identified genes were significantly reproduced across every mouse in this study (**Fig. 4C-D**; **Fig. S17**). And more generally, the gene-level expression trends are broadly supported by significant correlation between the TreeMetRate and several gene expression signatures (*63*) (**Fig. S16**), including interferon signaling programs (*64*), RAS pathways (*65*) (A549 cells are *KRAS*-mutant), cancer invasiveness (*66*), *and EMT (67*) *(consistent with increased NFκB signaling (68, 69*)).

**Fig. 4.**
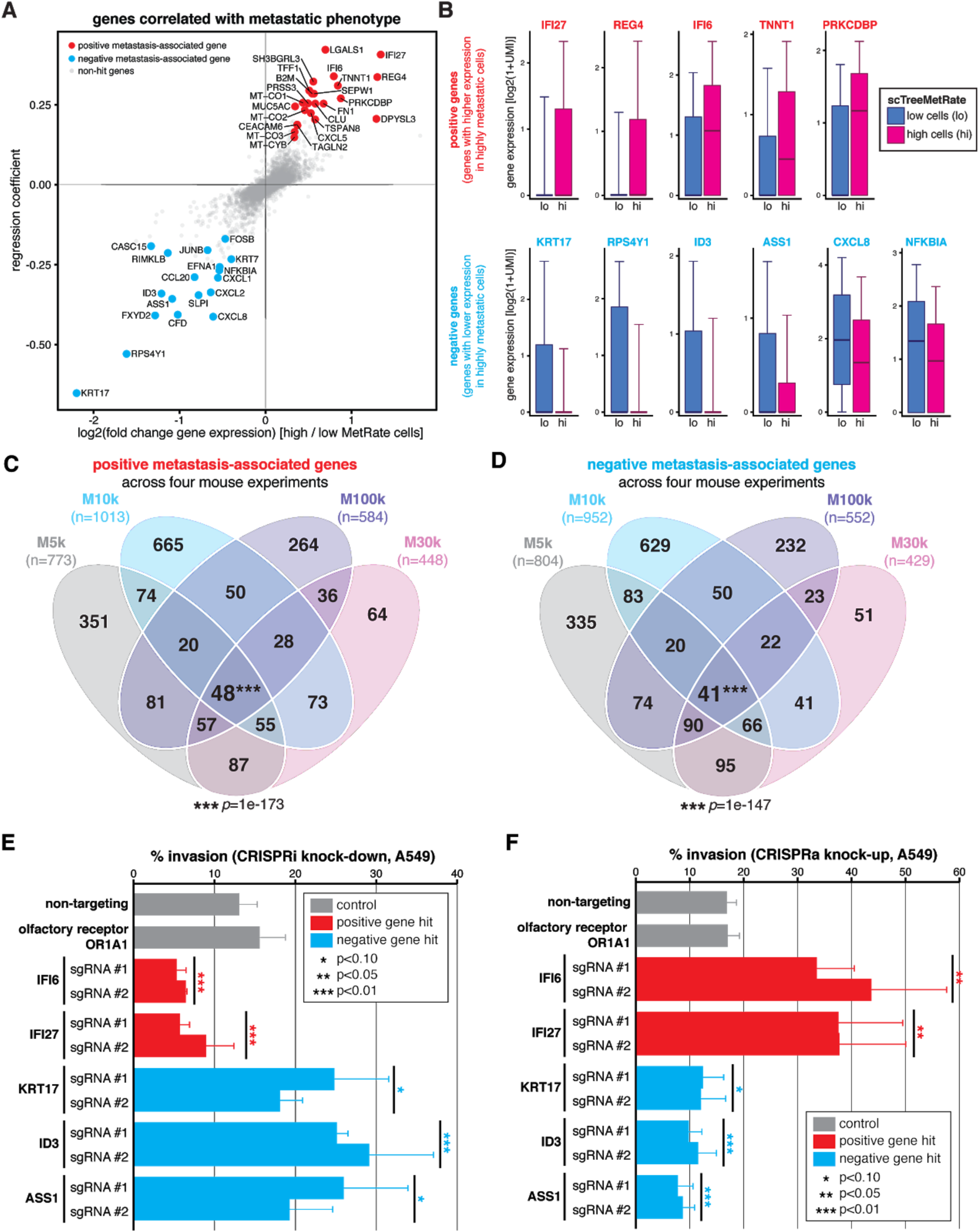
Divergent metastatic phenotypes are driven by differences in gene expression. (**A**) Poisson regression analysis of single-cell gene expression and scMetRate for all cells and all tissues; fold-change and coefficient of regression shown. The strongest and most significant positive and negative genes are annotated (red and blue, respectively; Methods). (**B**) Expression level of several positive and negative metastasis-associated gene candidates (top and bottom rows, respectively) in cells with low or high scMetRate (blue and magenta box-plots, respectively). Boxes: first, second, and third quartiles; whiskers: 9th and 91st percentiles of expression distribution. (**C** and **D**) Overlap of identified positive and negative metastasis-associated genes, respectively, from the four mouse experiments; number of genes indicated. Four-way intersections between gene sets are significant by *SuperExactTest* (*87*) *multi-set intersection test. (****E*** *and* ***F****) In vitro* transwell invasion assays following CRISPRi or CRISPRa gene perturbation, respectively, in A549 cells. Perturbations were performed using two independent sgRNAs per gene. Differences in invasion phenotype relative to two negative control guides (non-targeting and olfactory receptor) were significant by two-tailed *t*-test; error bars show standard deviation across triplicates.

**Fig. 5.**
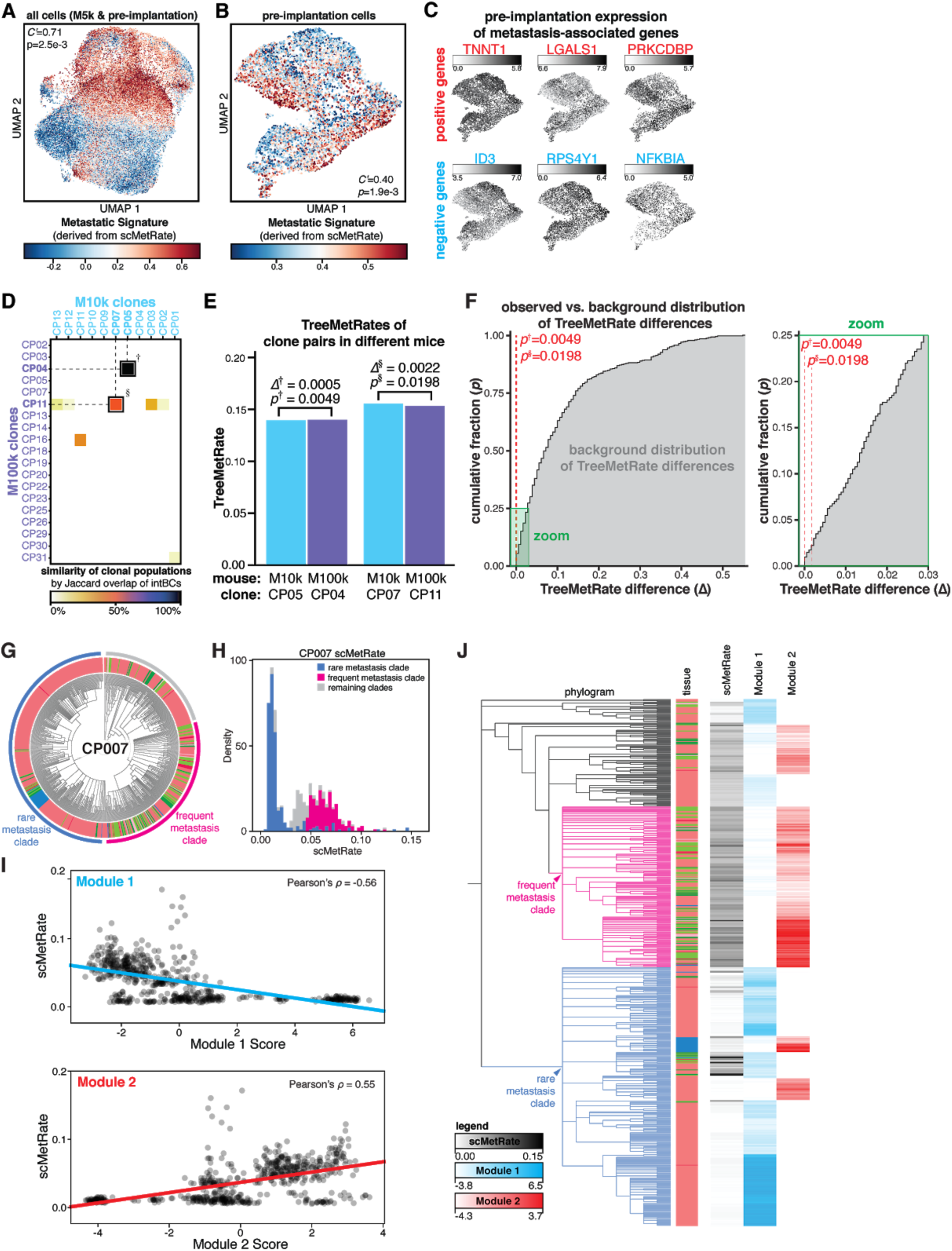
Metastatic phenotype is predetermined, heritable, and reproducible. (**A** and **B**) Projections of transcriptional states of M5k cancer cells and pre-implantation cells (**A**) or pre-implantation cells alone (**B**), colored by Metastatic Signature. Association between transcriptional state and Metastatic Signature is measured by inverted Geary’s *C’* and significance by false discovery rate (*p*). (**C**) Pre-implantation cells exhibit heterogeneity in expression of metastasis-associated genes. (**D**) Jaccard overlap of intBC sets between clonal populations in M10k and M100k mice. Two pairs of clonal populations (indicated by † and §) were related between the two mouse experiments (Jaccard overlap>50%). (**E**) Comparison of TreeMetRates from related clones implanted in M10k and M100k, showing minimal difference in metastatic rate (Δ) between clone pairs. (**F**) Cumulative distribution plot of the background distribution of all possible pairwise TreeMetRate differences between M10k and M100k clones (gray), with zoom to show low-Δ regime. Both of the observed differences are statistically smaller than expected (*p*^†^=0.0049 and *p*^§^=0.0198; red dashes). (**G**) Divergent subclonal metastatic behavior exhibited in the phylogenetic tree of clonal population #7, with annotated subclades; cells colored by tissue as in **Fig. 1E**. (**H**) The bimodal distribution of scMetRates for cells in CP007, with cells from the divergent subclades indicated. (**I**) Comparison of single-cell metastatic phenotype and *Hotspot* transcriptional module scores. (**J**) Overlay illustrating concordance between CP007 phylogeny, scMetRates, and *Hotspot* Module scores.

While we identified many interesting and reproducible gene candidates in our regression analysis above, it was unclear whether they were directly driving the metastatic phenotype or were merely associated with it. To address this important point, we next explored the functional impact on metastatic behavior of modulating the expression of five high-scoring gene candidates (IFI6, IFI27, KRT17, ID3, and ASS1). First, we engineered A549 cells to enable CRISPR-inhibition or -activation perturbations (CRISPRi/a; activity validated in **Fig. S18C,D)**, then increased or decreased expression, respectively, of the five gene targets using two independent sgRNAs per gene. Finally, we measured the perturbed cells’ invasion phenotype *in vitro* using a transwell invasion assay (**Fig. 4E,F**; Methods). As hypothesized, CRISPRi knock-down resulted in *decreased* invasiveness for positive metastasis-associated genes (IFI6 and IFI27; *p=*0.001, 0.005, respectively) and *increased* invasiveness for negative metastasis-associated genes (KRT17, ID3, and ASS1; *p=*0.054, 0.003, and 0.062, respectively; **Fig. 4E**). Conversely, we found that elevating candidate gene expression by CRISPRa produced the exact opposite results (**Fig. 4F**), indicating that the invasion phenotype can be quantitatively altered by both increased or decreased expression for each of the five candidate genes tested, including notably KRT17. We confirmed that the modulation of expression of each of these genes strongly and significantly modulated invasiveness (*p*<0.01) in a separate human lung cancer cell line (H1299 cells, which are *KRAS* wild-type, *TP53*-mutant, and harbor endogenous *NRAS*^*Q61K*^; **Fig. S18A,B**); though, for two of the genes (IFI27, IFI6), CRISPRa had a significant effect (*p*<0.01) while CRISPRi did not. Taken together, these results indicate that (i) the lineage tracer can meaningfully identify metastasis-associated genes *in vivo*, (ii) some of these gene candidates are sufficient to drive differences in metastatic phenotype, and (iii) these genes’ roles in mediating invasiveness extend beyond the one A549 cancer model and across different oncogenic backgrounds.

### Heterogeneity and heritability of metastatic behavior in pre-implantation cells

We next used the positive and negative metastasis-associated genes identified above (**Fig. 4A**) to define a *de novo* transcriptional signature (hereafter, “Metastasis Signature”; **Fig. 5A**; **Fig. S19A**). We found that even prior to implantation into the mice, the cells already exhibited meaningful heterogeneity in the Metastatic Signature (**Fig. 5B**), and metastasis-associated genes like ID3 and TNNT1 were similarly heterogeneously expressed pre-implantation (**Fig. 5C**). Next, we used the lineage barcodes to map cells from the *in vitro* pre-implantation pool to the clonal populations that engrafted *in vivo* (**Fig. S19B**). We then segregated these mapped cells into the top and bottom halves by their corresponding TreeMetRate, and queried their pre-implantation Metastatic Signatures. We found that cells from more metastatic clones in the mouse had modestly yet significantly higher metastatic signatures prior to implantation, and vice versa (**Fig. S19C**). This indicates that the pre-implantation transcriptional signature is mildly predictive of *in vivo* metastatic phenotype (**Fig. S19D**), though the distinction becomes more amplified *in vivo* (**Fig. S19C, D**). This result suggests that even before cells were xenografted into the mouse, they were primed for greater or lesser metastatic capacity *in vitro*.

While the pre-existing transcriptional heterogeneity in the pre-implantation cells was noteworthy, it remained unclear whether these differences were stochastic or intrinsic properties of the cells that could be robustly propagated *in vitro* and *in vivo*. One way to address this question is by implanting two cells from the same clone into two distinct mice and querying how well their metastatic phenotype is reproduced. Using the cells’ intBCs, which statically mark clones, we identified two such instances where cells from the same clonal population seeded tumors in two different mice (**Fig. 5D; Fig. S20**). Strikingly, for each of the two pairs of clonal populations, the TreeMetRates were nearly identical (**Fig. 5E**). In fact, one of these pairs had the most similar TreeMetRates across all pairs of clones in the two mouse experiments (Δ(TreeMetRate)=0.0005, *p*=0.0049; **Fig. 5F**). Taken together, these results indicate that (i) the diverse metastatic phenotypes *in vivo* are determined before implantation (also **Fig. 5B, C**), (ii) the metastatic phenotype is reproducible over generations and is thus heritable (**Fig. 5E,F**; also **Fig. S19C,D**), and (iii) our analytical approaches for quantifying the metastatic rate, including reconstruction of the phylogenies, are experimentally robust (**Fig. 5E**).

### Evolution of metastatic phenotype

Though we have thus far discussed how metastatic phenotype is clone-intrinsic and stably inherited, we identified a single example within the dataset that was the exception to this general rule. Specifically, Clone #7 (CP007) exhibited exceptionally distinct subclonal metastatic behaviors, wherein one clade metastasized frequently to other tissues while another clade remained predominantly in the right lung (**Fig. 5G**). This distinction is reflected in a bimodal distribution of scMetRates (**Fig. 3C**; **Fig. 5H**). To explore the relationship between subclonal structure and gene expression, we applied *Hotspot* (*70*) and identified two modules of correlated genes that exhibit heritable expression programs (**Fig. S21A**). Strikingly, the cumulative expression of genes in Module 1 is correlated with lower metastatic rates, while the opposite holds for Module 2 (**Fig. 5I; S21B,C**). Consistently, the two modules broadly correspond to the two clades with diverging metastatic phenotypes (**Fig. 5J**). This result is reproduced even in a control analysis of CP007 cells from the right lung only (**Fig. S21D-G**), indicating that these differences in gene expression indeed reflect differences in metastatic phenotype rather than tissue-specific effects. This example serves to illustrate that though the metastatic rate is stably inherited, it can also evolve – albeit rarely – within a clonal population, alongside concordant changes in transcriptional signature. Importantly, this finding could only be appreciated by virtue of the subclonal resolution of the lineage tracer.

### Tissue routes and topologies of metastasis

The phylogenetic reconstructions also made it possible to describe detailed histories about the tissue routes and the directionality of metastatic seeding. For example, the phylogenetic tree for CP095 reveals five distinct metastatic events from the left lung to different tissues, in a paradigmatic example of simple primary seeding (**Fig. 6A–B**). Other phylogenies revealed more complicated trajectories, such as CP019, wherein early primary seeding to the mediastinum was likely followed by intra-mediastinal transitions and later seeding from the mediastinum to the liver and right lung (**Fig. 6D–E**). To more systematically characterize the tissue transition routes revealed by the phylogenetic trees, we extended the Fitch-Hartigan algorithm *(46, 47*) to infer the directionality of each tissue transition (i.e., the origin and destination of each metastatic event) along a clonal population’s ancestry. Our algorithm, called *FitchCount*, builds on other ancestral inference algorithms like MACHINA (*71*) by scaling to large inputs and providing tissue transitions frequencies that are aggregated across all ancestries that satisfy the maximum parsimony criterion (Methods; Supplemental Text). Through simulation we show that *FitchCount* can accurately recover underlying transition probabilities better than a naive application of the Fitch-Hartigan algorithm (**Fig. S9G–H**; Methods), likely because the naive approach summarizes only a single optimal assignment solution, whereas *FitchCount* summarizes all optimal solutions. The resulting conditional probabilities of metastasis to and from each tissue are summarized in a tissue transition probability matrix (**Fig. 6C, F**). Notably, we found that these transition matrices are varied and distinct to each clone (**Fig. G; Fig. S22**). We next used principal component analysis (PCA) to stratify clones by their transition matrices (**Fig. 6H**) and identified descriptive features that capture differences in the metastatic tissue routes traversed by each clone (**Fig. 6I; Fig. S23**). These descriptive features include primary seeding from the left lung (as in CP095, **Fig. 6A–C**), metastasis from and within the mediastinum (CP098, **Fig. 6G, left**), or metastasis between lung lobes (CP070, **Fig. 6G**), and may reflect intrinsic differences in tissue tropism. From this feature analysis we also note that many clones primarily metastasized via the mediastinal lymph tissue (**Fig. 6H–I**), suggesting that the mediastinum may act as a nexus for metastatic dissemination in this mouse model. This observation is consistent with previous experiments in this model (*38*), bulk live imaging during tumor progression in this experiment wherein tumors appear to quickly colonize the mediastinum (**Fig. 1C**), and the terminal disease state wherein the mediastinum harbors the majority of the tumor burden (**Fig. S8**).

**Fig. 6.**
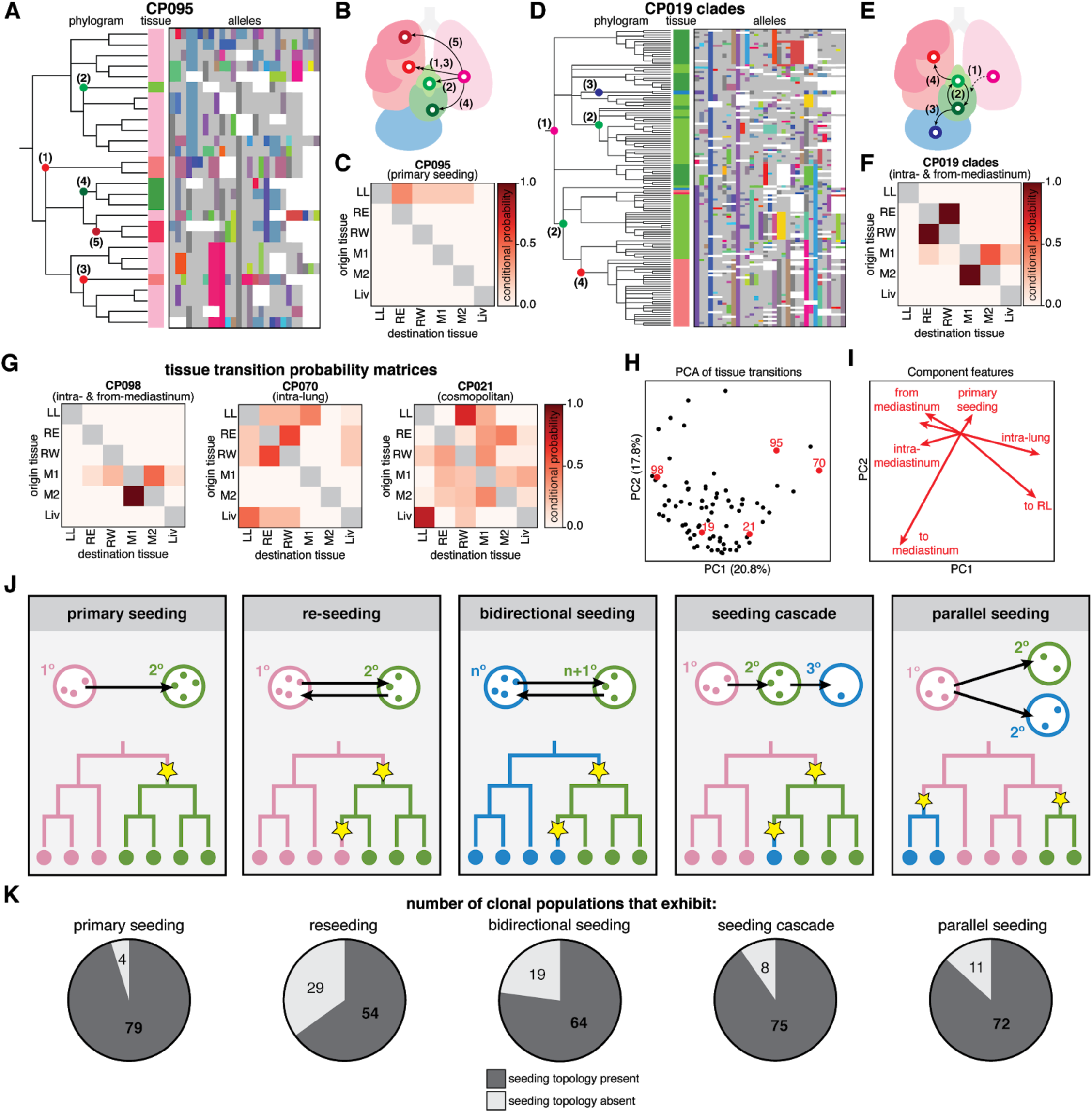
Metastases were seeded via complex tissue routes and multidirectional topologies. (**A** and **D**) Phylogenetic trees and lineage alleles for clonal population #95 and #19 clades, respectively. Notable metastatic events are annotated in the phylogram and represented graphically as arrows (**B** and **E**); cells colored by tissue as in **Fig. 1E**; lineage alleles colored as in **Fig. 2B**; dashed arrow indicates an assumed transition. (**C** and **F**) Tissue transition matrices representing the conditional probability of metastasizing from and to tissues, defining the tissue routes of metastasis for each clonal population. CP095 solely exhibits primary seeding from the left lung, whereas CP019 shows more complex seeding routes. (**G**) Tissue transition matrices illustrating the diversity of tissue routes, including metastasis from and within the mediastinum (**left**), between the lung lobes (**middle**), or amply to and from all tissues (**right**). (**H**) Principal component analysis (PCA) of tissue transition probabilities for each clonal population. Displayed clones are annotated in red; percentage of variance explained by components indicated on axes. (**I**) Component vectors of PCA with descriptive features. (**J**) Possible topologies of metastatic seeding, represented graphically and phylogenetically as in **Fig. 3A**. (**K**) Number of clonal populations that exhibit each metastatic seeding topology.

Many models of metastatic seeding topology (i.e, the sequence and directionality of metastatic transitions) have been described in cancer (*1*), including reseeding, seeding cascades, parallel seeding, and others; and each is characterized by a distinct phylogenetic signature (**Fig. 6J**). These different metastatic topologies can critically influence the progression, relapse, and treatment of cancers (*9, 72 – 74*); for example, reseeding of metastatic cells returning to the primary tumor site can contribute genetic diversity, resistance to treatment, and metastatic potential to tumors (*75, 76*). Within this single dataset, we find numerous examples of all of these topologies (**Fig. 6K**); in fact, we most often observe examples of all topologies within every clone (**Fig. S24**), as well as more complex topologies that defy simple classifications (e.g., **Fig. 6D** and **G, right**), further underscoring the aggressive metastatic nature of A549 cells in this xenograft model.

## Discussion

We applied our next-generation, Cas9-based lineage tracer to study metastasis in a lung cancer xenograft model in mice. We tracked metastatic spread at unprecedented scale and resolution that would have been unattainable by static barcoding experiments (i.e., “prospective” tracing), bulk tissue sampling, or single-cell RNA-sequencing methods alone, and were only made possible by experimental and algorithmic advances that we made on our molecular recorder platform (*23*). Experimentally, we increased lineage recorder information capacity and tuned the tracer dynamics for longer experimental timescales, thus allowing us to uniquely mark tens of thousands of cells descended from dozens of clonogens over several months. Analytically, enhancements to Cassiopeia, including *FitchCount*, allowed us to reconstruct accurate, informative, and deeply resolved phylogenetic trees, and interpret them to identify rare, transient events in cells’ ancestry (here, metastasis). Beyond the utility of this experimental and analytical framework for exploring many facets of cancer biology, we believe this tracing approach is broadly applicable to study the phylogenetic foundations of many biological processes that transpire over multiple cell generations.

When we applied this tracing strategy to a lung cancer xenograft model, several key insights emerged: (1) Single-cell lineage tracing revealed meaningful and reproducible differences in the frequency and directionality of metastatic dissemination that would not have been discernible from bulk experiments (e.g., gross distribution of clones across tumorous tissues). For example, even among clonal populations that are broadly disseminated across tissues, the lineage tracer reveals substantive differences in the underlying metastatic rates. (2) Even within a single cancer line and xenograft model, we found surprisingly diverse metastatic phenotypes, ranging from metastasis-incompetent to highly metastatic, which we then used to generate and test hypotheses regarding metastatic biology. We show that these distinctions in metastatic behavior were driven by specific differentially expressed genes, like IFI6, ID3, and KRT17, that can directly modulate invasion phenotype in multiple cancer contexts. Furthermore, we empirically demonstrate that these distinct metastatic phenotypes are pre-programmed prior to implantation, stably inherited over generations, and predictive of future metastatic behavior. (3) Metastatic dissemination was rapid, frequent, and complex in this model, transiting via different complicated seeding topologies, such as seeding cascades, parallel seeding, and more. Additionally, we illustrated that it is possible to capture subtle differences in tissue tropism and, using this strategy, we identify the mediastinum as a hub for metastatic seeding, perhaps because the mediastinal lymph is a favorable niche with extensive tissue connections (*77*). Extending beyond this xenograft model, these findings suggest that metastatic seeding patterns can be highly complex and possibly patient-specific.

As a first report, this work by necessity focuses on a single model of metastasis. Looking forward, it will be valuable to explore these findings in other experimental contexts, including those described below, wherein the lineage tracer may also be deployed. First, because A549s exhibit highly aggressive metastatic spread, the rapid and frequent metastatic events we observe may be most relevant to advanced stages of cancer progression. Future work could apply this lineage tracing approach to models of any stage in cancer progression, such as (i) other cell lines that represent earlier cancer stages, (ii) genetic models of inducible tumor initiation (*78*), or (iii) patient-derived xenograft (PDX) models (*79, 80*). Second, as is typical in xenograft experiments, this model requires immunodeficient host mice, and therefore does not reflect the pervasive influence wielded by the immune system on natural cancer progression (*81* – *83*); lineage tracing in syngeneic lines or autochthonous models of cancer could chart how an intact immune system influences cancer progression. Third, future work could build on our study by investigating the roles that other metastasis-associated gene candidates identified here possibly play in metastasis, or elucidate the molecular mechanism by which KRT17 diminishes metastatic phenotype *in vitro* and *in vivo*. Fourth, this work describes metastasis at the spatial resolution of tissues (i.e., not individual tumors) because we bulk tissues that potentially contain multiple tumors (including extensive micrometastases). An important direction would be to merge lineage data with high-resolution spatial information using the rapidly advancing techniques for spatial single-cell approaches (*24, 84* – *86*); this would clarify, among other features, the clonality of micrometastases, monophyletic versus polyphyletic dissemination (*12*), and the spatial constraints of tumor growth and metastasis.

More broadly, Cas9-enabled lineage tracing technologies should be readily deployable to explore aspects of cancer progression and evolution, especially the history and chronology of rare and transient events like metastasis. This will empower future work to more comprehensively describe cancer – as well as other biological phenomena – at unprecedented resolution and scale.

## Acknowledgements

We thank D. Yang, A. Khodaverdian, and R. Zhang for discussions; M. Jost and J.K. Nuñez for plasmids; and UCSF Center for Advanced Technology (E. Chow and S. Elmes) and UCSF Preclinical Therapeutics Core (B. Hann).

## Author contributions

All authors contributed to the design of experiments and analysis. J.J.Q. engineered cell lines, processed tissues, and prepared sequencing libraries. R.A.O. performed mouse surgeries and imaging. M.G.J. and J.J.Q. processed lineage tracing sequencing data. S.N. and J.J.Q. performed invasion assays. M.G.J. performed phylogenetic reconstruction and analyzed the trees and single-cell RNA-sequencing data. M.G.J. and N.Y. conceived and implemented the *FitchCount* algorithm. All authors aided in the interpretation of the analyses. J.J.Q., M.G.J., and J.S.W. wrote the manuscript, and all authors read and approved the final manuscript.

## Data and materials availability

The A549-LT cell line will be available via material transfer agreement. Raw sequencing reads and processed lineage tracing data files will be made available via GEO accession upon acceptance. Lineage tracer processing pipeline, phylogenetic reconstruction algorithm, and *FitchCount* are publicly available via the GitHub repository for Cassiopeia (github.com/YosefLab/Cassiopeia); all other analysis scripts and notebooks will be made publicly available upon acceptance on Github (github.com/mattjones315/MetastasisTracing).

## List of Supplementary Materials

Materials and Methods

Figs. S1-S24

Supplementary Text

sgRNA sequences for CRISPRi/a

Gene Lists

## Supplemental Materials

### Materials and Methods

#### Cell culture

A549 cells (human lung adenocarcinoma line, American Type Culture Collection CCL-185) and H1299 cells (ATCC CRL-5803) were maintained in Dulbecco’s modified eagle medium (DMEM, Gibco) supplemented with 10% (v/v) fetal bovine serum (FBS; VWR Life Science Seradigm), 2 mM glutamine, 100 units/mL penicillin, and 100 µg/mL streptomycin (hereafter “complete DMEM”). Cells were cultured at 37°C in a humidified 5% (v/v) CO_2_ atmosphere. Cells were split into fresh culture medium every two to three days by trypsinization with TrypLE reagent (Gibco) quenched with complete DMEM, and maintained at cell density as recommended by ATCC.

#### Plasmid design and cloning

The triple-sgRNA-BFP-Puromycin^R^ lentivector, PCT62 (to be made available on Addgene), was constructed using four-way Gibson assembly (NEB) as previously described (*88*), and expresses three sgRNA cassettes driven by distinct U6 promoters, with constitutive BFP and puromycin-resistance markers for selection. The three sgRNAs are complementary to the three cut-sites in the Target Site (PCT48), except for precise single base-pair mismatches that decrease their avidity for the cognate cut-sites and subsequently slow lineage recording kinetics (*89*). Guide RNAs for CRISPRi and CRISPRa experiments were chosen from the human CRISPRi/a v.2 libraries (Addgene #83969 and #83978, respectively) and were cloned into the pLG1 lentiviral backbone (Addgene #84832) using the annealing and ligation as previously described (*90*).

#### Lentivirus preparation and infection method

Lentivirus was produced by transfecting HEK293T cells with standard 4th generation packaging vectors delivered by TransIT-LTI transfection reagent (Mirus) as described in (*34*) and (*91*). Target Site (PCT48) lentiviral supernatant was concentrated 10-fold using Lenti-X Concentrator (Takara Bio) according to manufacturer’s instructions. Viral preparations were filtered and frozen prior to infection. Triple-sgRNA lentiviral preparation (PCT62) was titered and diluted to a concentration to yield approximately 50% infection rate. All infections were performed in 6-well plates with 2 mL media and 2×10^5^ adhered cells per well; 1 mL of titered lentivirus was added to the culture medium with 8 µg/mL of polybrene and incubated at 37°C overnight, after which media was replaced and cells were expanded to larger culture volumes as needed.

#### Cell line engineering and selection strategies

Two separate lineage tracing-competent A549 cell lines (hereafter, “A549-LT1” and “A549-LT2”) were engineered separately and by different methods, primarily distinguished by (i) the method of high copy-number transduction of the Target Site (piggyBac transposition and high-titer lentiviral infection, respectively) and (ii) the version of the “molecular recorder” technology used (the original components described in (*23*) and the improved components described in (*34*), respectively). For both A549-LT1 and -LT2, the cells were first infected with Luciferase-Neomycin^R^ lentivirus; two days following infection, *neomycin(+)* cells were selected by treatment with 800 µg/mL G418 Geneticin (Thermo Fisher) every second day for 10 days, expanding the cells as necessary. Next, the A549+Luciferase cells were infected with either Cas9-P2A-BFP or Cas9-P2A-mCherry lentivirus, respectively; three days following infection, *BFP(+)* or *mCherry(+)* cells were collected by fluorescence-activated cell sorting (FACS) using the BD FACS Aria II at the UCSF Center for Advanced Technology core. Next, the A549+Luciferase+Cas9 cells were serially transduced with different versions of the GFP-TargetSite vector (transposon-based PCT17 or lentivector-based PCT48, respectively). For the A549-LT1 line, the transposon transduction was performed by electroporation using an Amaxa Nucleofector (Lonza) with 100 ng of PiggyBac transposase plasmid (SBI System Biosciences) and 500 ng of PCT17 transposon plasmid. For the A549-LT2 line, the high-titer PCT48 lentiviral infections were performed as above, but in triplicate and pooled to further maximize diversity. Following GFP-TargetSite transduction, *GFP(+)* cells were collected by FACS. These steps were repeated two additional times for a total of three serial transductions of the Target Site, fluorescence-sorting cells with progressively higher GFP fluorescence after each transduction (**Fig. S2**). Finally, four days prior to implantation, A549+Luciferase+Cas9+TargetSite cells were transduced by titered triple-sgRNA (with BFP-Puromycin^R^ markers; PCT61) lentivirus; 36 hours following infection, *BFP(+)* cells were collected by FACS and returned to *in vitro* culture until final preparation for implantation, thus producing lineage tracing-competent A549-LT1 and -LT2 cell lines.

#### Mouse care

Mouse experiments were performed as in (*38*). Six-to eight-week-old female SCID C.B-17 mice (C.B-*Igh-1*^*b*^/IcrTac-*Prkdc*^*scid*^; Taconic) were maintained in specific-pathogen-free conditions in facilities approved by the American Association for Accreditation of Laboratory Animal Care. Surgical procedures were reviewed and approved by the UCSF Institutional Animal Care and Use Committee (IACUC), Protocol #AN107889-03C.

#### Orthotopic lung xenografts

To prepare A549-LT cell suspensions for implantation, cells were collected from culture by trypsinization and quenched with complete DMEM. Cells were then washed in cold PBS and resuspended with cold Matrigel matrix (BD Bioscience) at the appropriate final concentration (500, 1000, 3000, and 10,000 cells/µL for mice M5k, M10k, M30k, and M100k, respectively). The Matrigel cell suspensions were gently mixed and transferred into a 1-mL syringe and remained on ice until implantation. Orthotopic implantations were performed as in (*38*): mice were placed in the right lateral decubitus position and anesthetized with 2.5% inhaled isoflurane. A 1-cm surgical incision was made along the posterior medial line of the left thorax. Fascia and adipose tissue layers were dissected and retracted to expose the lateral ribs, the intercostal space, and the left lung parenchyma. Upon recognition of left lung respiratory variation, a 30-gauge hypodermic needle was used to advance through the intercostal space approximately 3 mm into the lung tissue. Care was taken to inject 10 µL (5,000, 10,000, 30,000, or 100,000 cells, respectively) of cell suspension directly into the left lung. Mice were observed post-procedure for 1–2 hours, and their body weights and wound healing were monitored weekly. Experiments were performed across two mouse cohorts: A549-LT1 cells were implanted in the first mouse cohort (including mice M10k and M100k); A549-LT2 cells were implanted in the second cohort (including mice M5k and M30k).

#### Bioluminescence imaging

Mice were imaged with the Xenogen IVIS 100 bioluminescent imaging system at the UCSF Preclinical Therapeutics Core. Bioluminescence monitoring of the tumor engraftment and metastatic progression was performed biweekly. Before imaging, mice were anesthetized with inhaled isoflurane and injected intraperitoneally with 200 µL of *D*-Luciferin at a dose of 150 mg/kg body weight. For each mouse, bioluminescent signal was measured from the thoracic cavity in the supine position and calculated automatically using Living Image Software; radiance units are photons/s/cm^2^/steradian. Mice were sacrificed after bioluminescent signal was anatomically extensive throughout the thorax but before the mice exhibited labored breathing (53 to 80 days post-implantation; varied by mouse). Mice were injected with *D*-Luciferin as before and subsequently killed; the heart and lungs were resected *en bloc*, the heart was removed, and the right and left lung lobes were separated. Tumorous tissues were imaged *ex vivo* by either bioluminescence (performed as before) or fluorescence using a stereo fluorescent microscope (Nikon).

#### CRISPRi/a perturbations and invasion assays

CRISPRi/a cell lines were first generated by infecting parental cell lines with lentivirus carrying either the CRISPRi (dCas9-BFP-KRAB; Addgene #46911) or CRISPRa (dCas9-XTEN-VPR-GFP (*92*)) genetic component, respectively; stably transduced cells were selected to purity by FACS. CRISPRi and CRISPRa activity was confirmed by perturbing the gene expression of the cell-surface marker genes CD81 and CD151 (for CRISPRi) and CXCR4 (for CRISPRa); perturbations were performed by infecting with lentivirus carrying sgRNA against the target marker genes (see sgRNA spacer sequences in the Supplemental Materials) and transduced cells were selected for 4 days with puromycin as described above. Seven days after transduction, the perturbed cells were collected and stained with APC-labelled antibodies against CD81, CD151, and CXCR4 (BioLegend), and the fluorescence of >10^3^ individual cells was measured by flow cytometry (Attune NxT, Thermo Fisher Scientific). For invasion assays, CRISPRi and CRISPRa cell lines were treated with lentivirus carrying sgRNAs against the targeted metastasis-associated gene candidates (see sgRNA spacer sequences in the Supplemental Materials). We selected five candidate genes of interest to assay based on the following criteria: the strongest positive (IFI27) or negative (KRT17) genes identified from mouse M5k; in the same gene family as IFI27 (interferon-induced; IFI6); or previous evidence of modulating invasion phenotype in similar models (ASS1, ID3) (*59, 60*). Cells transduced with sgRNAs against these genes were selected and cultured for one week, as above. For the invasion assays, DMEM supplemented with 10% FBS was added to the bottom chamber of a 24-well trans-well plate. The perturbed cells were counted, and 1.5×10^4^ cells were resuspended in serum-free media and added to the top chamber of 8-µm pore matrigel-coated (invasion) or non-coated (migration) trans-well inserts (Corning BioCoat). After 16 hours, non-invading cells on the apical side of inserts were scraped off and the trans-well membrane was fixed in methanol for 15 minutes and stained with Crystal Violet for 30 minutes. The basolateral surface of the membrane was visualized with a Zeiss Axioplan II immunofluorescence microscope at 10×. Each trans-well insert was counted manually at 10× and performed in triplicate. Invasion phenotype was calculated as the number of invading cells through the matrigel membrane divided by the mean number of cells migrating through control insert. Differential invasiveness was assessed using a two-tailed *t-*test comparing the invasion rate of the perturbation to both negative controls.

#### Tissue homogenization and single-cell preparation

Collected tumorous tissues were minced on ice and placed in separate gentleMACS C Tubes (Miltenyi Biotec) with 5 mL each of tissue dissociation buffer (30 mL DMEM/F-12 culture (Gibco) supplemented with 16 mg collagenase (Worthington Biochemical), 5 mg Liberase TL (Roche), and 100 U DNase I (Worthington Biochemical)). Tissues were homogenized on a gentleMACS Octo Dissociator (Miltenyi Biotec) for one minute, then incubated in a 37°C water bath for five minutes followed by repeated trituration using a 1000 µL pipette tip; incubation and trituration were repeated until tissues were completely dissociated, three to six times in total. Cells were pelleted by centrifugation at 500 rcf for 5 minutes at 4°C. To remove red blood cells, the cell pellets were resuspended in 1 mL ACK Lysing Buffer (Gibco) and incubated at room temperature for 3 minutes, followed by centrifugation. To achieve a single-cell resuspension, the cell pellet was resuspended in 500 µL TrypLE reagent and incubated at 37°C for 5 minutes followed by trituration; digestion was quenched with 1 mL complete DMEM medium, centrifuged, and resuspended in 1 mL of cold complete DMEM. Cells were strained through a 40 µm nylon filter-cap tube (Corning). Cancer cells were fluorescence-sorted (identified by high GFP and BFP fluorescence) at 4°C to remove mouse cells and debris, pelleted, resuspended in 1 mL cold phosphate-buffered saline with 0.04% bovine serum albumin, and filtered through a 40 µm FlowMi filter-tip (Bel-Art). The concentration of cells was determined on an Accuri flow cytometer (BD Biosciences), then pelleted. Finally, the cells were thoroughly resuspended in 20 µL phosphate-buffered saline with 0.04% bovine serum albumin, mixed with the Chromium Single Cell 3′ V2 Kit reagents (10X Genomics) according to the User Guide, and then loaded into droplet emulsions using the Chromium Controller, thus facilitating single-cell transcriptional measurements. Separate Chromium lanes were used for each tissue sample.

#### Sequencing library preparations

Gene expression libraries were prepared according to the Chromium Single Cell 3′ V2 Kit User Guide. To prepare the Target Site libraries, the amplified cDNA libraries were further amplified with Target Site-specific primers containing Illumina-compatible adapters and sample indices (forward: 5′-CAAGCAGAAGACGGCATACGAGATNNNNNNNNGTCTCGTGGGCTCGGAGATGTGTATAAGAGACAGAATCCAGCTAGCTGTGCAGC; reverse: 5′-AATGATACGGCGACCACCGAGATCTACACNNNNNNNNTCTTTCCCTACACGACGCTCTTCCGAT CT; “N” denotes sample indices) using Kapa HiFi HotStart ReadyMix (Roche), as described in (*34*). Approximately 24 fmol of template cDNA was used per sample, divided between four identical reactions to avoid possible PCR-induced library biases. PCR products were purified and size-selected using SPRI magnetic beads (Beckman) and quantified by BioAnalyzer (Agilent).

#### Library sequencing

Sequencing libraries from each tissue sample were pooled to yield approximately equal coverage per cell per sample; gene expression libraries and Target Site amplicon libraries were pooled in an approximately 10:1 molar ratio to yield more RNA reads than Target Site reads. Libraries were further pooled with approximately 5% PhiX genomic DNA library added for quality-control. The libraries were sequenced using a custom sequencing strategy on the NovaSeq S2 platform (Illumina) in order to read the full-length Target Site amplicons. Sample identities were read as dual indices (I1 and I2: 8 cycles each); the 10X cell barcode and unique molecular identifier (UMI) sequences were read first (R1: 26 cycles) and the Target Site sequence was read second (R2: >250 cycles). Over 4.70 billion sequencing clusters passed Illumina QC filters and were processed as described below. All raw and processed data will be made available on GEO (accession pending). Only the first 98 bases per read were used for analysis in the RNA expression libraries to mask the longer reads required to sequence the Target Sites.

#### TargetSite sequencing data processing

Raw Target Site sequencing data was processed using the Cassiopeia processing pipeline as defined (*34*). Briefly, reads with identical cellBC and UMI were collapsed into a single, error-corrected consensus sequence representing a single expressed transcript. Consensus sequences with poor quality or sequencing coverage were removed, according to pipeline-defined thresholds. Each consensus sequence was aligned to the wild-type reference Target Site sequence, and the intBC and indel alleles were called from the alignment. These data are summarized in a molecule table which records the cellBC, UMI, intBC, indel allele, read depth, and other relevant information.

#### Calling clonal populations

Collected cells were grouped into “clonal populations”, defined as populations of cells which descended from a single engineered clone and which therefore shared the same set of intBCs. This was accomplished using an iterative strategy of defining *de novo* sets of intBCs and assigning cells to clonal populations based on the sets, and repeating this process to further refine the assignments, similar to the strategy described in (*34*), as follows: (1) The most frequently observed intBC (“top intBC”) was identified from the molecule table. (2) A “clustered intBC set” was defined as the intBCs present in >20% of cells containing the top intBC. (3) Cells with >25% of their Target Site UMIs pertaining to the clustered intBC set were then collected into a “cluster” and removed from the molecule table. (4) This process of identifying the top intBC, clustered intBC set, and cell clusters was iterated until at least one of two stopping criteria was met: (i) the returned cluster of cells was empty, indicating that the clustering strategy had exhausted well defined clusters, or (ii) the total fraction of clustered cells exceeded 98% of all cells in the molecule table. (5) Next, a “clonal population intBC set” was defined for each cell cluster, defined as intBCs that were present in >20% of cells in the cluster. Some clusters were manually redefined based on overlapping clonal population intBC sets likely resulting from the serial transduction strategy of cell line engineering; i.e., some clusters were expected to share intBCs because they were clonally related during serial integration of the Target Site (see **Fig. S20B** for an example). (6) Finally, cells from the initial molecule table were assigned to a clonal population if >60% of its UMIs pertained to a single clonal population intBC set. Clonal populations with fewer than 25 cells were not considered, as lineage tracing over small cell numbers is less informative.

In the pre-implantation sample, we assigned cells to clonal populations observed in M5k by evaluating the proportion of TargetSite UMIs that corresponded to a clonal population intBC set. As above, we summed together the UMI proportions of each clonal population intBC set for each cell, thus yielding a similarity score that we could use to assign cells to clones. Here, we assigned a cell to a clone if at least 75% of its UMIs pertained to that clonal population intBC set.

#### Filtering cells and assembly of allele tables

Cells with poor Target Site capture (defined as cellBCs with <10 Target Site UMIs) were removed due to poor representation. Cells with ambiguous allele states were removed if >10% of the UMIs had conflicting allele states (i.e., intBC-distinguished Target Sites with more than one allele state), likely resulting from PCR errors or cell doublets (wherein both cells are in the same clonal population; so-called “intra-doublets”). Cells with ambiguous clonal population assignment were removed if <60% of its UMIs pertained to a single clonal population’s set of intBCs; this likely results from cell-free Target Site transcripts, PCR errors, or cell doublets (wherein cells are from different clonal populations; so-called “inter-doublets”). Finally, the filtered molecule table with clonal population assignments is collapsed into an allele table, which summarizes the cellBC, intBC, indel allele, number of UMIs, assigned clonal population, and other relevant information.

#### Calculation of Tissue Dispersal Score

The Tissue Dispersal Score is the inverted Cramér’s *V* (1 - *V*), a statistical measure of the association between two variables derived from the chi-squared test, ranging from 0 (no deviation from the background) to 1 (complete deviation from the background). For a given clonal population, we first perform a chi-squared test by forming a contingency table *X* over summarizing the number of cells found in each tissue for the clonal population, and the number of cells found in each tissue aggregated across all other clonal populations (referred to as the “background”). Importantly, the number of cells found in each tissue in the background are scaled such that the sums for both columns in *X* are equal. After performing a chi-squared test on the *r* × *k* contingency table, *X*, with a total of *N* counts across both columns, we derive the bias-corrected Cramér’s *V* test (*93*) statistic:

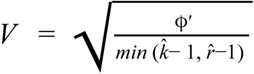

where 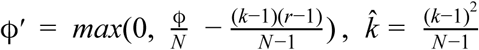, and 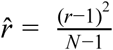. Finally, this statistic is inverted to obtain the Tissue Dispersal Score.

#### Filtering of clonal populations for tree reconstruction

We removed a minority of clonal populations that exhibited suboptimal lineage tracing parameters, as defined by <15% of cut-sites bearing indels (i.e., an estimate of the lineage recording kinetics) and <66.7% of unique cell allele states (i.e., an estimate of the lineage diversity and information content). Phylogenetic reconstruction in such poor parameter regimes resulted in low information trees (indicated by the shallow tree depth of filtered trees in **Figs. S6E** and **S7C**) that would not be interpretable for sensitive downstream analysis, such as calculating the inferred rate of metastasis.

#### Assembly of character matrices for phylogenetic tree reconstruction

To reconstruct lineages, we created “character matrices” from the allele tables of each clonal population using the Cassiopeia software. Specifically, we summarized the indels observed in each of the *N* cells in a clonal population across the *M* cut-sites to form an *N*x*M* matrix where each entry is an integer-representation of the indel observed in that cell at that cut-site. These entries are referred to as “character-states” or “states”; the columns of this matrix are referred to as “characters”; and the rows are referred to as “cells” or “samples”, interchangeably. Missing data was specified as the string ‘-’, and uncut sites were specified as ‘0’.

#### Cassiopeia-build pipeline overview

We reconstructed each clonal population’s phylogeny using the Cassiopeia-Hybrid module, as described in (*34*). Briefly, Cassiopeia-Hybrid takes as input the *N*x*M* character matrix and first uses the Cassiopeia-Greedy heuristic to split cells into small groups based on the presence, or absence, of states that occurred early in the phylogeny (referred to as “character-splits”); then, each subproblem is solved precisely with the Steiner-Tree-based Cassiopeia-ILP module; finally, the completed subproblems are merged together to form the final tree. All clonal populations were reconstructed with a maximum neighborhood size of 10,000 (--max_neighborhood_size 10000) and a maximum time to convergence of 3.5 hr (--time_limit 12600).

While missing data is handled in Cassiopeia-ILP by considering all possible assignments, Cassiopeia-Greedy handled missing data with two heuristics: first, cells with missing data in a character-split were classified based on their 10 closest neighbors by a modified Hamming distance normalized by the number of overlapping characters two cells shared (using the “knn” greedy missing data mode in Cassiopeia: --greedy_missing_data_mode knn --num_neighbors 10); second, characters with greater than 30% missing data were not selected as character-splits (i.e., --greedy_max_missing_rep 0.3).

We determined prior-probabilities for each indel by using the proportion of times it appeared, independently, on an intBC in a clonal population (accounting for all clonal populations in our data). These priors were used in two ways: first, they were used to select indels as character-splits that likely occurred early in the phylogeny (i.e. they appeared in several cells, but had a relatively low prior) for Cassiopeia-Greedy; second, we used the priors to weight edges in the Steiner-Tree optimization within Cassiopeia-ILP (invoked with the --weighted_ilp argument to Cassiopeia’s command line interface). Priors were provided to Cassiopeia’s reconstruction algorithm using the --mutation_map argument.

To transition from Cassiopeia-Greedy to -ILP, we introduced a modified criteria based on the maximum distance from a cell population’s Latest Common Ancestor (LCA) to any given cell in the population. Specifically, a new Cassiopeia-ILP subproblem is spawned if the distance to an LCA of a group of cells is below a user-defined threshold (--cutoff <lca_cutoff>). An appropriate LCA distance threshold was determined, iteratively, such that no Cassiopeia-ILP subproblem exceeded the user-defined maximum neighborhood size. The LCA transitioning criteria was invoked with --hybrid_lca_mode.

#### Neighbor-Joining reconstructions

For neighbor-joining reconstructions, we used the --neighbor_joining_weighted reconstruction option in the Cassiopeia package, which uses a scikit-bio implementation of neighbor-joining (version 0.5.5). To define distances between cells as input, we utilized a version of the modified Hamming distance function that incorporated prior-probabilities (prior-probabilities are passed in via the --mutation_map argument). Specifically, we defined *d*′(*a, b*) as the sum of all the pairwise *h*′_*p*_(*a*_*i*_, *b*_*i*_) values across the cut-sites in a clonal population:

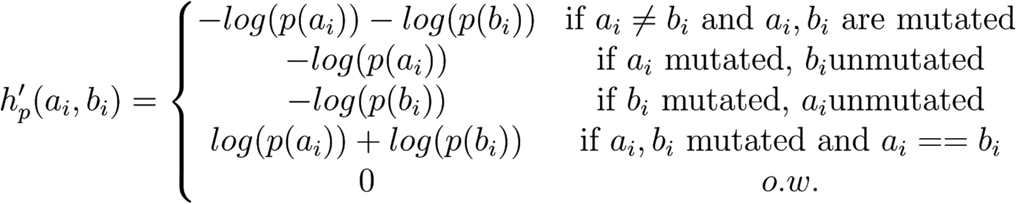

Finally, distances *d*′(*a, b*) were normalized by the number of cut-sites overlapping between cells *a* and *b*.

#### Allelic vs. phylogenetic distance calculations

The concordance between allelic and phylogenetic distances were used to assess the agreement between a tree’s phylogenetic structure and the observed mutations. To quantify allelic distances, we defined a modified Hamming distance between cells *a* and *b*,

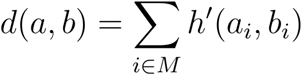

Where *M* is the number of cut-sites in the clonal population that cells *a* and *b* come from, *a*_*i*_ and *b*_*i*_ are the states observed at the *i*^*th*^ character, and *h’* is defined as follows:

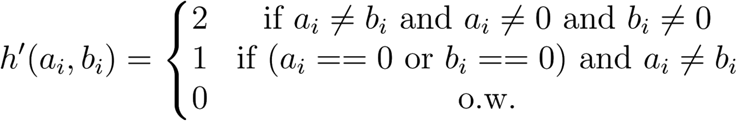

Importantly, these values were normalized by 2 * *M*, the maximum distance for a pair of cells.

The phylogenetic distance was calculated for all pairs of cells as the number of non-zero length edges that separated the two cells in the tree. The phylogenetic distances were normalized by the “diameter” of the phylogeny, i.e., the maximum distance between any pair of cells.

#### Derivation of the AlleleMetRate

The AlleleMetRate is provided as a “tree-agnostic” measurement of a clonal population’s intrinsic metastatic potential. Intuitively, the rate measures the proportion of cells that do not reside in the same tissue as their closest relative (as determined by the modified Hamming distance between two cells’ character-states). Importantly, if a cell has more than one closest relative, each of their votes are normalized by the number of relatives this cell has. More formally, the contribution a cell has to the AlleleMetRate is

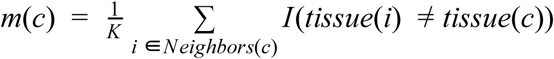

Where *K* is the number of closest relatives a cell has, and *I(*)* is an indicator function that equals 1 if the tissue of cell *i* is different from the tissue of cell *c*. The AlleleMetRate reported is

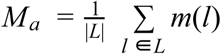

Where *L* is the set of all leaves in the tree.

#### Derivation of the TreeMetRate

The TreeMetRate is derived from the Fitch-Hartigan maximum parsimony algorithm (Fitch, 1971; Hartigan, 1973). Briefly, the Fitch-Hartigan algorithm takes in a rooted tree with labelings at the leaves (in our case labels indicate which tissue the cell was obtained from) and in our case reports two items: (a) a distribution of labels at the internal nodes that minimize the number of transitions between tissues (i.e., achieves maximum parsimony); and (b) the minimum number of transitions between tissues (referred to as the “parsimony score”). The Fitch-Hartigan algorithm is an efficient (scaling with *n* · *k* where *n* is the number of cells and *k* is the number of tissues) and common algorithm for solving the “Small Parsimony Problem”, as opposed to the “Large Parsimony Problem” which attempts to find the *tree* of maximum parsimony.

The Fitch-Hartigan algorithm operates in two phases: first, by propagating up from the leaves the set of possible labelings at each internal node; second, by traversing in depth-first order from the root of the phylogeny and selecting one state for each internal node from its set of optimal labels. Conveniently, the parsimony score can be obtained from this second phase. To finally report the TreeMetRate, we simply normalize this parsimony score by the number of edges in the tree.

#### Derivation of the single-cell MetRate

The scMetRate for a given cell is defined as the average of all TreeMetRates for the clades that contain that cell. Specifically, we employ the following algorithm:

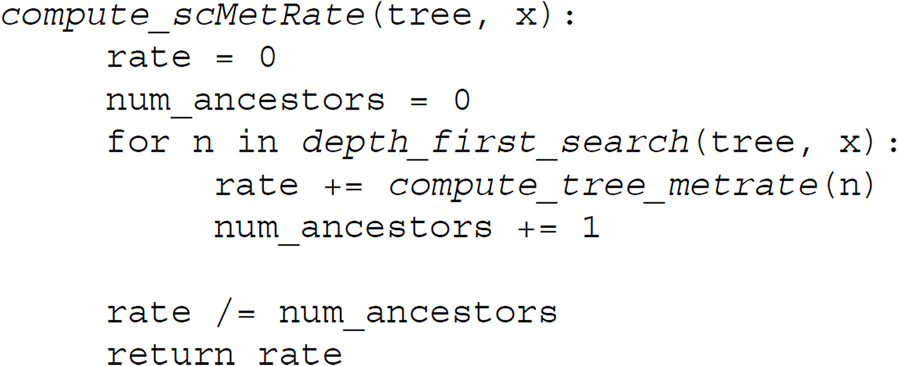

Where *compute_tree_metrate(n)* is a function that computes the TreeMetRate for the subtree rooted at an internal node *n* (as described above).

#### scRNA-seq preprocessing and normalization

RNA-seq libraries were quantified using CellRanger version 2.1.1 with the GRCh38 genome build for M5k and M30k; and CellRanger version 2.0.0 with the hg19 genome build for M10k and M100k. Cells not found in the Target Site Library were filtered out. All RNA-seq datasets were normalized identically – we first normalized the UMI counts in every cell to the median number of UMIs found in each library (referred to hereafter as “UMI-normalized” counts) and then each matrix of UMI-normalized counts was log-transformed (after adding a 1-pseudocount to preserve 0’s in the data; hereafter referred to as “log-transformed” counts). In the analysis presented in Figure 5A we applied scVI (version 0.6.5) to the raw counts of the M5k and the pre-implantation cells to obtain a shared, batch-corrected latent space (learning rate of 1e-3, 400 epochs, 10 latent dimensions). This space was used for downstream projections and *Vision* analyses.

#### Vision analysis on M5k

*Vision* (version 2.1) was applied to the UMI-normalized counts for the 35,006 filtered and clonal population-assigned cells that overlapped between M5k’s scRNA-seq and TargetSite libraries. Genes were filtered out using the Fano-filter procedure in *Vision*, leaving 4,005 genes. To provide a latent space to *Vision*, we computed a reduced dimension embedding using scVI (*94*) on the filtered gene matrix of raw counts, using a learning rate of 1e-3 and 40 epochs to produce 10 latent components. No batch correction was used. Signatures were obtained from MSigDB (*95*).

#### Differential expression of left lung samples

We used the UMI-normalized counts from M5k to perform differential expression between cells in the left lung, stratified by whether or not they were from a clonal population that metastasized from the left lung at any point in the experiment. All cells found in the left lung that were from clonal populations that metastasized are labeled as “Metastatic”. We performed 5 differential expression tests for the cells in each group: CP029 vs. “Metastatic”; CP036 vs. “Metastatic”; CP078 vs. “Metastatic”; CP094 vs. “Metastatic”; and “Metastatic” vs. all cells in CP029, CP036, CP078, and CP094. Tests were performed using the Wilcoxon rank sums test as implemented in Scanpy (*96*).

#### Differential expression of single-cell MetRate by Poisson regression

Genes differentially expressed between highly and lowly metastatic cells were determined by using a Poisson regression scheme (as implemented in the GLM package in *Julia*, version 1.3.7). Cells were first segmented into “Low” and “High” groups (referred to as *m*) based on the scMetRate. Then, the following model was employed on the UMI-normalized counts:

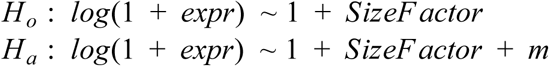

Where *SizeFactor* was defined as the number of genes detected in that cell and *expr* refers to the UMI-normalized expression count of a particular gene. A Likelihood Ratio Test (LRT) was used to determine the significance of the alternative hypothesis (in particular, the significance of the model fit improvement due to the variable *m*). Log2 fold-changes (Log2FC’s) were calculated by comparing the mean expression of a gene in the “High” group to that of the “Low” group of the UMI-normalized counts with a pseudocount of 0.01 added to account for zero counts. P-values were adjusted using the Benjamini-Hochberg false discovery rate procedure. Genes were considered significant if their FDR-adjusted p-value was less than 0.01.

Cells with greater than 20% counts attributed to the mitochondrial genome were filtered out and genes observed in fewer than 10% of cells were filtered. We proceeded with 35,005 cells and 4,993 genes for M5k; 1,492 cells and 11,171 genes for M10k; 17,175 cells and 7,620 genes for M30k; and 2,105 cells and 10,371 genes for M100k. Analysis was performed similarly for the M10k, M30k, and M100k mice, with considerations of their underlying scMetRate distributions taken into account. To ensure an accurate reflection of High and Low metastatic groups, cells were stratified according to the median in M10k, the 25th percentile in M100k; and in M30k because the scMetRate was not bimodal, we stratified cells into groups below the 25th and above the 75th percentiles.

Positive and negative gene “hits” in the M5k analysis were determined using a “discriminant” score defined as the absolute value of log2(fold-change) times the negative log10(FDR) of the gene. Genes with a discriminant score greater than 600 were annotated as “hits”. For the analysis in Fig. 4C,D we identified significant gene sets as those with an FDR-corrected p-value < 0.01 and log2FC > 0 or log2FC < 0 for positive and negative sets, respectively. Significance of gene set overlap was assessed with the R package *SuperExactTest (87)*, version 1.0.7) Finally, genes were considered “reproducible” in the M10k, M30k, and M100k analyses if they were found to be significant (FDR < 0.01) and their effect was in the same direction as in M5k.

A consensus “Metastatic Signature” was formed from the top genes in the positive and negative direction from each mouse. Specifically, we first ranked each list by the discriminant score described above. Then, for each direction in each mouse as measured by log2FC, we selected the top 30 genes. We then took the unique genes in each direction across all mice and formed a directional signature from these two lists. Scores for this signature were obtained for each cell with *Vision*.

#### Identifying gene signatures correlated with TreeMetRates

Gene signature scores for single cells were calculated with *Vision*, and pseudo-bulked by clonal population by taking the mean signature score for each signature across the cells in a clonal population. We then calculated the Spearman correlation using the scipy Python package (version 1.2.2) between the pseudo-bulked gene signature score and the TreeMetRate for a given clonal population.

#### Hotspot *analysis for CP007*

*Hotspot* (version 0.9.0) was acquired from Github (https://github.com/YosefLab/Hotspot) and run on the UMI-normalized counts for the 603 cells in CP007 and all genes expressed in at least 10% of cells.We used the UMI-adjusted negative binomial model (“danb”) as the background gene expression model and distances between cells were computed from the CP007 tree reconstructed with Cassiopeia. We used 40 neighbors for the *Hotspot* analysis. Modules of at least 120 genes were identified from all genes with an FDR < 0.1, and the *Hotspot* module scores were used to annotate the tree. The right lung (E) only (“RE-only”) control was performed identically, except for first removing cells that were not from the RE tissue sample in CP007 (437 cells).

#### *Inference of Tissue Transition Matrices with* FitchCount

To infer the relative propensities of a clonal population’s cells to transition between any two tissues, we developed an efficient algorithm for aggregating together optimal solutions proposed by the Fitch-Hartigan algorithm (see **Supplementary Text** for algorithm and proof). Briefly, *FitchCount* is a dynamic programming algorithm that operates on a rooted phylogeny. It begins by performing the “bottom-up” phase of the Fitch-Hartigan algorithm to obtain a distribution of optimal labels over each internal node and then employs a computationally efficient algorithm for computing the number of times a given transition occurs in all optimal solutions proposed by the Fitch-Hartigan algorithm. Finally, this algorithm returns a square matrix, *M*, where each value *m*_*i,j*_ indicates the number of times a transition from tissue *i* to tissue *j* was observed in the tree over all optimal solutions.

In **Fig. 6** and **Fig. S22**, we report transition matrices for the probability of a cell metastasizing from one tissue to another, given that the cell metastasizes; i.e., *P* (*m*_*i,j*_ | *i* ! = *j*). To obtain these conditional probability tables, *P*, we first set *diag(M)* to 0 (indicating that the probability of self-transition is 0) and re-normalize each row to sum to 1.

#### Feature selection and Principal Component Analysis (PCA) of tissue transition matrices

To identify the trends in the tissue transition matrices presented in **Fig. 6**, we performed dimensionality reduction using Principal Component Analysis (PCA) on the flattened tissue transition matrices. Beyond all the conditional probability of transitioning between tissue samples summarized in the tissue transition matrix *M*, we included additional features which we hypothesized would aggregate important signals. In particular, the following were added by deriving statistics from *P*, the conditional probability matrix, and *M*, the unnormalized tissue transition matrix (recall that we observed 6 tissue samples in M5k: left lung [LL], right lung W [RW], right lung E [RE], mediastinum 1 & 2 [M1 & M2] and liver [Liv]):

- The Left Lung reseeding rate (defined as the sum of the probabilities in the LL-column of *P)*
- The Mediastinum tropism rate (defined as the sum of the probabilities in the M1 or M2 columns of *P*)
- The Liver tropism rate (defined as the sum of the probabilities in the Liver column of *P*)
- The Right Lung tropism rate (defined as the sum of the probabilities in the Right Lung column of *P*)
- The Left lung to Right Lung seeding rate (defined as the sum of the probabilities *p*_*LL, RW*_ and *p*_*LL, RE*_)
- The Right Lung to Left Lung reseeding rate (defined as the sum of the probabilities *p*_*RW, LL*_ and *p*_*RW, LL*_)
- The “intra lung” metastatic rate (defined as the sum of the probabilities leading from the LL to either RL sample and vice versa in *P*)
- The “intra mediastinum” metastatic rate (defined as the sum of probabilities going between *M1* and *M2* samples in *P*)
- The Primary Seeding density (defined as the density of transitions observed in the LL row of *T*).
- The M1 seeding density (defined as the density of transitions in the M1 row of *T*).
- The M2 seeding density (defined as the density of transitions in the M2 row of *T*).
- The Mediastinum seeding density (defined as the sum of densities of transitions in the M1 and M2 rows of *T*).

This resulted in a set of 48 features used for dimensionality reduction. To perform PCA on this matrix, we concatenated the 48 features for each clonal population, standardized the features, and used PCA as implemented in the scikit-learn Python package (version 0.21.3).

#### Classification of Seeding Topologies

To detect seeding topologies, as presented in **Fig. 5J** and **K**, we devised simple algorithms from the tree structure and the conditional probability tissue transition matrices, *P*.

- To identify clonal populations that exhibited primary seeding, we evaluated if there existed some *p*_*LL, x*_ > 0 for some *x* ∈ {*RW, RE, M* 1, *M* 2, *Liv*}.
- To identify clonal populations that experienced reseeding to the left lung, we evaluated if there existed some *p*_*x, LL*_ > 0 for some *x* ∈ {*RW, RE, M* 1, *M* 2, *Liv*}.
- To identify clonal populations that exhibited bidirectional seeding (i.e. seeding to any tissue that previously served as a source for a metastatic event), we evaluate whether or not there exist two metastatic events, the second of which returns to the source of the first metastatic event. To do so, we sampled 100 solutions from the Fitch-Hartigan top-down procedure (which assigns labels to internal nodes) and evaluated whether or not we observe a path from the root to any leaf that follows a bidirectional seeding pattern (not necessarily on consecutive edges).
- To identify clonal populations that exhibited a seeding cascade, we evaluated whether or not there existed any tissue transition in a clonal population from *s*_*i*_ → *s*_*j*_ such that *s*_*i*_ ≠ *s*_*j*_ ≠ *LL*. Even if we do not observe any cells in the left lung of a given tumor, we know that the pattern previously described is evidence of a seeding cascade because all tumors began in the left lung of the mouse.
- To identify clonal population that exhibited parallel seeding, we evaluated whether or not there existed two conditional probabilities in *P*, 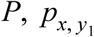 and 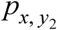, that were non-zero from any tissue *x* to any pair of tissues *y*_1_ and *y*_2_ such that *y*_1_ ≠ *y*_2_.

#### Model framework, parameters, and assumptions of metastasis simulator

To simulate metastatic events, we built on the simulation framework present in Cassiopeia as previously described in (*34*). The framework simulates a series of *D* binary splits over cells with *M* characters & *S* states per character; mutations are introduced every generation for each cell according to a per-character mutation rate, *m*, and dropout is simulated at the end according to the dropout rate, *d*.

To simulate metastatic processes, we introduced three new parameters: a cell-division rate (α), a metastatic rate (μ), and a probability map of transitioning between tissues should a metastatic event occur (*P)* that follows the same structure as the conditional tissue transition matrices discussed above. We assume that every generation, a cell first has the opportunity to divide (by evaluating whether *Unif* (0, 1) < α) and then if so, each of its offspring has the opportunity to metastasize (by evaluating *Unif* (1) < μ). If the daughter cell metastasizes, a tissue is chosen randomly according to the probabilities specified in *P*. To note, in this new simulation framework, we choose to control the size of a clonal population randomly with α, instead of subsampling at a predetermined rate as described previously.

Importantly, we can also apply this simulation framework on top of a tree by simply traversing the edges of the tree and simulating metastasis without cell divisions. For the results presented in this study, we used the M5k mouse to parameterize the simulations - especially in regards to the number of tissues that cells can metastasize to (6) and the distribution of metastatic rates (between 0 and 0.3).

#### Assessing accuracy of the TreeMetRate

We used the simulation framework to evaluate how well the Tissue Dispersion Score, AlleleMetRate, and TreeMetRate measurements were able to capture the underlying metastatic rate μ of a simulation. To do, we simulated trees of variable depth *D* and parameterized with some doubling rate, α ∼ *U nif* (0, 0.7), and metastatic rate, μ ∼ *U nif* (0, 0.3). We simulated 1000 such trees, with *D* ∈ {10, 14} (90% of trees were simulated with *D = 10*, and 10% of trees were simulated with *D = 14*, to roughly reflect the distribution of trees we see in our data). Target Sites were simulated using empirically relevant parameters: 40 characters, 40 states, a dropout rate of 18%, and a mutation rate of 2.5%. For the purposes of estimating the TreeMetRate, *P* contained uniform probabilities of transitioning to any other tissue from a given tissue.

For each tree, the AlleleMetRate and TreeMetRate were calculated directly from the cells or tree; the Tissue Dispersion Score was calculated with a background derived from the tissue distribution across the 1,000 trees. Performance was assessed by the agreement between a specific score and the underlying rate of metastasis for that tree, as in **Fig. S9A-D**.

#### Bootstrapping analysis of TreeMetRate stability

Bootstrapping analysis was performed on a set of 10 simulated trees from (*34*). For each simulated tree, we sampled with replacement the cut-sites of the character matrices 100 times, resulting in 100 “bootstrapped character matrices” for each tree. We reconstructed each of these bootstrapped character matrices with Cassiopeia-ILP (maximum_neighborhood_size = 10,000, time_limit = 12,600). This left us with 1,000 reconstructions where each reconstruction corresponded to one of 10 simulated trees.

Then, for each simulated tree, we overlaid 50 metastatic processes on to the tree (here, because the tree was already defined, it was not necessary to use α). Each time, we transferred the labels to each of the 100 reconstructions for that simulated tree and evaluated the mean TreeMetRate, as well as the standard error, defined as:

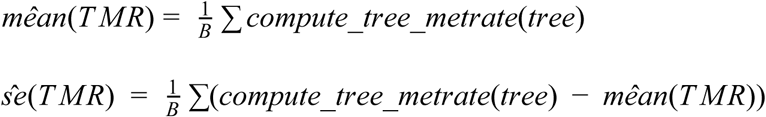

where *B* is the number of bootstrapped samples for a given metastatic process (here, *B = 100)*. The coefficient variation for each metastatic process was defined as the ratio 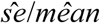.

#### Assessing accuracy of the tissue transition matrices

We evaluated the accuracy of conditional tissue transition matrix inference by utilizing a similar approach above: we simulated trees (α ∼ *Unif* (0, 0.7), μ ∼ *U nif* (0, 0.3)) with *D* ∈ {10, 14} that allowed cells to move between 6 tissues (these probabilities were specified in the conditional probability matrix *P*). For each simulation, we evaluated how well a particular statistic was able to capture the underlying *conditional* tissue transition matrix *P’* (i.e. the non-diagonal probabilities, corresponding to the probability that a cell will move to some tissue, given it is found in a particular tissue and is metastasizing) calculated from the ground-truth internal labels.

Conditional tissue transition matrices were simulated according to two models: one where the rates of metastasizing to any other tissue were uniform (hereafter referred to as the “Uniform” model) and one where rates could be biased to some tissues, i.e., experience various tissue tropisms (hereafter referred to as the “Biased” model). To model both scenarios, we sampled each tissue’s conditional transition probabilities from a Dirichlet distribution, parameterized with a length 5 array 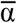 (recall that we are simulating conditional tissue transitions, and are not concerned with the probability that a cell remains in place) that essentially provides the “density” towards any outcome. To simulate the “Uniform” model, 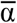 was parameterized as [50, 50, 50, 50, 50] - yielding roughly uniform probabilities across the 5 tissues. To simulate the “Biased” model, we used a “flat Dirichlet distribution”, corresponding to 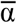 parameterized as [1, 1, 1, 1, 1], which yielded potentially very biased probability distributions. We simulated 500 trees per model.

We used three algorithms for inferring the conditional tissue transition matrix *P* for a given tree:

1. *FitchCount* (as described above, and in the **Supplementary Materials**).
2. Naive-Fitch (in which the internal nodes were assigned from a single solution drawn from the Fitch-Hartigan top-down algorithm and used to infer *P’*).
3. Majority-Vote (in which the internal nodes were assigned the label that appeared at the greatest frequency at the leaves below the node and these assignments were used to infer *P’*).

In both the Naive-Fitch and Majority-Vote case, after internal nodes were assigned a label we performed a depth-first traversal on the tree and counted the number of times a parent transitioned to a child. Ground-truth conditional transition matrices were found similarly, using the simulated ground-truth labels for internal nodes.

To evaluate accuracy, we computed the Spearman correlation (using Python’s scipy library, version 1.2.2) between the flattened matrix of the ground-truth conditional transition matrix and the transition matrices inferred by one of the three algorithms described above (*FitchCount*, Naive-Fitch, and Majority-Vote).

### Supplementary Figures and Legends

**Fig. S1.**
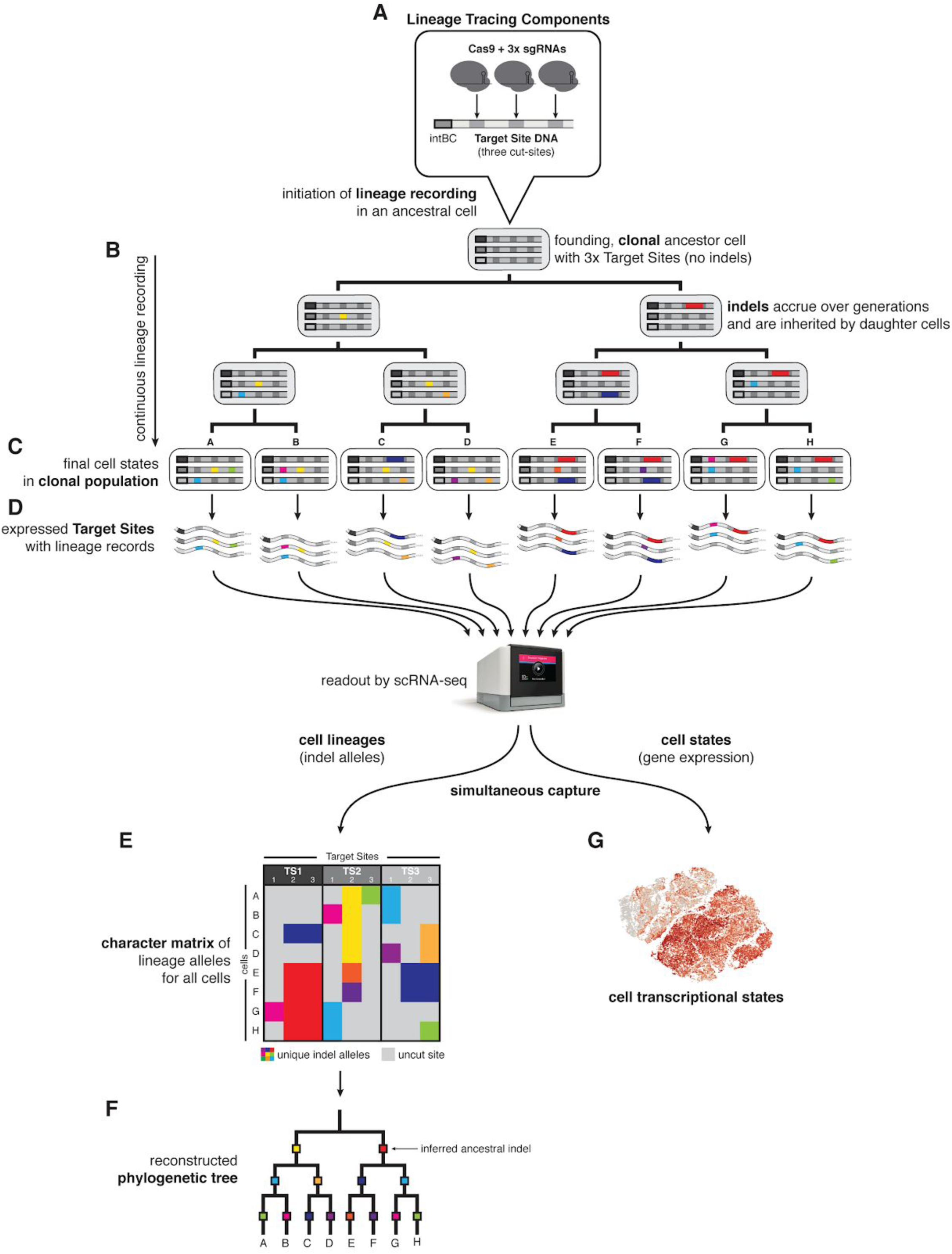
Detailed schematic of lineage tracing methodology. (**A**) Cells are genetically engineered with the lineage tracing components: (i) Cas9, (ii) multiple copies of the Target Site, and finally (iii) three sgRNAs that are complementary to three cut-sites on each Target Site. Multiple copies of the Target Site per cell are distinguished by unique integration barcodes (intBCs). Cas9-induced double-stranded breaks at cut-sites are repaired with high-diversity insertions or deletions (indels), which act as stable, heritable markers of cell lineage. (**B**) Upon initiation of lineage recording in a founding cell (i.e., one “clone”), indels continuously accrue on the Target Sites (colored boxes), which are inherited by descendents over subsequent generations. (**C**) At the end of the lineage recording experiment, the final population of descendent cells (i.e., one “clonal population”) are collected and (**D**) their expressed Target Site mRNAs are captured by single-cell RNA-sequencing (e.g., by the 10X Genomics Chromium platform) alongside transcriptome-wide expressed genes. (**E**) The indel allele information is read from the Target Site sequences and summarized in a “character matrix” of indel allele states (values) for each cut-site (columns) in each cell (rows). (**F**) From the pattern of cells’ shared and distinguishing indel alleles, a tree reconstruction algorithm builds a phylogenetic model that best captures cell–cell relationships (e.g., by maximizing parsimony), thus producing a detailed map of cell lineage. (**G**) Simultaneously, single-cell RNA-sequencing captures the transcriptional profiles of the observed cells, allowing for direct comparisons between cell lineage and cell state.

**Fig. S2.**
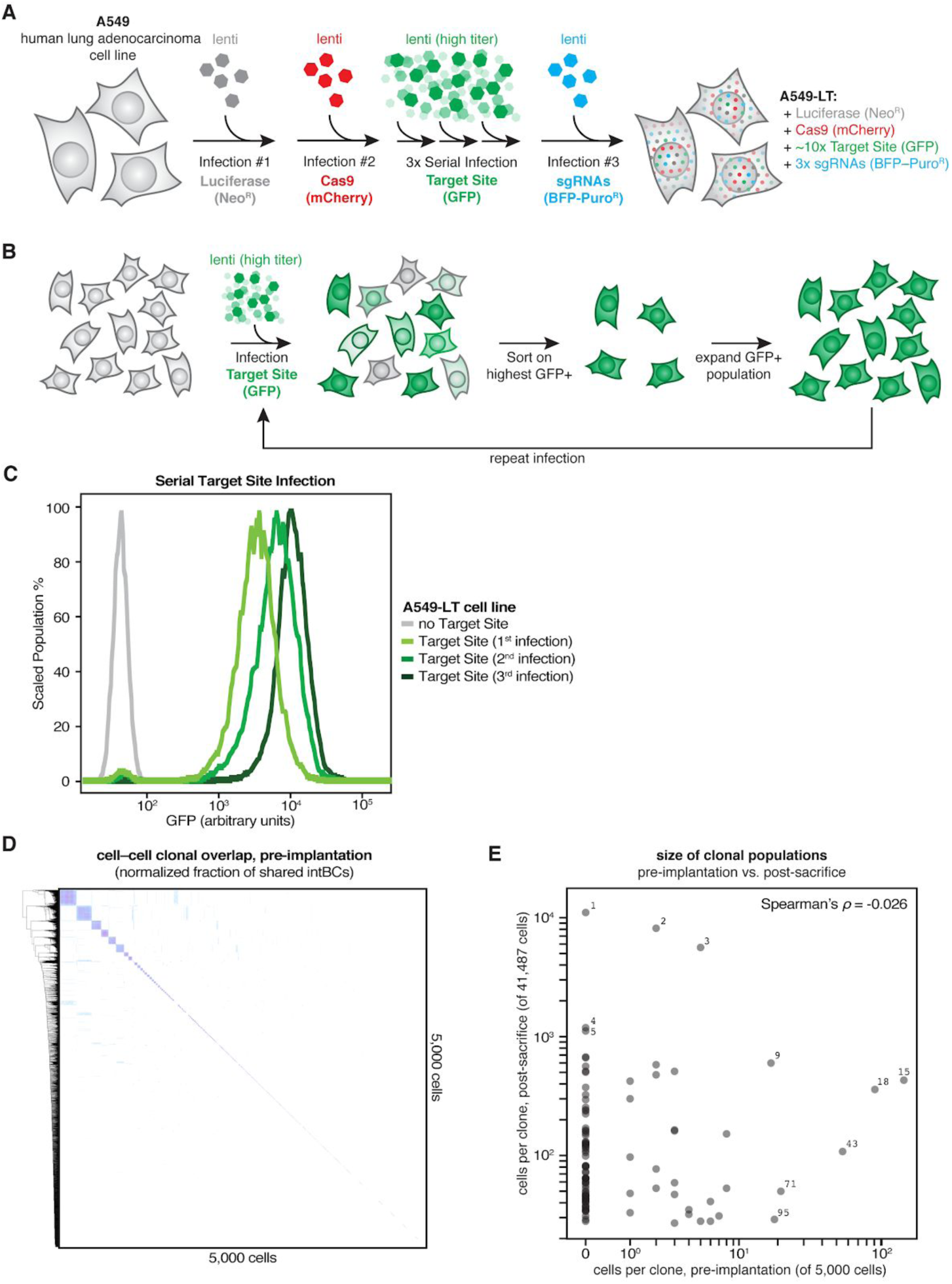
Cell line engineering strategy and estimation of clonal diversity. (**A**) Human lung adenocarcinoma (A549) cells were genetically engineered with the lineage tracing components by lentiviral transduction with (1) Luciferase-Neomycin^R^ and antibiotic-selected, (2) Cas9-mCherry and fluorescence-sorted, (3) serial, high-titer TargetSite-GFP and fluorescence-sorted, and finally (4) triple-sgRNA BFP-Puromycin^R^ and fluorescence-sorted, thus producing lineage tracing-competent A549-LT cells. (**B**) Serial, high-titer TargetSite-GFP lentiviral transduction strategy to achieve high copy-number. (**C**) Cells with high-shifted GFP fluorescence after successive TargetSite-GFP infections, indicating increasing copy-number of the Target Site. (**D**) Sample of 5,000 A549-LT cells prior to injection to estimate initial clonal diversity. Shown is a heat-map of the fraction of shared intBCs in all cell–cell comparisons. On average, approximately 22,000 unique, high-quality intBCs were identified per random sample of 5,000 cells; thus, we estimate approximately 2,150 distinct clones per 5,000 cells (assuming 10.3 intBCs per clonal population; **Fig. S6A**) at the beginning of the experiment. (**E**) Comparison of the size of each clonal population observed in mouse M5k in a pre-implantation sample of 5,000 cells (*in vitro*) and post-sacrifice (*in vivo*). There is no correlation between the *in vitro* and *in vivo* population sizes (Spearman’s *ρ*=-0.026).

**Fig. S3.**
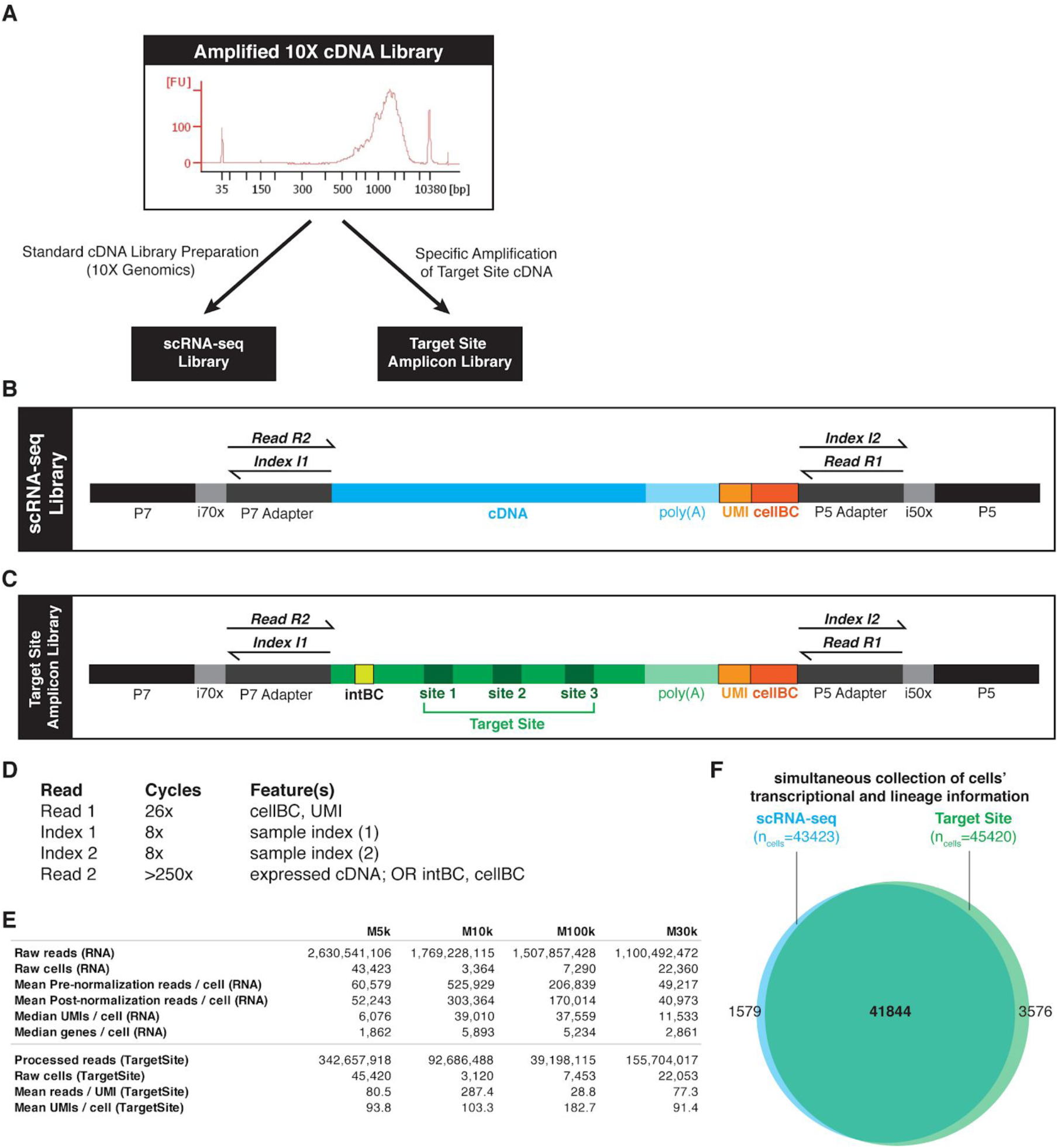
Sequencing library construction and metrics. (**A**) The amplified cDNA (shown here as a BioAnalyzer trace) from the Chromium 3’ Single Cell V2 kit (10X Genomics) serves as the template for both (**B**) the single-cell gene expression (“RNA”) library and (**C**) the single-cell Target Site amplicon library (Methods). (**D**) The cellBC and unique molecular identifier (UMI) are sequenced from Read R1, the sample identities are sequenced from Indices I1 and I2, and the expressed cDNA or the Target Site amplicon (including the intBC and cut-sites 1, 2, and 3) are sequenced from Read R2. (**E**) Library sequencing metrics for TargetSite and RNA libraries for each mouse. (**F**) There is vast overlap in the cells identified from the gene expression and lineage sequencing datasets, as shown for mouse M5k.

**Fig. S4.**
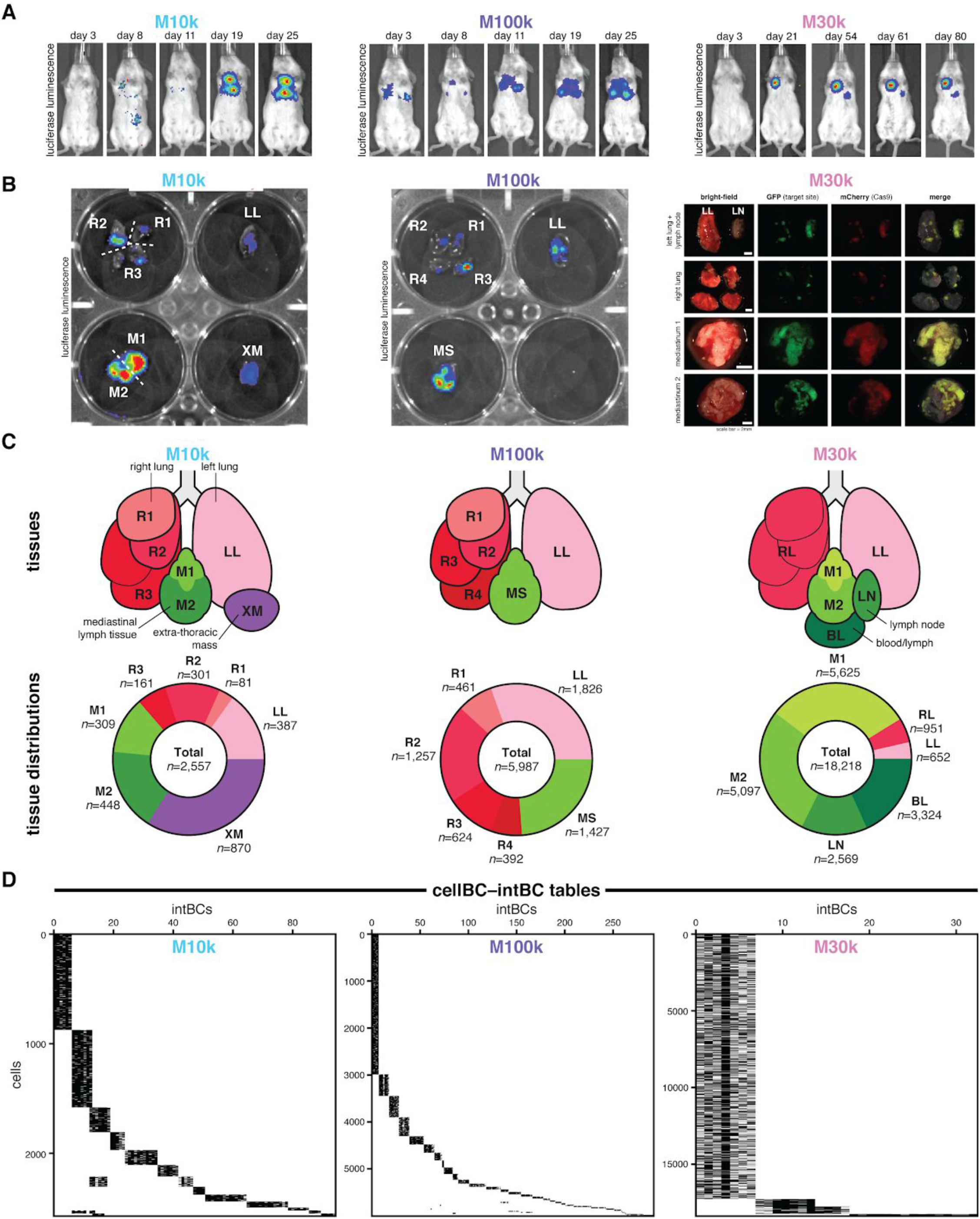
Tracing the cell lineages of metastatic progression in three additional mice. In addition to mouse M5k discussed in throughout the main text, we also traced the lineages of metastatic dissemination in three additional mice orthotopically xenografted with 10,000 (M10k, left), 100,000 (M100k, middle), and 30,000 (M30k, right) A549-LT cells in two cohort experiments (first cohort, A549-LT1: M10k and M100k; second cohort, A549-LT2: M5k and M30k). (**A**) *In vivo* bioluminescence imaging of cancer cell engraftment and metastatic spread at indicated times post-implantation. The mice were sacrificed 53, 67, and 80 days post-implantation, respectively. (**B**) *Ex vivo* imaging of tumorous tissues by bioluminescence (left and middle) or fluorescence (right) imaging with the tissue samples indicated. In addition to extensive tumors in the lungs and mediastinum, M10k had one large solid tumor located ventral to the left lung and embedded in the ribcage (called here an “extra-thoracic mass”; XM). (**C**) Anatomical representation of collected tumorous tissues (top) and the number of cells collected for each tissue and each mouse (bottom). Notably, M30k had a large lymph node (LN) on the left lung, as well as diffuse bloody lymph (BL) in the thoracic cavity upon sacrificing. (**D**) CellBC-intBC tables showing clonal populations of cancer cells in each mouse, as in **Fig. S5A,B**. Some intBCs are shared between some clones (most notably in M10k) which likely resulted from cells that were clonally related at the stage of serial Target Site transduction during cell line engineering (Methods).

**Fig. S5.**
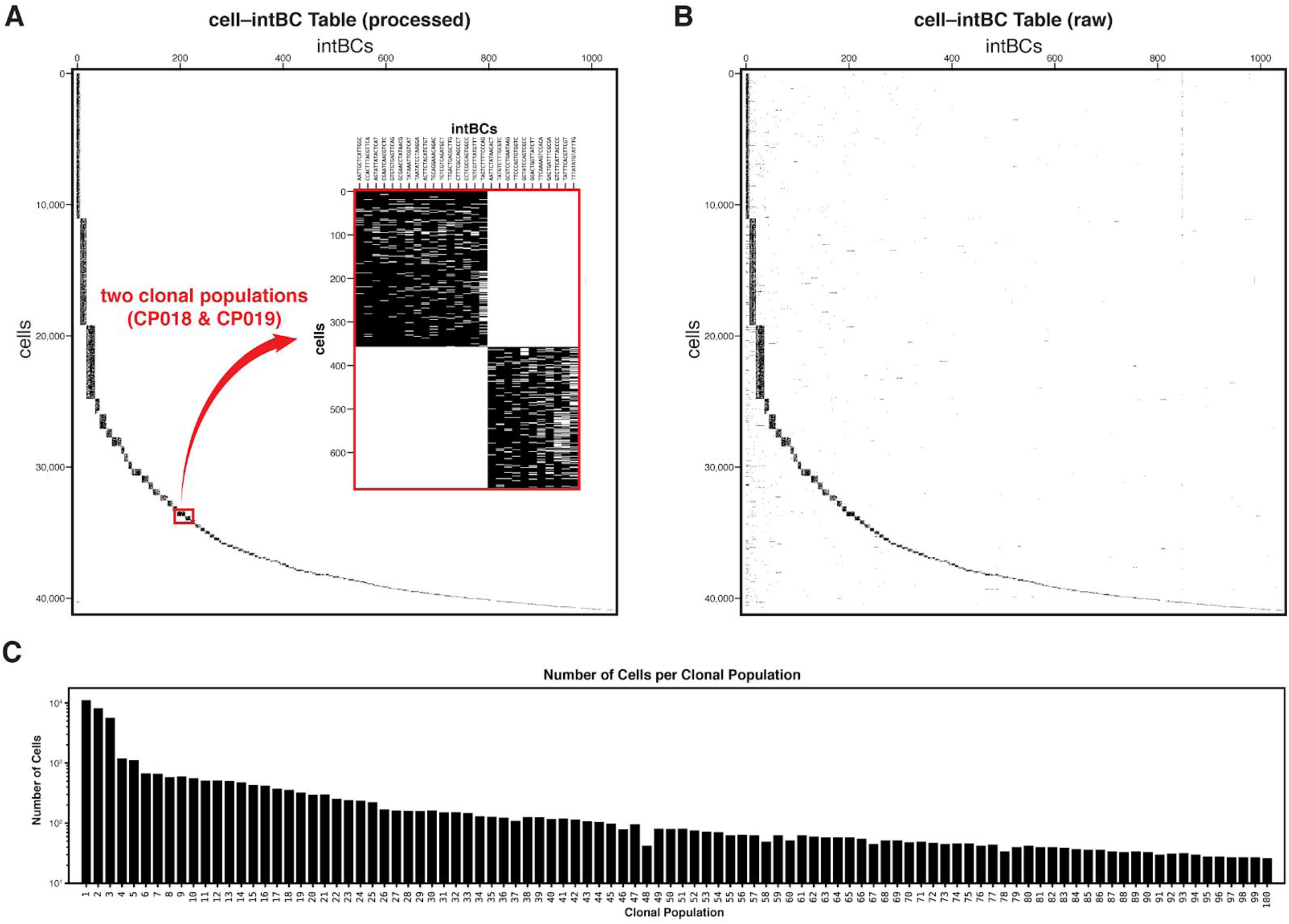
Identifying clonal populations by shared integration barcodes (intBCs). (**A** and **B**) Tables representing the >1,000 unique integration barcodes (intBCs; columns) and >40,000 cells (rows) observed in mouse M5k, after processing (**A**) and before processing (**B**). Because intBCs are clonally inherited, cells that share identical intBCs are grouped into clonal populations (here, black blocks). (**A; inset**) The set of intBCs is generally exclusive to a single clonal population, such as those observed in clonal populations #18 and #19 (CP018 and CP109). (**B**) In the raw, unprocessed cell–intBC table, cells may be associated with intBCs that are not in their defined clonal set (indicated here by density outside of the black blocks). Through the processing pipeline, these conflicting intBCs (and/or the cells to which they pertain) are identified as cell doublets, cell-free transcripts in emulsion droplets, or sequencing artifacts and removed (Methods). (**C**) The number of cells per clonal population, generally numbered by descending population size.

**Fig. S6.**
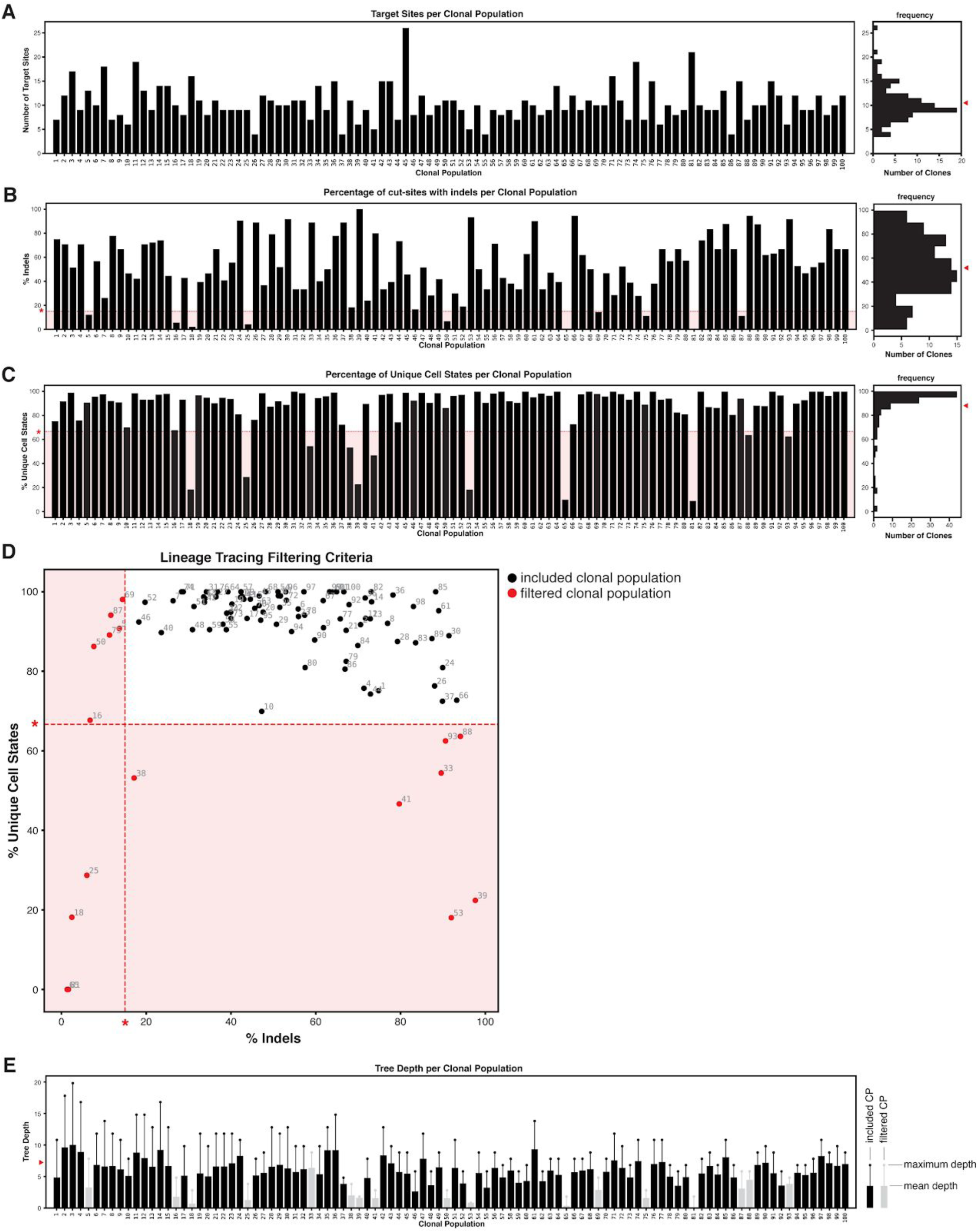
Characteristics of the lineage tracer and quality-control of clonal populations. (**A**) The copy-number of integrated Target Sites per clonal population, as determined by the number of unique intBCs. (**A, right**) The distribution of Target Site copy-number per clonal population; mean copy-number of Target Sites indicated (red arrowhead). (**B**) The percentage of cut-sites bearing lineage indel alleles per clonal population. Clonal populations with <15% indels are excluded (red asterisk, red underlay). (**B, right**) The distribution of indel-bearing cut-sites per clonal population; mean percentage indicated (red arrowhead). (**C**) The percentage of unique cell lineage states per clonal population (i.e., lineage diversity). Clonal populations with <66.7% diversity were excluded (red asterisk, red underlay). (**C, right**) The distribution of lineage diversity per clonal population; mean percentage indicated (red arrowhead). (**D**) Comparison of lineage tracing characteristics (% indels and % unique cell states) to define quality-control filtering criteria. Clonal populations removed by the filter (red closed circles) and filtering thresholds (red asterisks, red underlay) are indicated. (**E**) The depth of the reconstructed phylogenetic trees for each clonal population. Mean tree depths are shown as closed bars; maximum tree depths are shown as whiskers. Clonal populations that were excluded due to suboptimal lineage tracing characteristics (gray) and the average tree depth across all clonal populations (red arrowhead) are indicated.

**Fig. S7.**
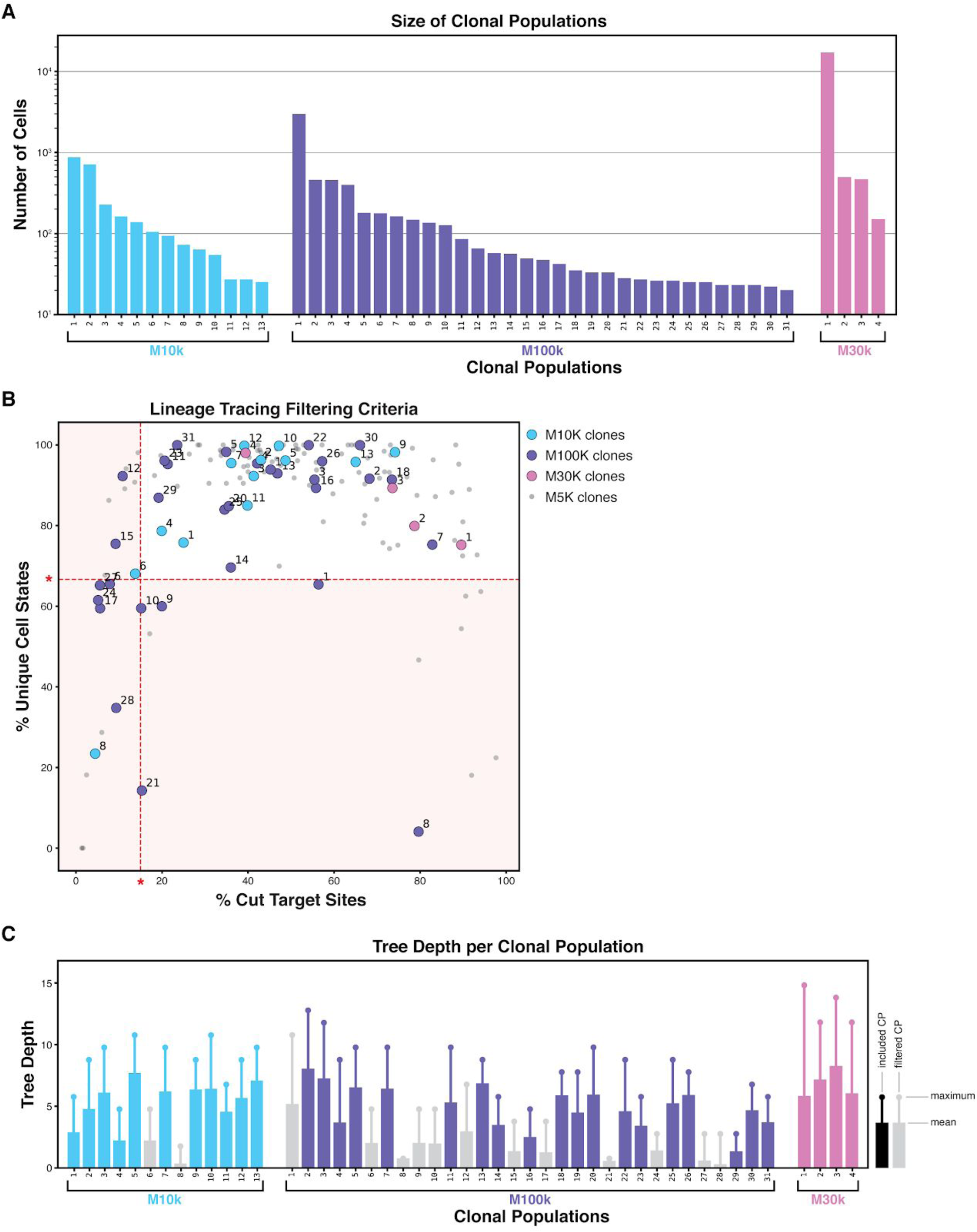
Lineage tracing characteristics of clonal populations in additional mice. (**A**) Number of cells in each clonal population in M10k, M100k, and M30k mice. (**B**) Scatter plot of the percentage of cut-sites bearing indels and the percentage of unique cell states per clonal population, which are characteristics of the lineage tracer that influence tree reconstructability. Some clonal populations exhibited suboptimal parameters (red asterisks and red field) and were excluded from reconstruction and downstream analyses. (**C**) The mean and maximum depths of the reconstructed phylogenetic trees for each clonal population in additional mice, as in **Fig. S6E**.

**Fig. S8.**
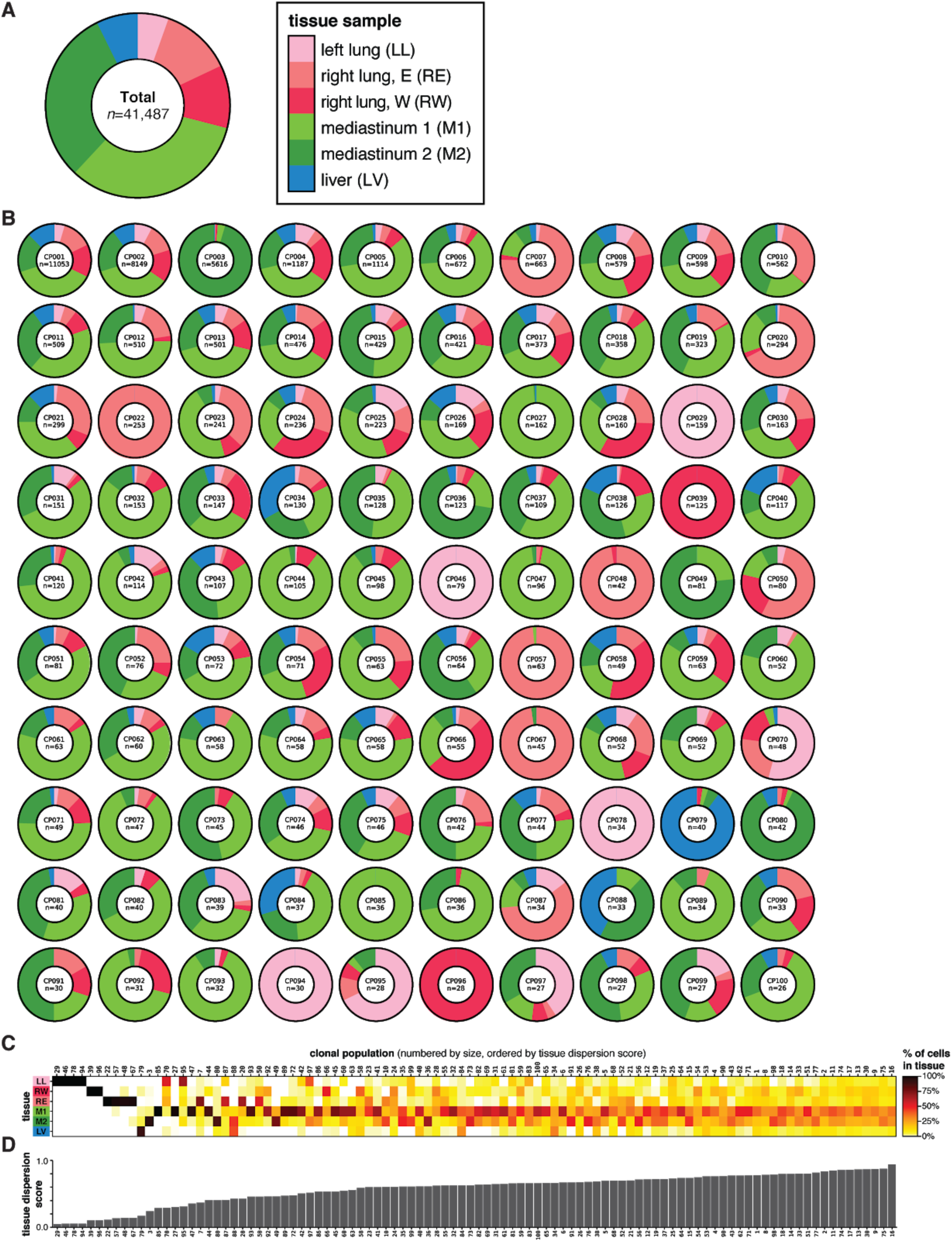
Clonal populations exhibit distinct tissue distributions. (**A**) The bulk distribution of all collected cells across the six tissue samples. (**B**) The distributions of cells from each clonal population across the six tissue samples. Some clonal populations were exclusive to the primary tissue (e.g., clonal population CP046), whereas some clones exhibited biased tissue distributions (e.g., CP003) and many others were observed broadly distributed across all tissues (e.g., CP011). (**C**) Tissue distributions of the largest 100 clonal populations. (**D**) The Tissue Dispersion Score is a statistical measurement of the distribution across tissues for each clonal population; *x*-axis shared with (**C**).

**Fig. S9.**
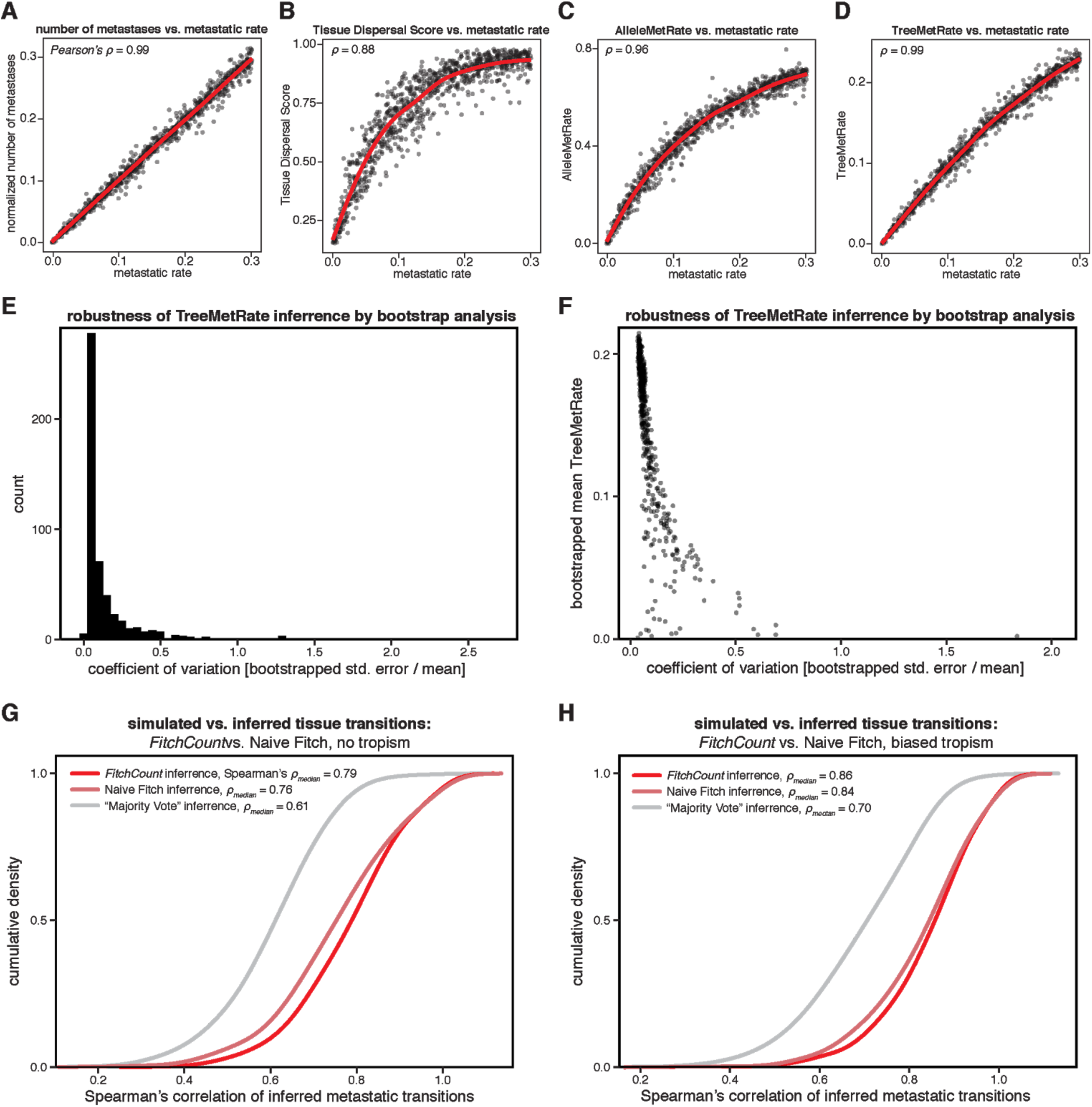
Assessing the accuracy of different measurements of metastatic rate and inference of tissue transitions using simulated lineages. (**A–D**) Comparison between the simulated metastatic rate and various lineage tracer-derived statistics, with the correlation (Pearson’s *ρ*) indicated; red lines represent moving average. (**A**) The normalized count of simulated metastatic transitions is very well correlated with the simulated metastatic rates, and serves as a ground-truth benchmark. (**B**) The Tissue Dispersal Score, which is a statistical measure of how closely a clone’s tissue distribution matches the background tissue distributions, is correlated with the metastatic rate, but saturates at intermediate metastatic regimes. (**C**) The AlleleMetRate, or the proportion of cells whose closest relative by allele similarity is in a different tissue, is better correlated with metastatic rate. (**D**) The TreeMetRate, or the proportion of inferred metastases in a reconstructed phylogeny, is the best lineage-derived measurement of metastatic phenotype by correlation. (**E** and **F**) Robustness of the TreeMetRate inference by bootstrap analysis. (**E**) The distribution of the coefficients of variation of the TreeMetRate across 50,000 simulated, bootstrapped phylogenies (Methods). The small coefficient of variation indicates that the TreeMetRate is a robust measurement. (**F**) The TreeMetRate coefficient of variation is only relatively large when the mean TreeMetRate is small. (**G** and **H**) Cumulative density plots assessing the accuracy of the *FitchCount* strategy for inferring ancestral tissue transition from simulated phylogenies with or without simulated biased tissue transitions (thus approximating tropism; Methods: “*Assessing accuracy of the tissue transition matrices*”). *FitchCount* outperforms other inference approaches, as measured by the Spearman correlation between the inferred and the ground-truth conditional tissue transition probability matrices. *FitchCount* was benchmarked against two other inference methods: (i) a “naive” single-solution implementation of the Fitch-Hartigan maximum parsimony algorithm or (ii) “Majority Vote”, which infers ancestral tissue location as the tissue in which the majority of cells below each clade-level reside. Conditional probabilities for each algorithm were obtained by row-normalizing the count matrix with respect to the non-diagonal counts in each row.

**Fig. S10.**
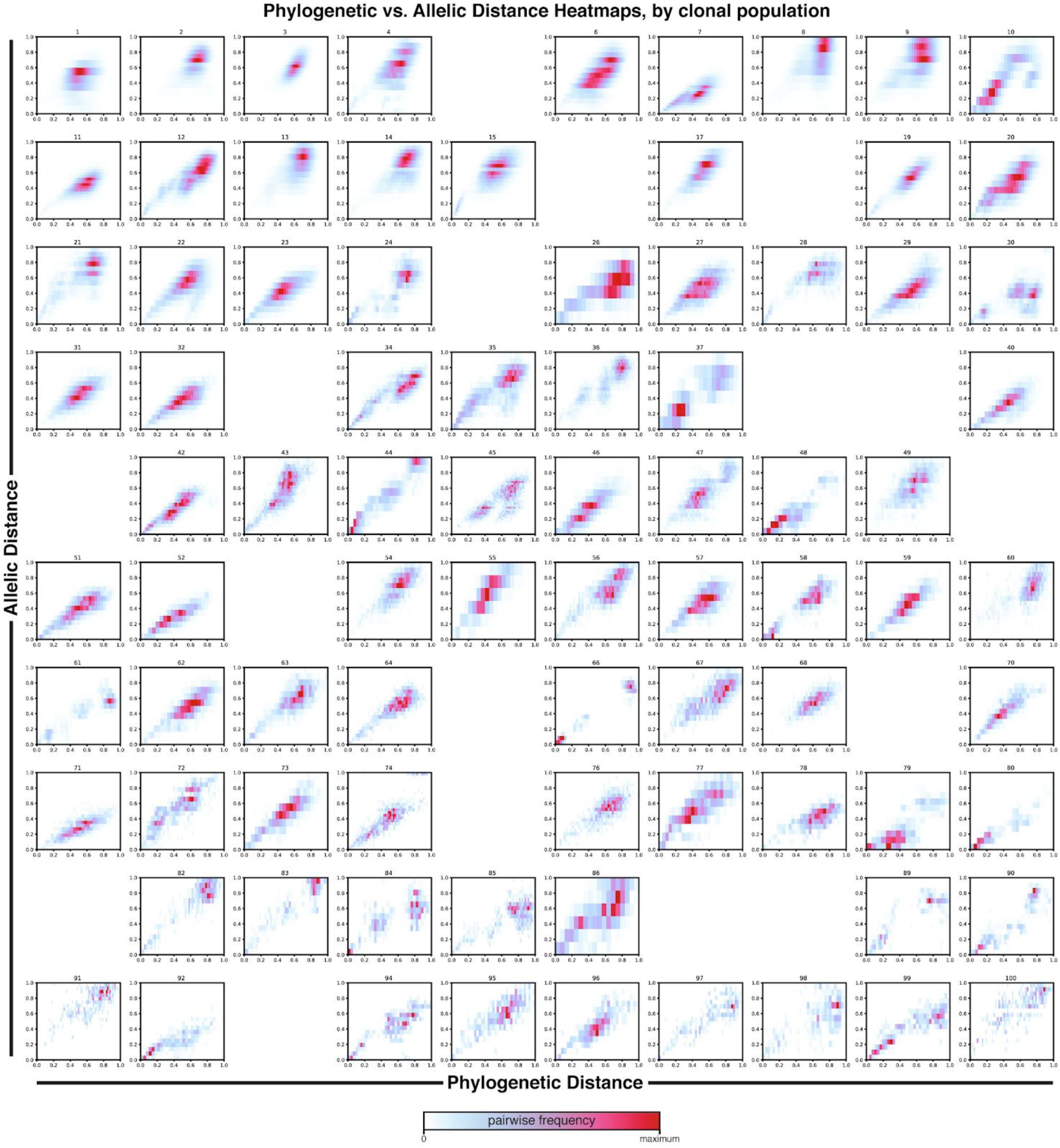
Relationship between phylogenetic distance and allelic distance for each clonal population. Density heat-maps comparing phylogenetic distance (i.e., the normalized tree branch distance between two cells) and allelic distance (i.e., the normalized difference in lineage allele state between two cells) for all pairwise cell–cell relationships and for each clonal population, as in **Fig. 2C**. As expected, phylogenetic and allelic distances are correlated, suggesting that the reconstructed trees are a good phylogenetic model of cell–cell relationships. Excluded clonal populations are not shown.

**Fig. S11.**
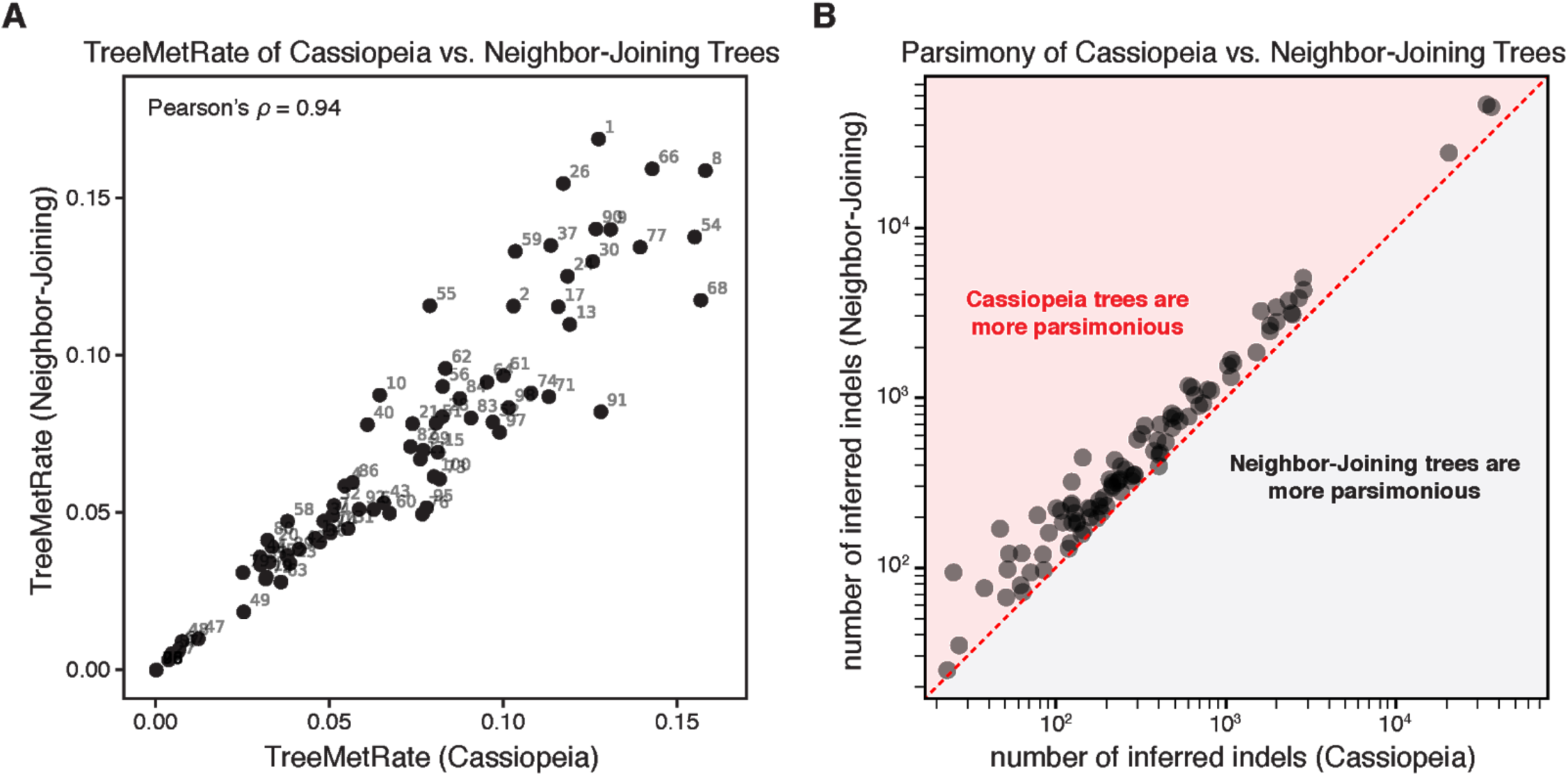
The TreeMetRate is stable across tree reconstruction algorithms (Cassiopeia versus Neighbor-Joining). (**A**) Comparison of the TreeMetRates for each clonal population from Cassiopeia trees and Neighbor-Joining trees. The TreeMetRates are correlated for both the Cassiopeia and Neighbor-Joining trees (Pearson’s *ρ*=0.94). (**B**) Comparison of the parsimony of Cassiopeia trees and Neighbor-Joining trees, defined as the number of inferred indels in each tree. Notably, the Cassiopeia trees are more parsimonious than the Neighbor-Joining trees (i.e., they have fewer inferred indels; red overlay).

**Fig. S12.**
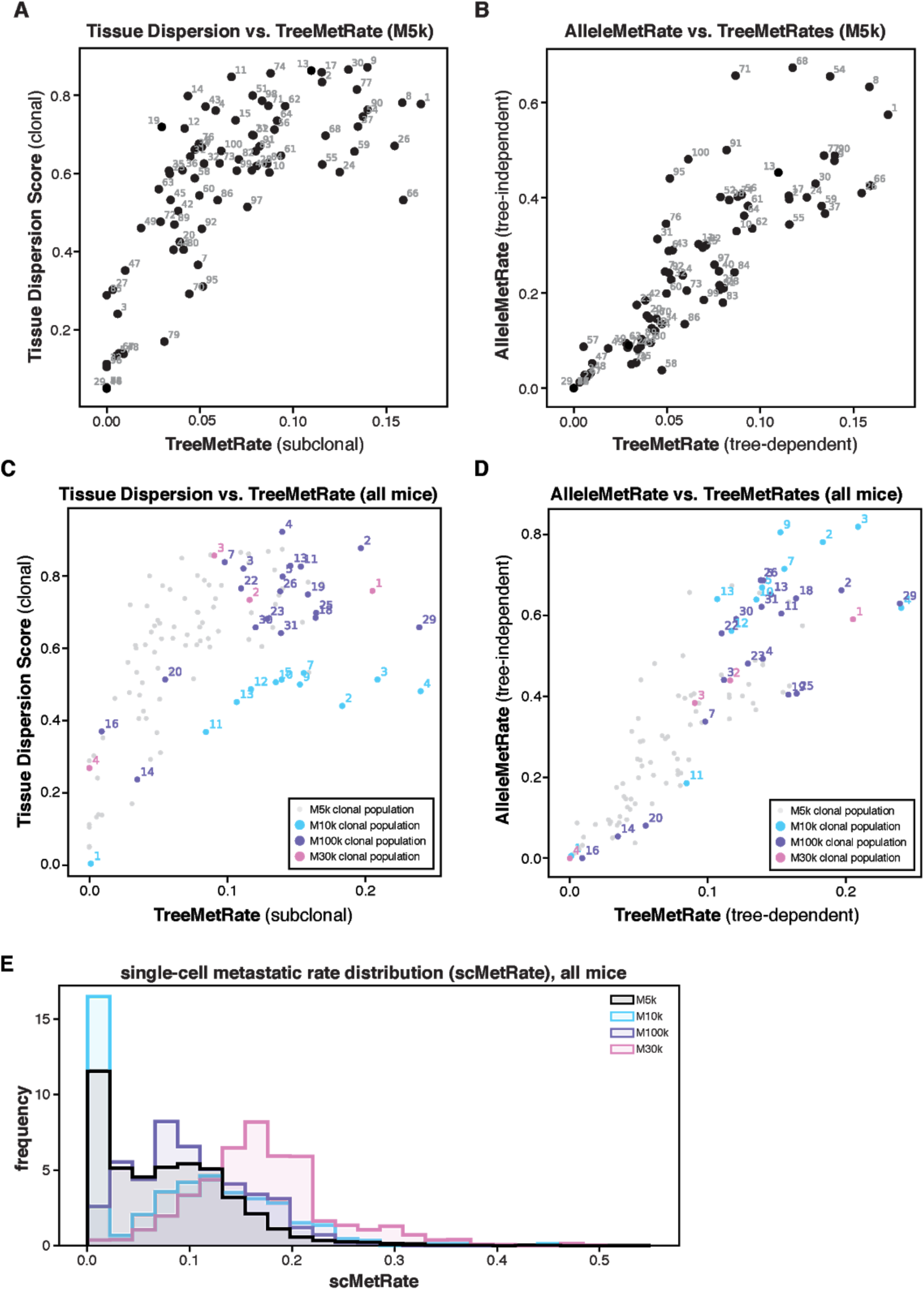
Clonal populations exhibit broad metastatic phenotypes, measured by Tissue Dispersal Score, AlleleMetRate, and TreeMetRate. (**A–D**) We evaluated the metastatic phenotype of each clonal population from mouse M5k (**A, B**) and the additional mice M10k, M100k, and M30k (**C, D**) using our three lineage-derived measurements: Tissue Dispersal Score (as in **Fig. S8D**), AlleleMetRate, and TreeMetRate (as in **Fig. 3C**). These three measurements follow similar relative trends across all four mouse experiments. First, the clonal populations exhibit a broad range of metastatic phenotypes. Second, all three measurements are correlated with one another, though simulations indicate that the TreeMetRate is the most accurate for estimating the underlying metastatic rate (**Fig. S9**). Third, Tissue Dispersal Score saturates at intermediate metastatic regimes (**A** and **C**). (**E**) The distribution of single-cell-resolution metastatic rates (scMetRates) across all cells for each mouse (as in **Fig. 3D**). Though all mice have broad distributions of metastatic phenotypes, mouse M5k (black) is particularly well represented by cells in low-to-intermediate metastatic regimes whereas mouse M30k (pink) has very few cells in the low metastatic regime.

**Fig. S13.**
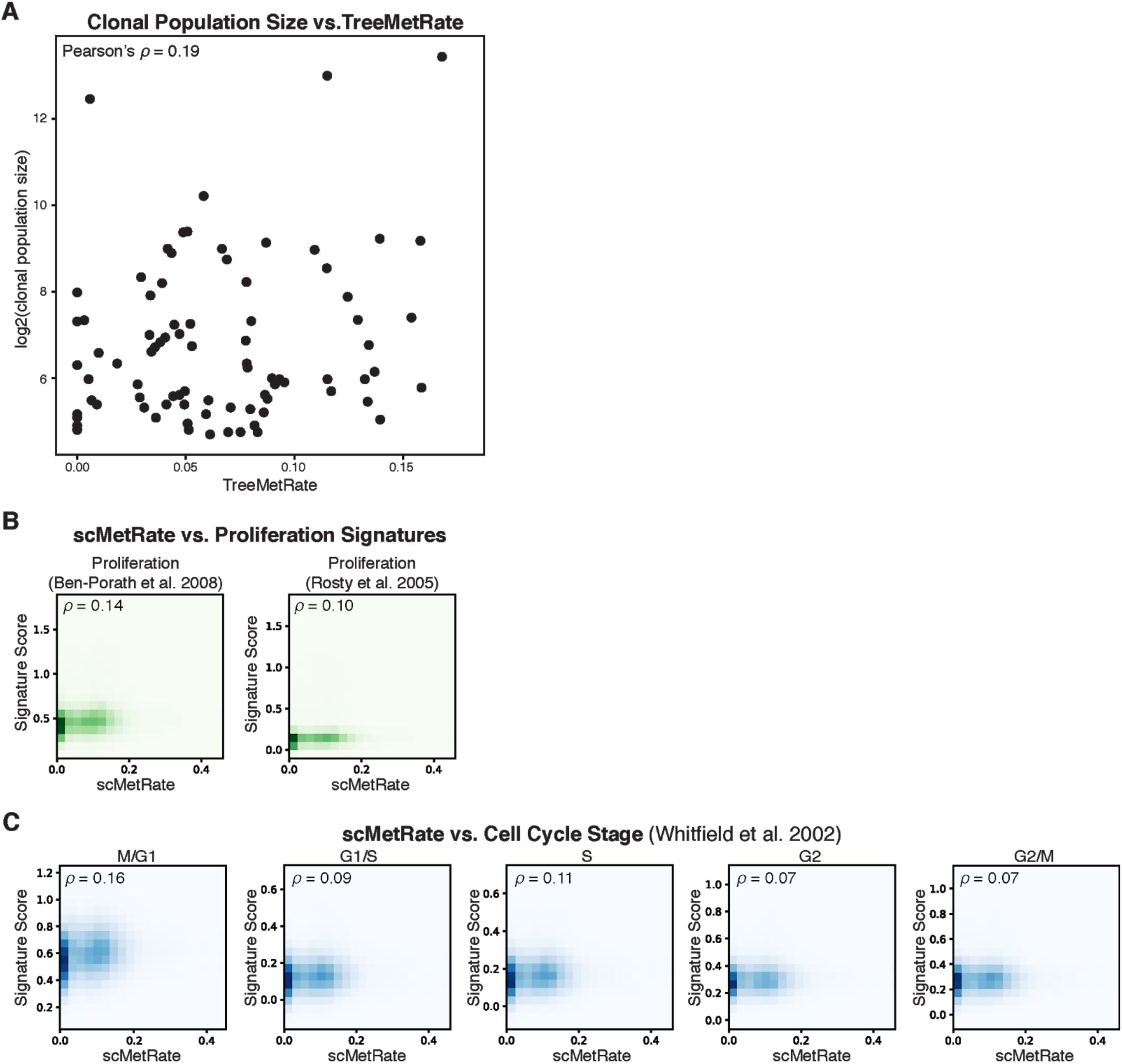
The scMetRate measures metastatic potential decoupled from proliferative capacity. (**A**) There is poor correlation between the scMetRate and the log2-transformed clonal population size, a proxy for clonal fitness. (**B** and **C**) Density heat-maps comparing the scMetRate and various transcriptional signatures. Notably, the scMetRate is poorly correlated with transcriptional signatures of proliferation (**B**) nor stages of the cell cycle (**C**). Pearson’s correlations (*ρ*) are indicated for each subplot.

**Fig. S14.**
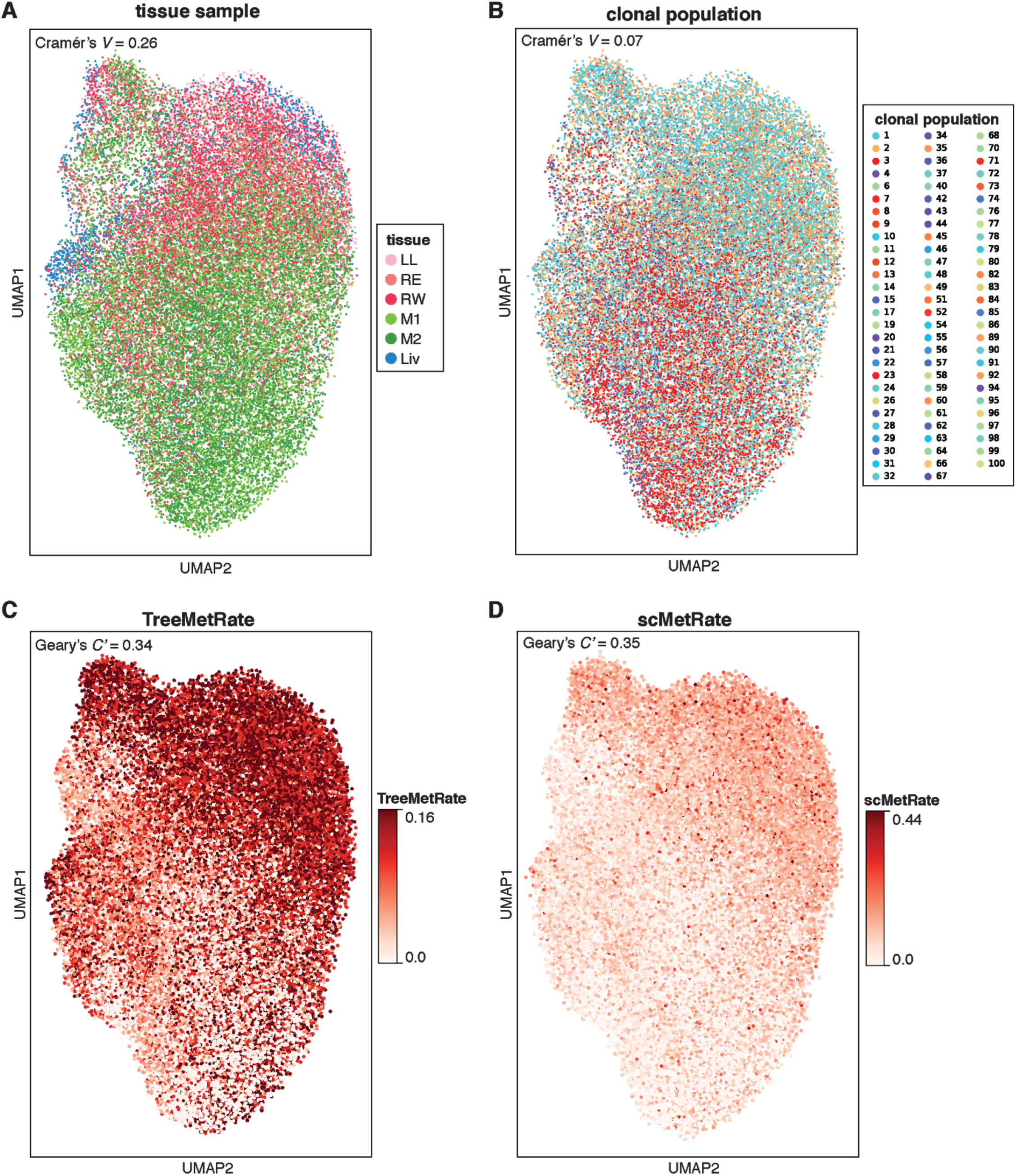
Effects of tissue sample, clonal population, and metastatic rate on transcriptional state. To identify global trends in the gene expression data, we used *Vision* (*52*) to statistically assess the transcriptional effect of four features: (**A**) tissue sample, (**B**) clonal population identity, (**C**) TreeMetRate, and (**D**) scMetRate. The transcriptional states are represented here as a two-dimensional projection using Uniform Manifold Approximation and Projection (UMAP; B). Distinctions in transcriptional state are not predominantly explained by the clonal population (**B**; by Cramér’s *V*), though there is modest association between transcriptional state and both tissue sample and metastatic phenotype (by Cramér’s *V* and inverted Geary’s *C’*, respectively).

**Fig. S15.**
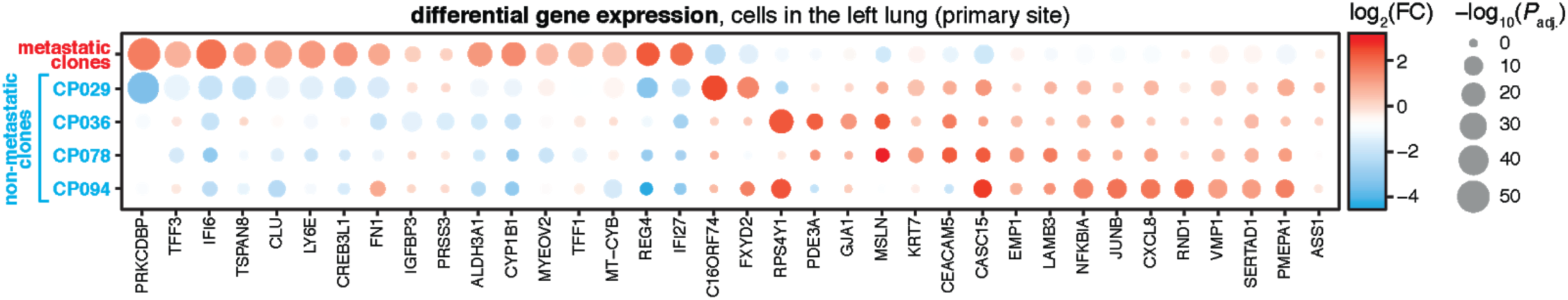
Differential expression between non-metastatic and metastatic clonal populations in the primary tissue. Differential gene expression analysis comparing four non-metastatic clonal populations (CP029, 36, 78, and 94) and all metastatic clonal populations in the primary tumor tissue (i.e., all other cells in the left lung). Significantly differentially expressed genes are colored by the log2-transformed fold-change in gene expression and scaled by the adjusted Wilcoxon rank-sum test *P*-value.

**Fig. S16.**
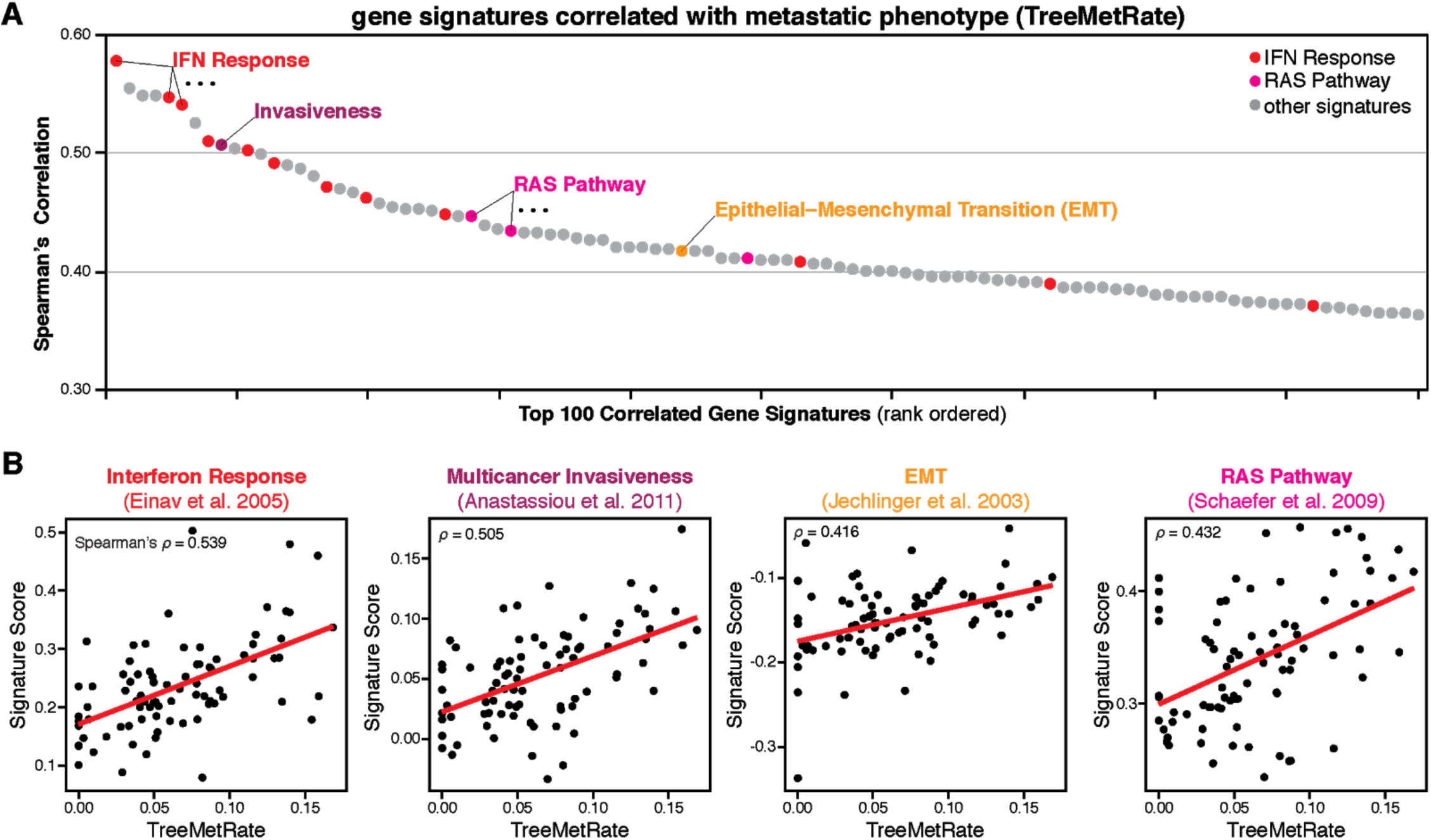
Metastasis-related gene signatures are correlated with metastatic potential. (**A**) Rank-ordered gene signatures that are the most positively correlated with TreeMetRate, including many related to interferon response (red) and RAS pathways (magenta), as well as other metastasis-related signatures. (**B**) Scatter plots showing the correlation between TreeMetRate and noted gene signature scores per clonal population; Spearman’s correlation (*ρ*) indicated.

**Fig. S17.**
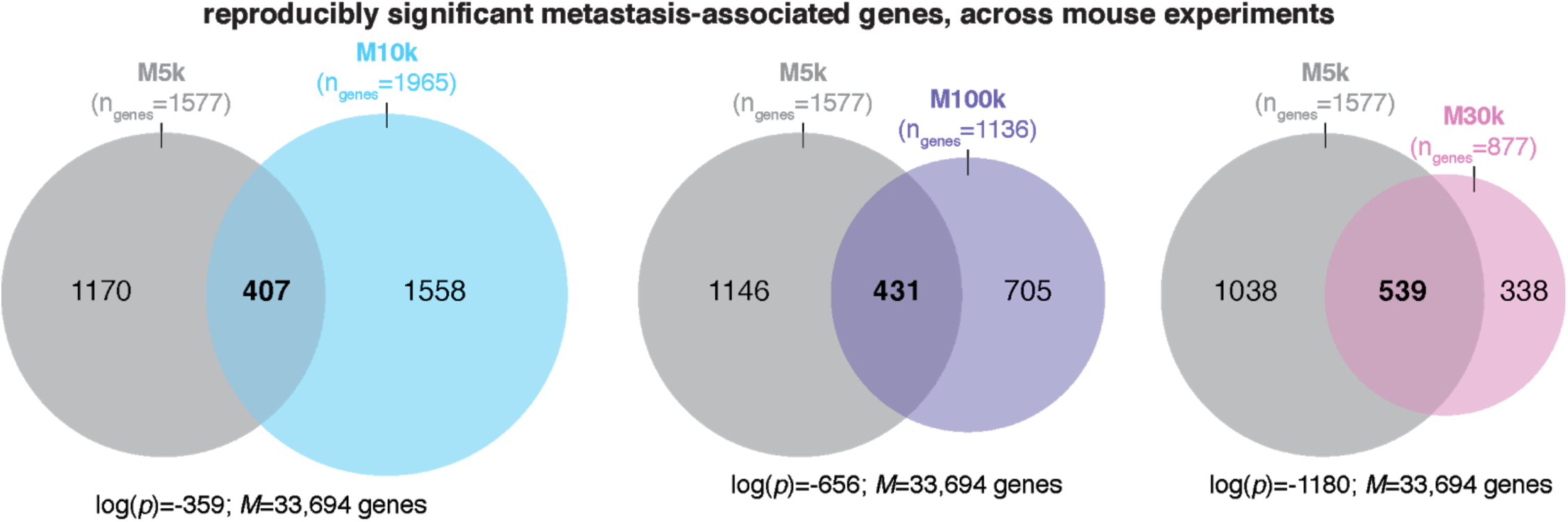
Many of the same genes are associated with metastatic phenotype across all mice. Using the same regression strategy as in the analysis of mouse M5k, we found many genes with expression that is significantly associated with high or low scMetRates. The number of significant genes for each mouse (FDR < 0.01) and their overlap in the same direction with mouse M5k (gray) are shown (*n*_*genes*_). In all cases, the overlap between mouse M5k and each additional mouse is significant by hypergeometric test (*p*-value and *M* indicated).

**Fig. S18.**
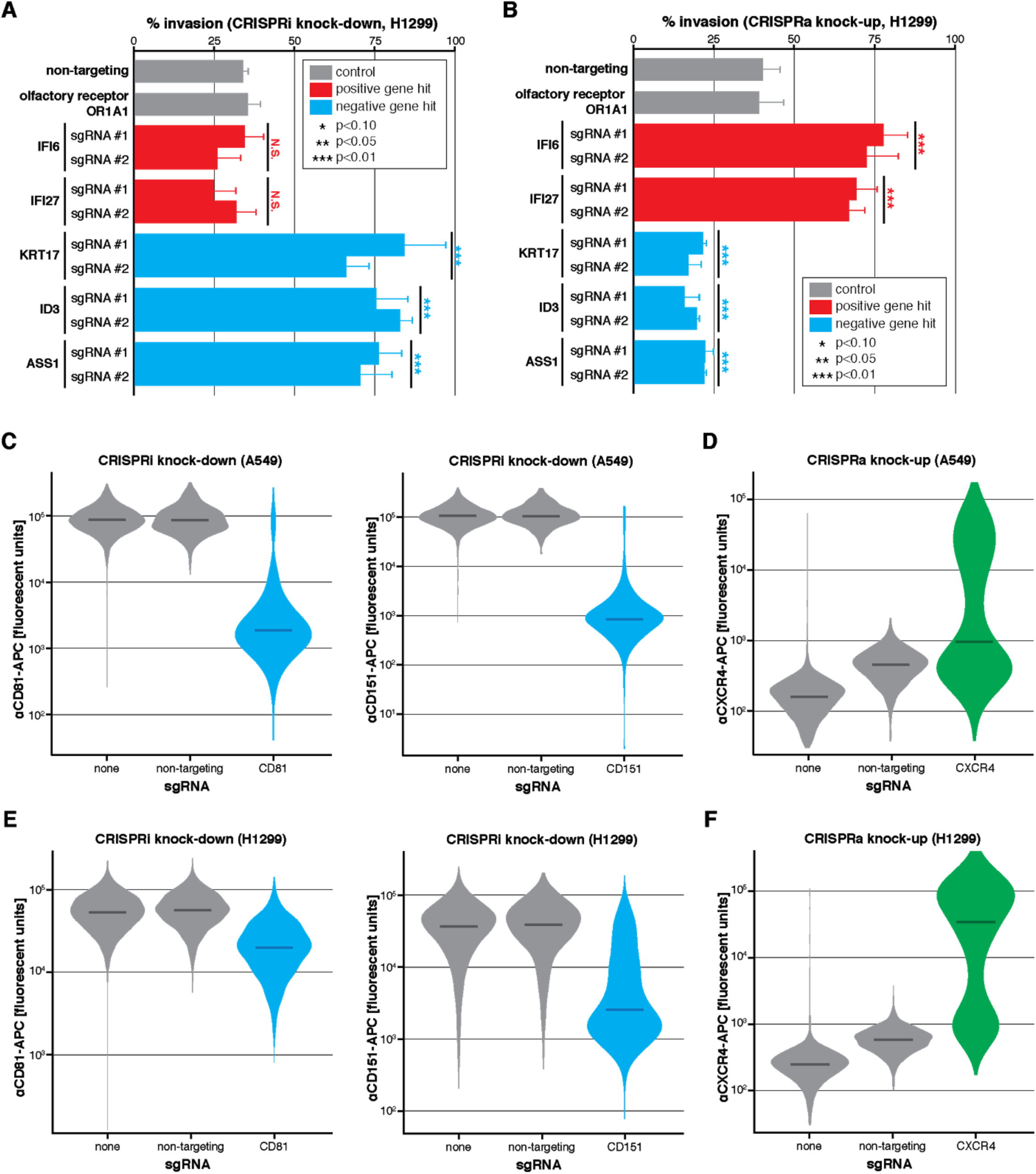
Functional validation of five gene candidates in a different cell line (H1299s) and validation of CRISPRi and CRISPRa activity. (**A** and **B**) *In vitro* transwell invasion assays following CRISPRi or CRISPRa gene perturbation, respectively, in H1299 cells; as in **Fig**.**4E, F**. Perturbation of positive and negative metastasis-associated gene candidates were performed in triplicate using two independent sgRNAs per gene. Differences in invasion phenotype relative to two negative control guides (non-targeting and olfactory receptor) were significant by two-tailed *t*-test. N.S., not significant; error bars show standard deviation. (**C, E**) A549-CRISPRi cells or H1299-CRISPRi cells, respectively, were treated with no sgRNA, non-targeting sgRNA, or sgRNAs against either CD81 or CD151, two highly expressed cell-surface markers. One week following treatment, the cells were collected, stained with APC-labelled anti-CD81 or anti-CD151 antibodies and their fluorescence was measured by flow cytometry. Shown here is substantial knock-down of CD81 or CD151 gene expression relative to no sgRNA or non-targeting sgRNA controls. (**D, F**) The same validation experiment as conducted in **C** and **E**, but for A549-CRISPRa cells or H1299-CRISPRa cells, respectively, treated with an sgRNA against CXCR4, a lowly expressed cell-surface marker. Shown here is substantially increased CXCR4 gene expression relative to no sgRNA or non-targeting sgRNA controls. Violin plots show distribution of fluorescent signal for each cell identified by flow cytometry; median marked by dark bar.

**Fig. S19.**
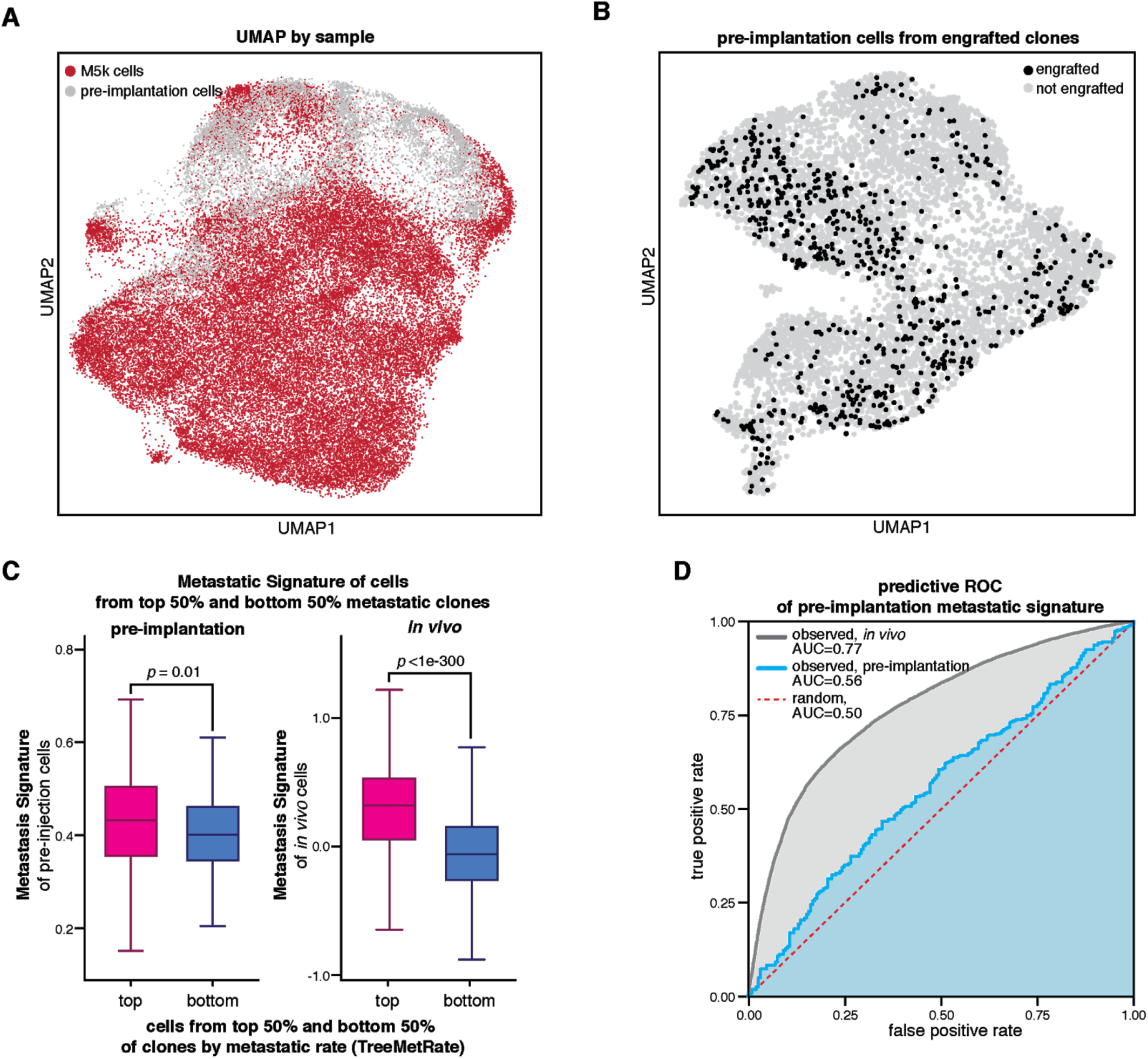
The cells in the pre-implantation pool heterogeneously express metastasis-associated genes, which are modestly predictive of their *in vivo* metastatic phenotype. (**A**) Projection of transcriptional states of M5k and pre-implantation cells, colored by sample, as in **Fig. 5A**. (**B**) Some of the cells from the pre-implantation pool could be assigned to the ∼100 clonal populations that engrafted and proliferated in mouse M5k based on their clonal barcodes (i.e., intBCs). Shown are the pre-implantation cells that could be assigned to an engrafted clone (black) on a projection of pre-implantation transcriptional states, as in **Fig. 5B,C**. (**C, left**) Pre-implantation cells from the top 50% (most) metastatic clones *in vivo* have higher Metastatic Signature scores than pre-implantation cells from the bottom 50% (least) metastatic clones *in vivo* (Mann-Whitney *U p*-value=0.01). (**C, right**) The Metastatic Signature of the most and least metastatic clones is more pronounced *in vivo* than in the pre-implantation cells (*p*-value<1e-300). (**D**) For the pre-implantation cells, the difference in Metastatic Signature scores between the most and least metastatic clones is modest, yet significant by ROC (receiver operator characteristic) analysis of false positives vs. true positives (area under the curve, AUC=0.56), indicating that the Metastatic Signature score pre-implantation is a modest predictor of *in vivo* metastatic phenotype. The predictive power for the *in vivo* population of cells is greater (AUC=0.77).

**Fig. S20.**
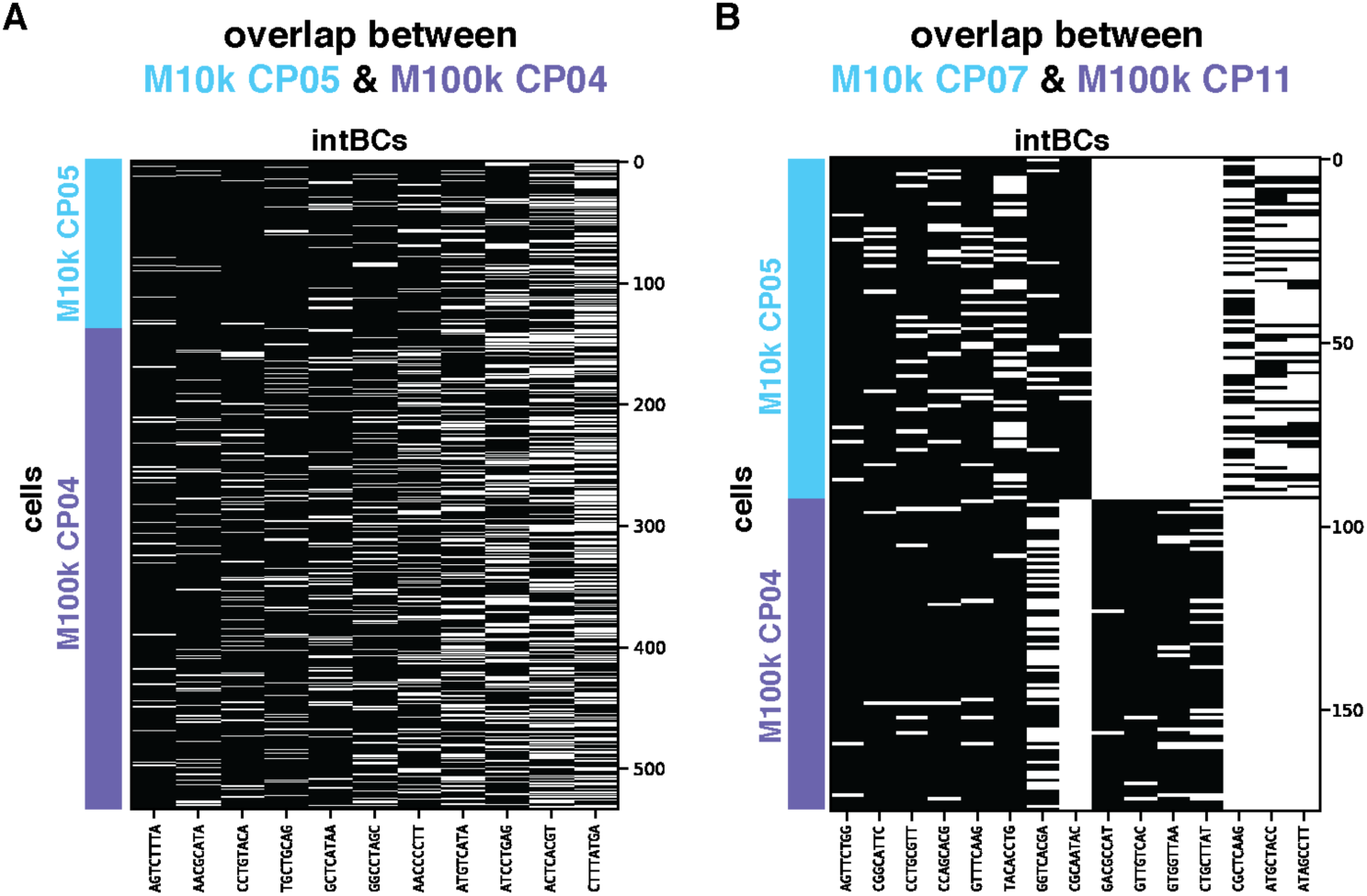
Two pairs of clonal populations from mice M10k and M100k are related, enabling an experiment to determine the robustness and reproducibility of metastatic phenotype across independent mouse experiments. Each intBC (columns) observed for each cell (rows) from (**A**) M10k CP05 and M100k CP04 and (**B**) M10k CP07 and M100k CP11. Cells from M10k are shown in light blue; M100k in purple. The clonal populations in each of these pairs are related to one another based on their shared sets of intBCs, as in **Fig. 5D**. The paired clonal populations here are the most closely related between the two mouse experiments.

**Fig. S21.**
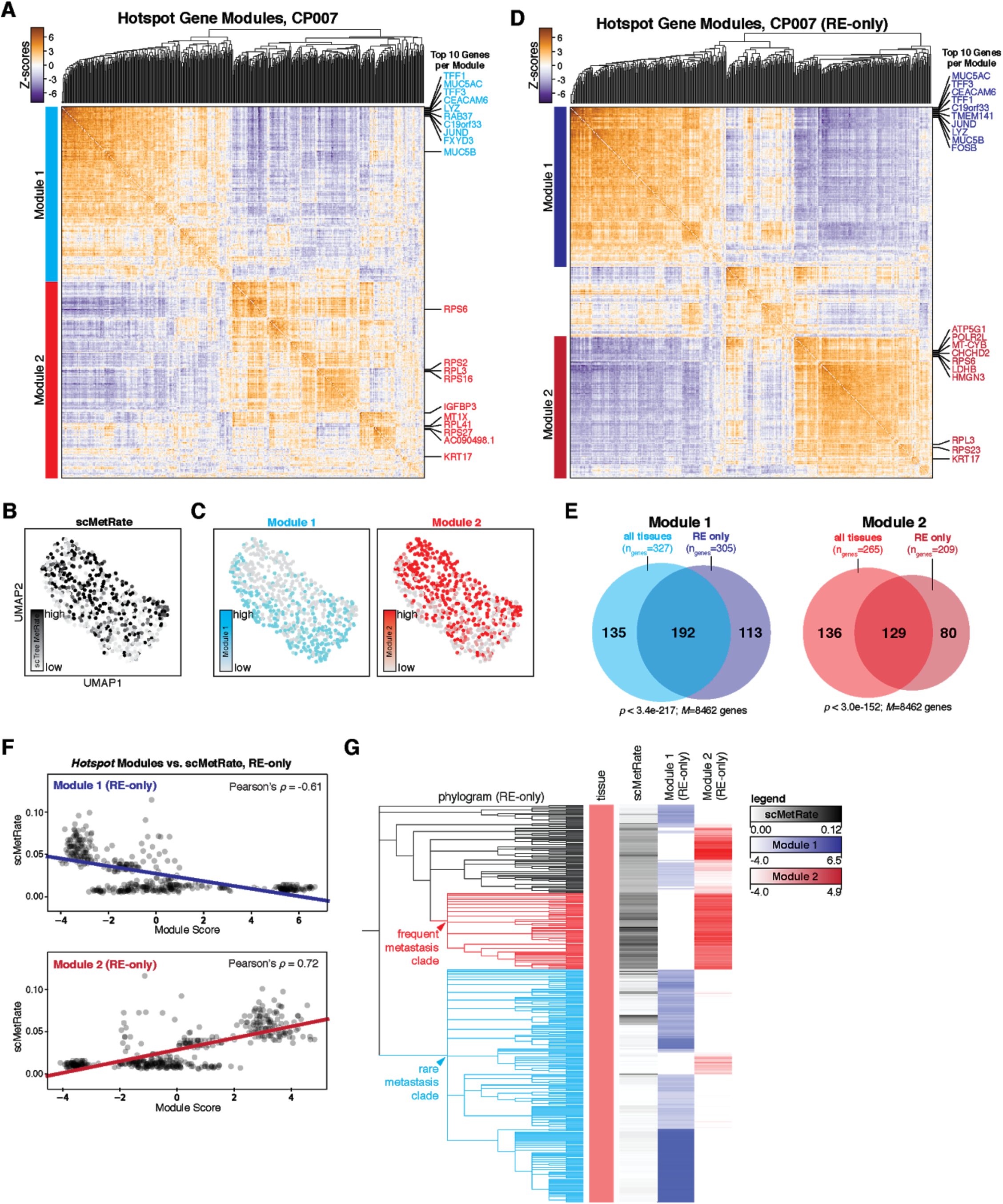
Distinct transcriptional modules underlie distinct clade-specific metastatic behaviors in Clone #7. (**A**) *Hotspot* analysis identifies two gene modules that have heritable expression patterns in CP007 (Modules 1 and 2; indicated in cyan and red, respectively). Pairwise local correlations of genes (with FDR < 0.1) calculated with *Hostpot* are shown by heat-map; the top 10 most significant genes each for Modules 1 and 2 are annotated in cyan and red, respectively. (**B** and **C**) The transcriptional states for all cells in CP007 represented in a two-dimensional projection (UMAP) and colored by scMetRate (**B**) or *Hotspot* Module scores (**C**). (**D**) When restricting *Hotspot* analysis to only cells from the “RE” tissue sample in CP007, two heritable gene modules are identified (dark blue and dark red). As before, the top 10 genes from each module are annotated. (**E**) The gene modules identified from all cells or from only RE-only cells in CP007 overlap significantly by hypergeometric test. Number of genes in each set (*n*_*genes*_), number of expressed genes in at least 10% of cells in CP007 (*M*), and *p*-value of the hypergeometric test are indicated. (**F**) RE-only Modules 1 and 2 are negatively and positively associated with the scMetRate, respectively, as in the analysis for all cells from CP007 (**Fig. 5I**). (**G**) Overlay of the phylogram for RE-only cells from CP007, scMetRate, and *Hotspot* module scores (RE-only), showing concordance between the frequently metastasizing clade (red) and Module 2 and the rarely metastasizing clade (cyan) and Module 1, as in **Fig. 5J**.

**Fig. S22.**
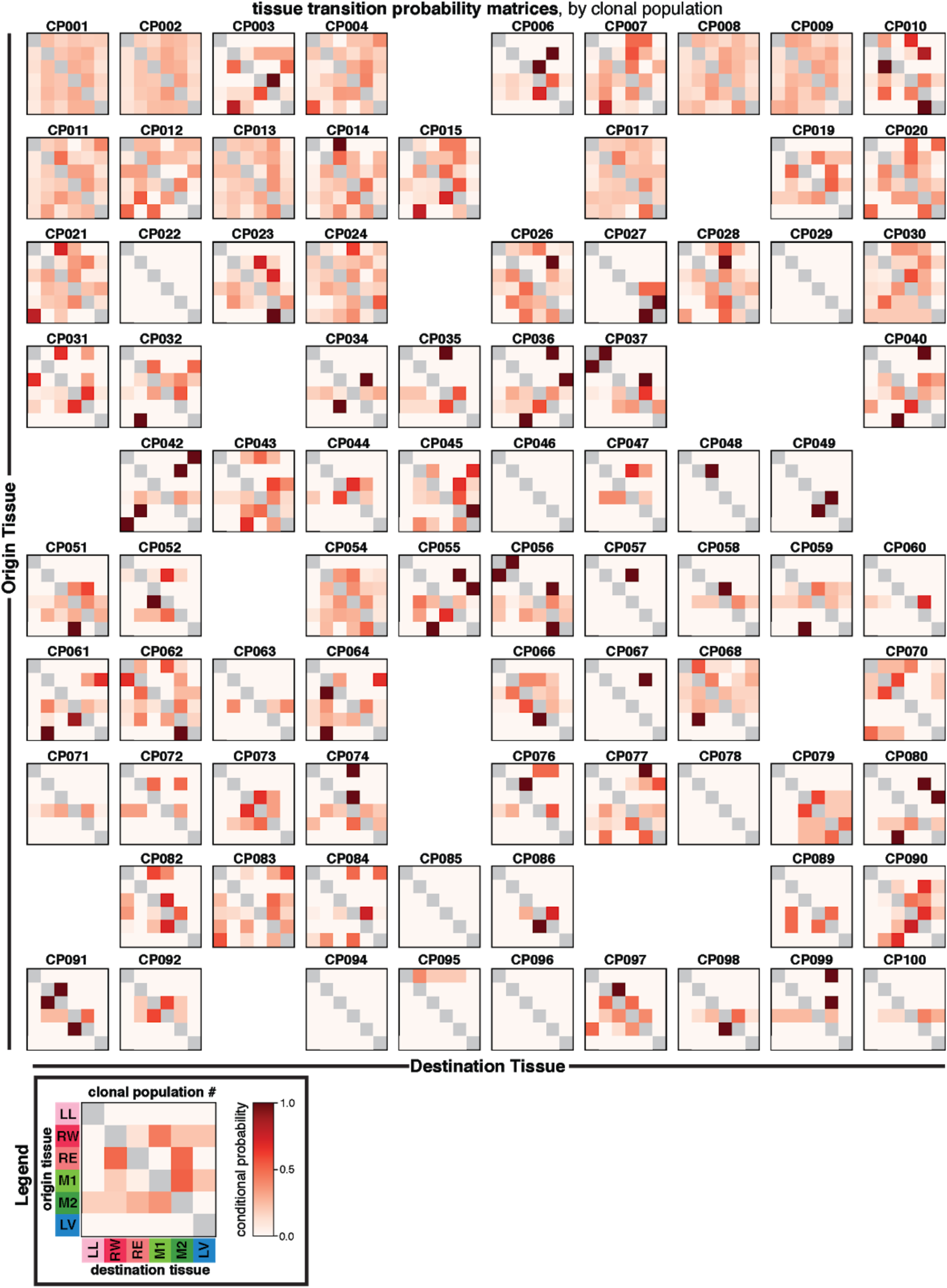
Tissue transition probability matrices for each clonal population. The conditional probability of transition from and to each tissue inferred from the phylogenetic trees (calculated with *FitchCount)* of each clonal population, thus summarizing the most probable tissue routes of metastasis. Legend (lower left) indicates the color bar showing conditional probability and the tissue labels, as in **Fig. 1E**. Notably, the transition matrices are varied and distinct to each clonal population.

**Fig. S23.**
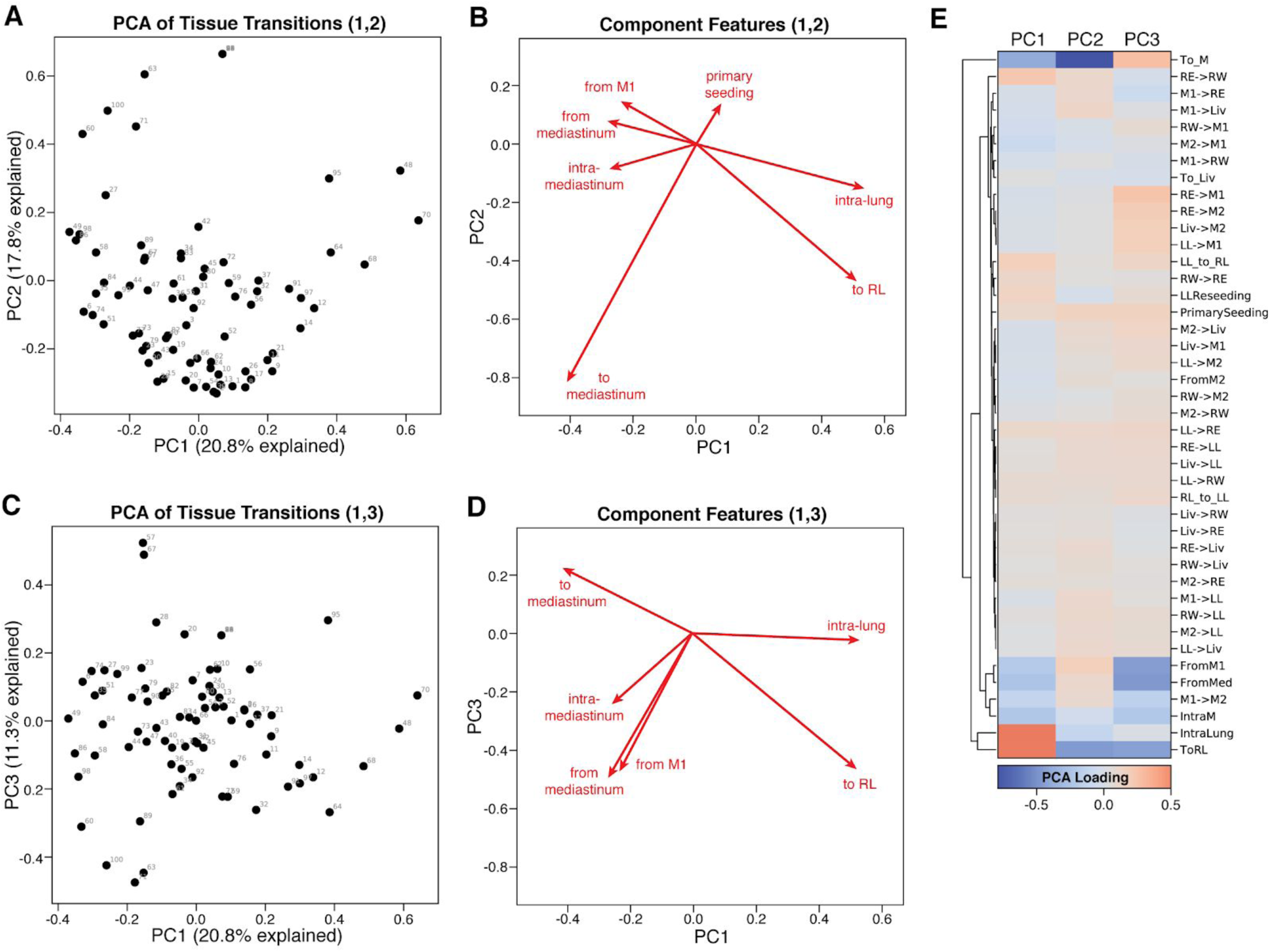
Describing the principal features of metastatic seeding routes. (**A, C**) PCA projections of the metastatic tissue transitions for each clonal population (annotated). The percentage of the variance explained by each component is indicated on the axes for the first and second (**A**) or first and third (**C**) components. (**B, D**) Biplot vectors representing the most explanatory features of the first, second, and third principal components, annotated by descriptive features of metastatic transitions. The length and angle of the vector describe the scale and direction, respectively, of each descriptive feature. (**F**) The PCA loadings of the metastatic transition features for each principal component.

**Fig. S24.**
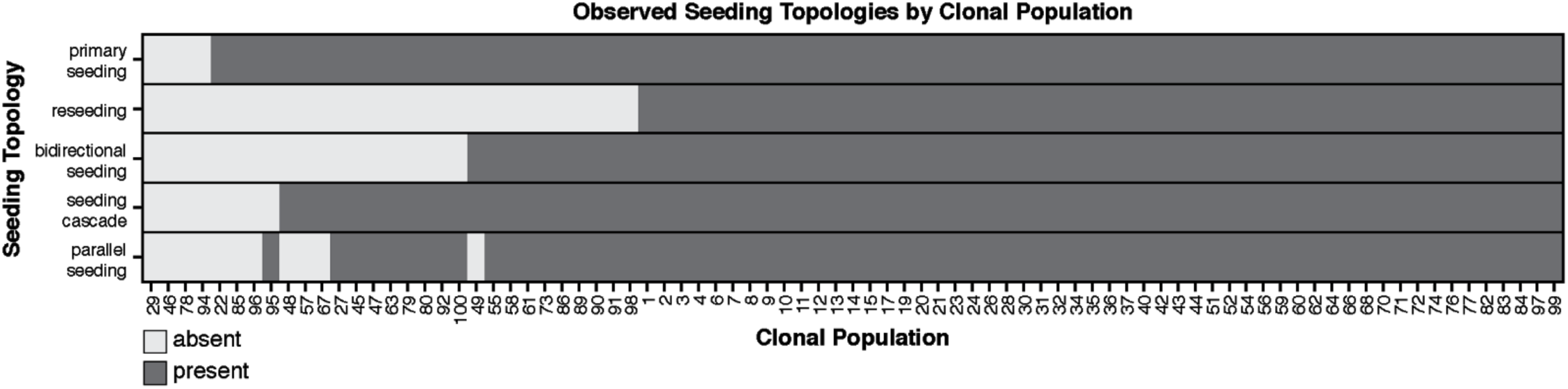
Seeding topologies observed in each clonal population. A table describing classified seeding topologies (rows) that are present or absent (dark or light gray, respectively) in each clonal population (columns). The majority of clonal populations exhibit examples of all seeding topologies.

### Supplementary Text

#### *FitchCount:* an Efficient Algorithm for Inferring Transition Matrices on Phylogenetic Trees

##### 1 Deriving transition matrices from phylogenetic trees

In this document, we describe our approach for solving the following problem: Consider a phylogenetic tree 𝒯 rooted at vertex r over V vertices and E edges which we denote as 𝒯^*r*^. In this tree, each leaf l is assigned a state state(l), where states are drawn from a state space Σ (i.e. ∀v, state(v) ∈ Σ). Here, leaves are single cells derived from a single-cell lineage tracing experiment (Chan et al. 2019) studying cancer metastasis, the tree describes the cells’ evolutionary history (as inferred by Cassiopeia (Jones et al. 2020)), and the states represent the tissues from which the cells were obtained. Our task is to derive summary statistics obtained by assigning states to the internal nodes of the tree(i.e., ancestral cells that were not observed in the study). Specifically, the summary statistics we are interested at are: (1) the overall number of metastatic events (i.e., transition between tissues) that occurred in a clone’s history and (2) the frequency (i.e. number) of transitions between each pair of tissues s_*i*_ and s_*j*_.

This class of problems typically requires what is known as “ancestral state reconstruction” (Joy et al. 2016; Slatkin and Maddison 1989; McPherson et al. 2016), which in essence attempts to assign an ancestral states to every node in a given tree that minimizes some function. Here, we use parsimony as our objective function. Our work relies on classical algorithms (Fitch 1971; Hartigan 1973) and is also inspired by recent algorithms using these principles to infer metastatic histories like MACHINA (El-Kebir, Satas, and Raphael 2018).

As noted by El-Kebir et al (El-Kebir, Satas, and Raphael 2018), while there exist several algorithms for effectively inferring clone trees from DNA samples of metastatic cancers (Reiter et al. 2017; Deshwar et al. 2015; El-Kebir et al. 2015), none of them except for MACHINA propose an ancestral node labeling and subsequent classification of metastatic topologies. This is because a metastatic history does not follow uniquely from a tree structure, and metastasis itself is not necessarily unidirectional (i.e. there exist polyclonal & reseeding events that may introduce cyclic topologies). Though MACHINA represents a significant advance, we build on it by reporting summary statistics over *all* optimal solutions rather than one, and additionally circumvent the computationally-intensive Integer Linear Programming (ILP) optimization routine in favor for a dynamic programming approach.

Below, we describe our algorithmic strategy (and prove it) for inferring both summary statistics. We begin by describing how the overall number of transitions can be derived from the Fitch-Hartigan algorithm (Fitch 1971; Hartigan 1973). Next, we introduce *FitchCount*, an algorithm for counting the number of transitions between any two tissues in a given phylogeny over all optimal solutions to the Fitch-Hartigan algorithm.

#### 2 Algorithmic strategy

The first summary statistic that we are interested ind is the minimal possible number of state transitions in the tree that is sufficient to explain the state assignment to the leaves (which is given as an input). In other words, out of all possible assignment of tissue labels to the ancestral cells (which can be exponentially many), consider the assignments that entail the minimum number of cases in which a parent node and a child node come from a different tissue (i.e., state transition). Our first goal is to retrieve the number of state transitions in these optimal assignments, but not the assignments themselves (note that the number of transition is the same [i.e., minimum possible] in all optimal assignments). In our second goal, we are interested in the number of specific transitions across all optimal assignments, which means that we would have to also look at the optimal assignments themselves.

While our first goal can be readily addressed by the classical algorithm of Fitch (Fitch 1971) and Hartigan (Hartigan 1973) or using another algorithm by Sankoff (Sankoff 1975), the second goal requires an additional procedure. The reason for this is that the existing algorithms are able to retrieve only one specific optimal assignment in each run through the tree. However, we would ideally like to base our summary statistics on the space of all possible optimal assignments. Since there can be exponentially many optimal assignments, we needed to find an efficient way to extract the summary statistic without actually enumerating all algorithms.

Notably, the Fitch (Fitch 1971) algorithm was originally designed for binary trees. An important property of this algorithm is that the optimal assignments that it produces guarantee optimality even if we consider every sub-tree in isolation. This is different from a scenario where we allow state assignments that may not be optimal when we are considering only a certain sub-tree but become optimal due to compensation elsewhere in the tree. The Hartigan algorithm extends it to non-binary trees and can also be modified such that its optimal assignments remain optimal in every sub-tree. We refer to this modification as the Fitch-Hartigan algorithm. This algorithm operates in a time linear in the input size (i.e., it scales proportionally to *n · k* where *n* is the number of cells and *k* is the number of possible states [tissues, in our case]). The Sankoff algorithm (Sankoff 1975) uses a more involved formulation with a slower run time (it scales proportionally to *n · k*^2^) that can account for different penalties to different state transitions. It also returns all possible optimal solutions, including solutions that may not be optimal for every sub-tree, if it is considered in isolation. Here, we employ the Fitch-Hartigan approach due to its simplicity and speed and since we reasoned that it is desirable that our solutions remain optimal, not just for the entire clone, but also for every sub-clone individually.

#### 3 Finding the minimal number of transitions

The Fitch-Hartigan algorithm begins with a “bottom-up” procedure in which labels at the leaves are propagated up to internal nodes in the tree. This “bottom-up” phase assigns a set of labels (tissues) *opt*[*v*] to each node v in the tree that satisfy the optimality demands (namely, maximum parsimony). Specifically, for every s ∈ *opt*[*v*] there exists at least one state assignment to the nodes in *T* ^*v*^ (the tree rooted by *v*) where the state of *v* is s and that is optimal for *T* ^*v*^ and for every sub-tree of *T* ^*v*^. Furthermore, the set opt[v] includes all such states. For completeness, we provide a proof for these claims in the appendix (Claim 3).

In addition to generating the sets opt the algorithm can also count for every node v the minimum possible number of state transitions *n*(*v*) required in the sub-tree rooted by *v* (Observe that *n*(*r*) for the tree rooted at r corresponds to the number of transitions across the entire tree). The “bottom-up” procedure *opt* is applied in **post-order traversal** (evoked by applying it on the root node) with the following pseudocode:

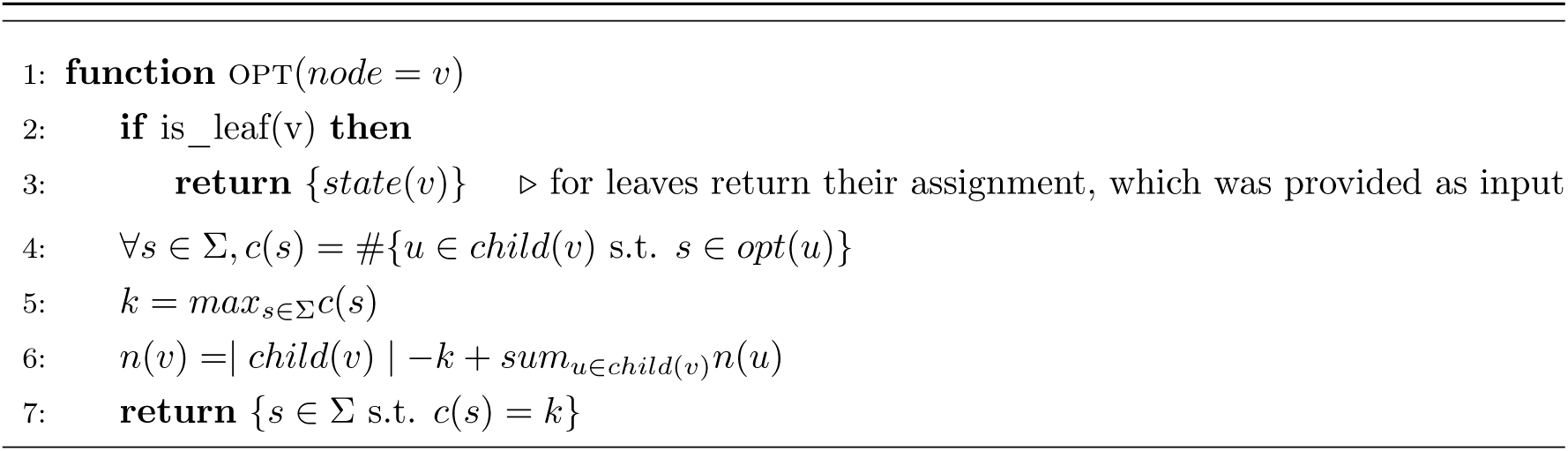

Note that in the original formulation by Hartigan, an additional complication is added to increase the number of optimal assignments that can be retrieved by the algorithm. This is done by accumulation of an additional set of states *opt*_*2*_(*v*) for every node v that can be used to derive solutions that are globally optimal, but are not optimal in the sub-tree rooted by v. We therefore excluded this part of the algorithm (see appendix for proof [Claim 3]).

#### 4 Inferring the frequency of different state transition events

The second part of the Fitch-Hartigan algorithm is a top-down procedure for finding one (out of potentially many) optimal solution, i.e., a labeling state : V → Σ of each node *v* ∈ *V* in the tree with a state *s* ∈ Σ. It starts by randomly selecting a state for the root node r out of the set *opt*(*r*) and then continues to select legal (see Definition 1) states for child nodes, based on the value assigned to their parent.

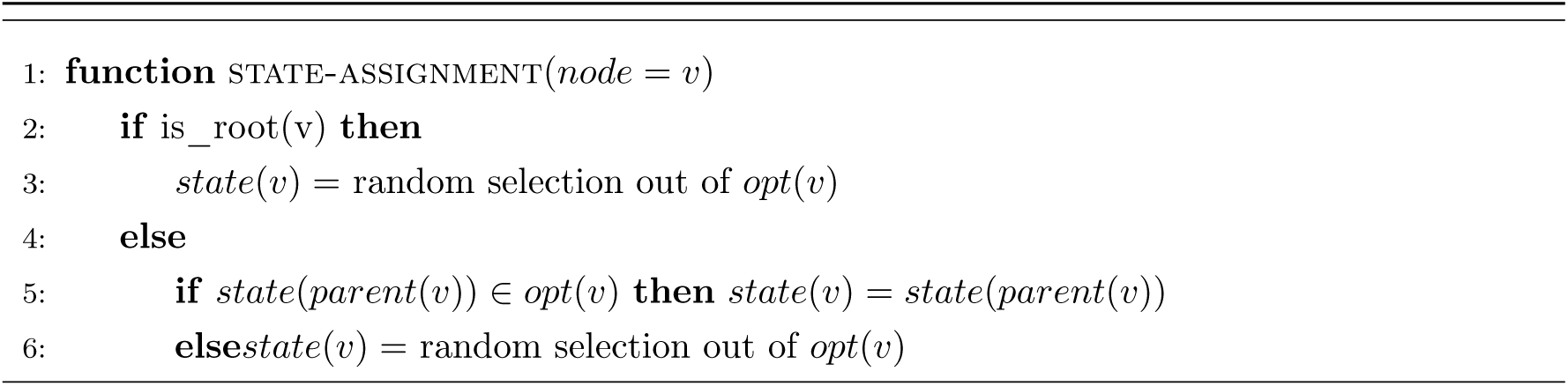

The above procedure can face many ties during its execution, and can thus potentially return all optimal solutions (provided that they remain optimal for every sub-tree) if it is applied many times. When we compare Fitch-Hartigan to FitchCount in the main text, we use one top-down round (i.e., consider one optimal solution) for the former.

The *FitchCount* procedure was designed to provide a comprehensive evaluation of state transition frequencies, by basing its estimation on the space of all possible optimal state assignments, instead of only a single or few optimal assignments. Compared to the naive approach to enumerate all possible optimal state assignments given by the Fitch-Hartigan algorithm (which may require exponential number of executions), *FitchCount* performs in 𝒪 (*n · k*^3^) time for a tree with n leaf nodes and k possible states (assuming each internal node has at least two child nodes, which is the case in our work).

##### 4.1 The algorithm

We define several arrays for storing necessary information:

1. *opt*[*v*]: The set of optimal assignments for a node v given by the Fitch-Hartigan bottom up approach procedure (defined in the algorithm *opt*).
2. *N*[*v, s*]: The number of optimal solutions below the node v given that it takes on the state *s*.
3. *C*[*v, s, s*_*i*_, *s*_*j*_]: The number of transitions from state *s*_*i*_ to state *s*_*j*_ in all optimal solutions of the tree rooted at *v*, given that *v* takes on the state *s*.
4. *M*[*i, j*]: The number of transitions between *s*_*i*_ and *s*_*j*_ observed across all optimal solutions to the Fitch-Hartigan algorithm.

Our overall objective is to fill in the dynamic programming matrix *M*, which will subsequently require knowledge of the other dynamic programming arrays. Note that in the following we refer to the arrays using either rounded parenthesis or rectangular parenthesis. The former denotes a function call and the latter denotes a retrieval of an already-computed entry (which we assume to get populated automatically after the respective function call and available globally to the algorithms). Our Main function is:

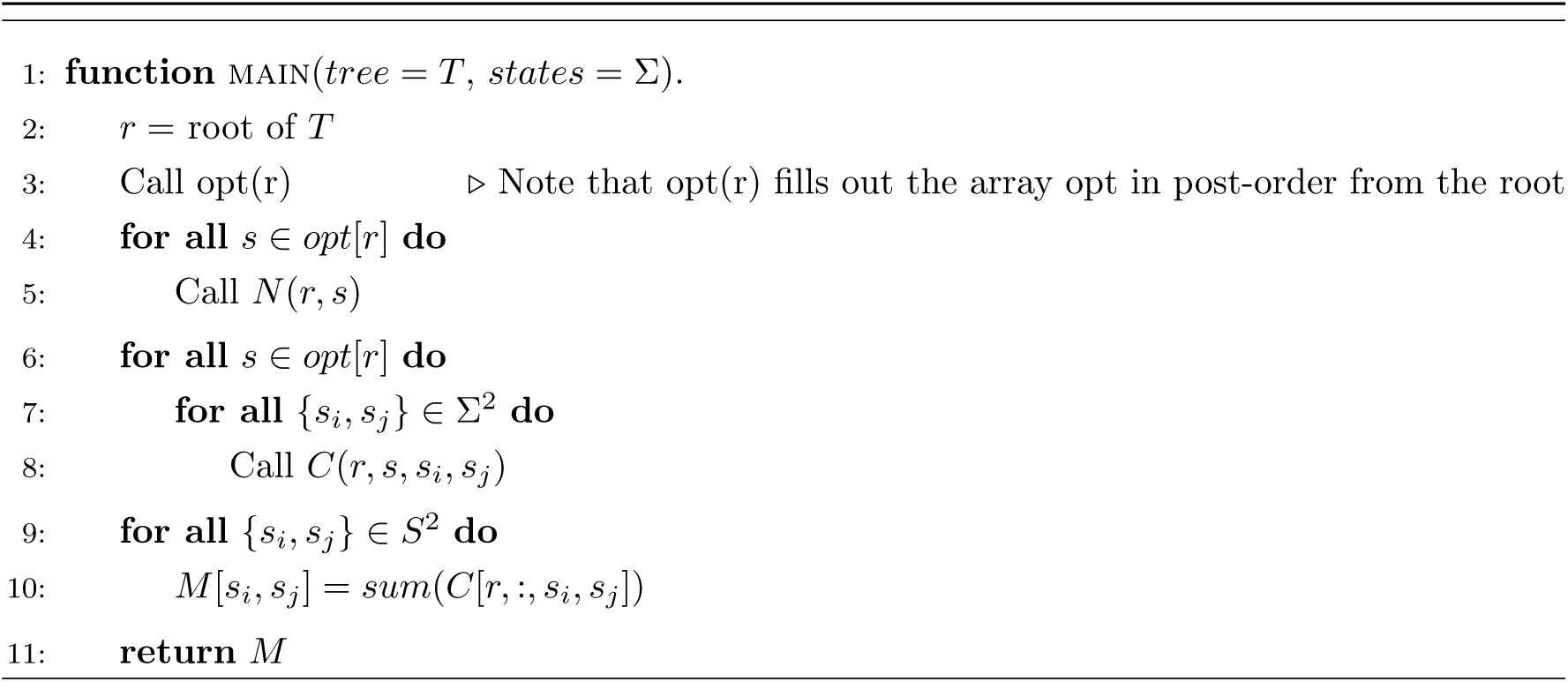

The following algorithms for filling in specific entries to *N*[*v, s*], and *C*[*v, s, s*_*i*_, *s*_*j*_]:

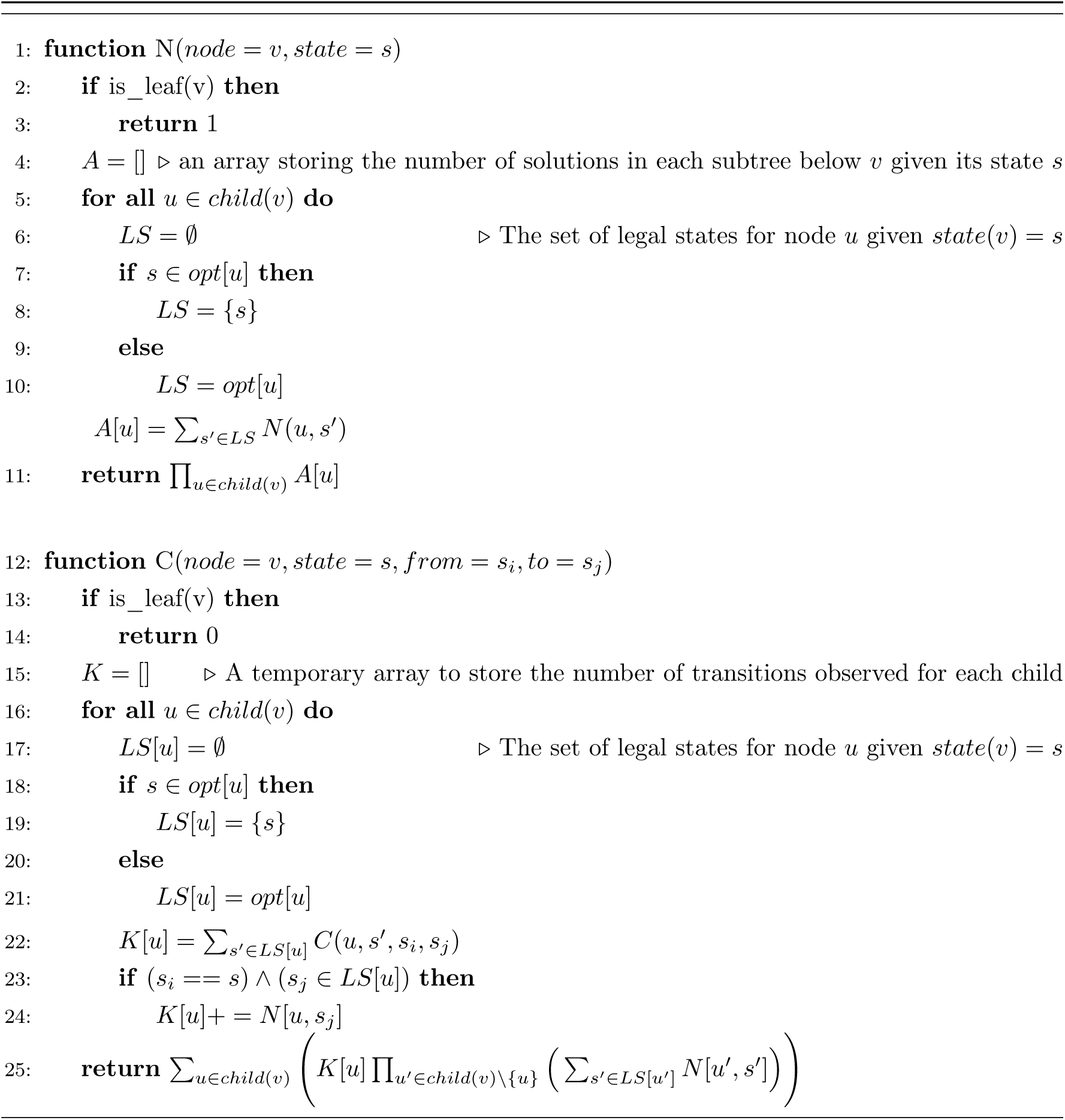

##### 4.2 Proof

We prove correctness of the dynamic programming matrices N, and C.

**Claim 1**. *For any node* v *and state* s ∈ *opt*[*v*], *N*[*v, s*], *is precisely the number of optimal solutions in T* ^(*v*)^ *where* state(*v*) = *s*.

*Proof*. To prove the correctness of the dynamic programming array N, we’ll proceed by induction over the height of the tree *height*(*T*) = *h*. For convenience, we’ll also make use of a temporary dynamic programming array A which stores the number of solutions for each child *c* ∈ *child*(*v*), aggregated across each possible state of that child given the parent’s state *s*.

###### Base Case #1, h = 0

*N*[*l, state*(*l*)] = 1. This relation is trivially true for the case where *h* = 0 and there exists a single leaf l_1_ in which case the only solution consists of state(*l*) = *s*.

###### Base Case #2, h = 1

Consider a tree *T* ^(*v*)^ rooted at node v, where *child*(*v*) = *{l*_1_, …, *l*_*m*_*}*. We know that each leaf l_*i*_ has a single assignment state(*l*_*i*_) from the definition of the small-parsimony problem. Thus, the number of possible solutions for this tree with *state*(*v*) = s is always one, namely with each leaf taking on their only state. Specifically, in this base case, we observe that *A*[*u*]= 1 ∀ *u* ∈ *{l*_1_, …, *l*_*m*_*}*. To show this, we consider two cases:

- If s ∉ *opt*(*l*_*i*_): 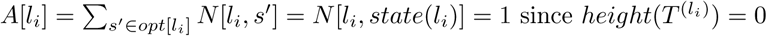.
- If *s* ∈ *opt*[*l*_*i*_]: *s* == *stat*e(*l*_*i*_) and *A*[*l*_*i*_]= *N*[*l*_*i*_, *s*]= 1 *since height*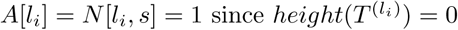. and the relation

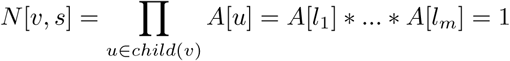

thus is correct.

###### Inductive Hypothesis

For a tree *T* ^(*v*)^ of height *h*, and some *s* ∈ *opt*[*v*], *N*[*v, s*] exactly stores the number of optimal solutions in the tree rooted at *v*.

###### Inductive Step

Consider a tree *T* ^(*v*)^ of height *h* + 1 and some state s ∈ *opt*[*v*]. We will show that both the array *A* correctly stores the number of solutions for the child *u* given *state*(*v*) = *s* and that the relation *N*[*v, s*]= _*u∈child*(*v*)_ *A*[*u*] is correct.

First, we note that for the tree to be globally optimal, for each *u* ∈ *child*(*v*), *state*(*u*) must be *s* if *s* ∈*opt*[*u*]; else, any state from *opt*[*u*] can be assigned to u as each incurs a cost of 1 to the overall parsimony of the tree (see Claim 3). These choices for “optimal” states are stored in the array *LS*[*u*].

Second, we know from our inductive hypothesis that *N*[*u, s*^*0*^] is correct for any child *u* ∈ *child*(*v*) and any state *s*^*0*^ ∈ *opt*[*u*] as the tree *T* ^(*u*)^ has a height h. Thus, it is clear that 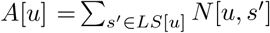 correctly returns the number of solutions in the subtree rooted at *u* over all possible legal states that u can take on.

Finally, we observe that given *state*(*v*) = *s*, each child can be treated independently as we consider global solutions that in the tree *T* ^(*v*)^ with *state*(*v*) = *s*. Because of this, the number of such solutions is the size of the permutation of all optimal sub-trees rooted at each *u* ∈ *child*(*v*) - i.e. the product of all *A*[*u*]. To show this, consider *v* has m children. Let the set of optimal internal labellings to 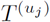 be denoted as 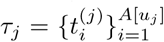 where 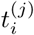 is the i^*th*^ solution for the tree rooted at *u*_*j*_ given *state*(*v*) = *s*. Then, the possible set of solutions is the Cartesian Product between *τ*_1_, *τ*_2_, …, *τ*_*m*_:

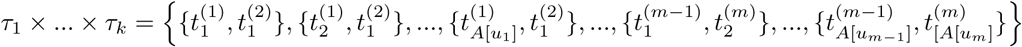

This Cartesian product has a size |*τ*_1_| × |*τ*_2_|… × |*τ*_*m*_| = *A*[*u*_*i*_] × … × *A*[*u*_*k*_]= _*u*∈*child*(*v*)_ *A*[*u*]. Thus, this relation holds T ^(*v*)^ where height(*T* ^(*v*)^) = *h* + 1.

**Claim 2**. *For any node* v *and state s* ∈ *opt*[*v*] *assigned to* v *and {s*_*i*_, *s*_*j*_*}* ∈ 2^Σ^, *the array C*[*v, s, s*_*i*_, *s*_*j*_] *correctly stores the number of transitions from* s_*i*_ → s_*j*_ *in T* ^(*v*)^.

*Proof*. We will prove by induction over the height of the tree, h, that for a node *v*, a state *state*(*v*) = *s*, and some (s_*i*_, s_*j*_) ∈ 2^Σ^ both *K*[*u*] ∀*u* ∈ *child*(*v*) and

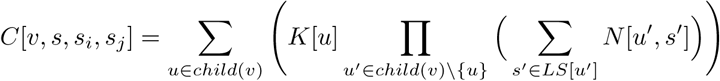

are correct. Here *K*[*u*] is the number of transitions from *s*_*i*_ to *s*_*j*_ that exist by considering child *u* of node *v* given *state*(*v*) = *s* and *LS*[*u*] is a function that finds the set of legal (Definition 1) assignments to *u* given the parent’s state is *s*.

###### Base Case #1,*h*= 0

The relation trivially holds for a tree of height 0 as there cannot exist any transitions for a tree without edges. As calculated, *C*[*v, s, s*_*i*_, *s*_*j*_] = 0 for all leaves and thus the relation holds.

###### Base Case #2,*h*= 1

Consider a tree of height 1, *T* ^(*v*)^, where child(*v*) = *{l*_1_, …, *l*_*m*_*}* and that *N*[*l*_*i*_, *state*(*l*_*i*_)] = 1. We can count the number of transitions by considering for every edge the following:

- s ≠ = *s*_*i*_ then *C*[*v, s, s*_*i*_, *s*_*j*_] is necessarily 0.
- *s* == *s*_*i*_, then *C*[*v, s, s*_*i*_, *s*_*j*_] is the number of leaves that have state *s*_*j*_.

By construction, for some child *l*_*i*_, *C*[*v, s, s*_*i*_, *s*_*j*_] must be

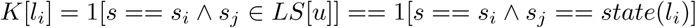

Then *C*[*v, s, s*_*i*_, *s*_*j*_]_*l*∈*child*(*v*)_ *K*[*l*_*i*_]. We’ll prove that this is equal to the relation described above

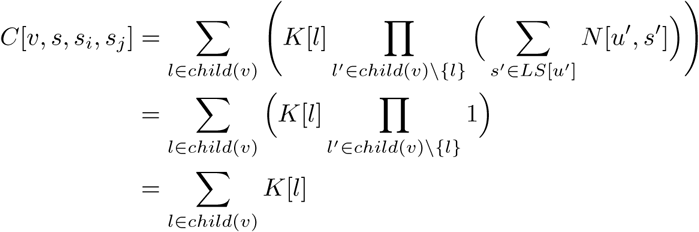

Thus, our relation holds for *h* = 1.

###### Inductive Hypothesis

Assume for a tree *T* ^(*v*)^ of height h where *state*(*v*) = *s, C*[*v, s, s*_*i*_, *s*_*j*_] correctly computes the number of transitions from s_*i*_, s_*j*_ ∈ 2 ^Σ^ for the tree.

###### Induction Step

Now consider a tree of height *h* +1 rooted at v where *state*(*v*) = *s*. We’ll show that both *K*[*u*] is correct for all *u* ∈ *child*(*v*) and that the relation for calculating *C*[*v, s, s*_*i*_, *s*_*j*_] holds.

We’ll first show that *K*[*u*] is correct ∀*u* ∈ *child(v)*. As defined above, *K*[*u*] is the number of s_*i*_ → s_*j*_ transitions that are due to the node *u* given *state*(*v*) = *s*. We know that given our inductive hypothesis, for the subtree rooted at u, *T* ^(*u*)^, C[*u, s*^*0*^, *s*_*i*_, *s*_*j*_] for any state s^*0*^ ∈ *opt*[*u*] is correct. Then, the number of *s*_*i*_ → *s*_*j*_ transitions under *u*, given *state*(*v*) = *s*, is equal to the sum of all *C*[*u, s*^*0*^, *s*_*i*_, *s*_*j*_] for those *s*^*0*^ ∈ *LS*[*u*] as those are the only solutions that would be considered by the Fitch-Hartigan algorithm (note that we are guaranteed to have optimal state assignments to chose from for LS[u] as we prove in Claim 3), plus the transition (if it exists) from *v* to *u*.

Now, we’ll show that the relation for *C*[*v, s, s*_*i*_, *s*_*j*_] holds. For the tree *T* ^(*v*)^ let’s assume that v has m children: 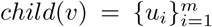. As above, we’ll maintain the notation that the set of legal assignments given that *state*(*v*) = *s* for *u*_*i*_ to be *LS*[*u*_*i*_]. Furthermore, let the set of *s*_*i*_ → *s*_*j*_ transitions underneath *u*_*j*_ be 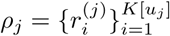 and the set of trees that are legal, optimal assignments under *u*_*J*_ be 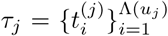 where 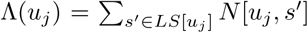, assuming state(*v*) = *s*. We can see then that the total number of transitions from s_*i*_ → s_*j*_ is

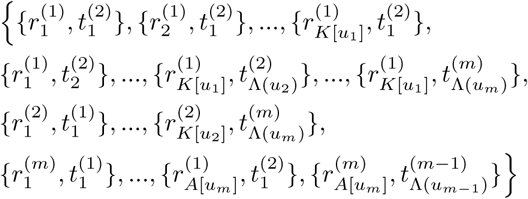

Which is equal to the sum of the following Cartesian Products:

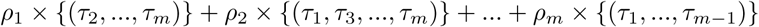

where the cardinality of this set is

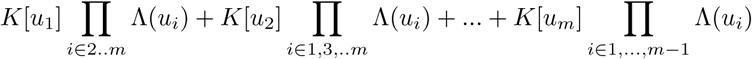

which can be further simplified to

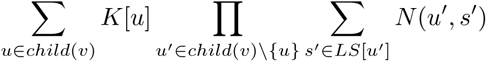

Thus the relation is correct and *C*[*v, s, s*_*i*_, *s*_*j*_] is correct by induction.

#### 5 Appendix

##### Definition 1.

*(Legal Assignment). An assignment state(v) = s is* ***legal*** *for a node v and given state(parent(v)) = s*^*0*^ *if either s == s*^*0*^ *or s*^*0*^ ∈*/ opt(v). Observe that only legal assignments are explored in the Fitch-Hartigan algorithm, and are guaranteed to be optimal in the sub-tree rooted at v*.

**Claim 3**. *Consider any node v and let T (v) denote the sub-tree rooted at v. The bottom up procedure above returns a set opt(v) such that for every s* ∈; *opt(v): (1) there exists a solution (state assignment) that is optimal for T (v) and every sub-tree of T (v), in which the state of v is s; and (2) there does not exist a solution that is optimal for T (v) and for every sub-tree of T (v), in which the state of v is some s*^*0*^ ∉ *opt(v)*

*Proof*. Proof by induction on tree height h (max length from root to any leaf).

##### Base Case #1, *h* = 0

In this case the tree consists of a single leaf node, for which state assignment is already fixed and the claim follows trivially.

##### Base Case #2, *h* = 1

In this case, we have one internal node v with n leaves as its immediate descendants. Here, we define *opt*(*v*) as the set of all states that are found in k out of n descendants, where k is maximal. Clearly, the value of the optimal solution in this case has n *-* k state transitions, which can be obtained by assignment of any state in opt(v) to v. Furthermore, the solution is trivially optimal for every sub-tree (i.e., singleton), thus proving the first part of the claim. Furthermore, assignment of any s^*0*^ ∉ *opt(v)* to v will necessarily entail strictly more than *n - k* state transitions, thus proving the second part of the claim.

##### Inductive step

Assume an internal node v whose corresponding sub-tree is of height h. Let *C* denote the set of its child nodes. From the induction, we assume that the claim holds for every node *u* ∈ *C. opt(v)* is defined as the set of all states that are found in k out of the m =| C | child nodes, where k is maximal. First let us denote by optval(v) the value (number of state transitions) of the optimal solution for T (v). From the assumption of the induction, it is easy to see that optval(v) = sum_*u2C*_optval(u)+ m *-* k. This value can be reached following the top-down procedure of the Fitch-Hartigan algorithm: (i) assign v with some state s *2* opt(v); (ii) assign s to all child nodes u where s *2* opt(u) (iii) assign each remaining child node u^*0*^ with some other state from opt(u^*0*^) (iv) consider some optimal solution for each of the sub-trees that are rooted by the child nodes. Note that these optimal solutions must exist due to the assumption of our induction. Clearly, the solution that we built satisfies the first part of our claim. For the second part of our claim, assume by contradiction that there exists a state assignment in which the state of v is s^*0*^ ∉*opt*(*v*) that achieves optimality for T (v) and all of its sub-trees. However, from the assumption of the induction we must choose an assignment for every child u out of its set *opt(u)*. It therefore follows that the value of any such solution must be at least sum_*u2C*_optval(*u*)+ *m - k* + 1. We note that in the original paper by Hartigan, a solution is possible where for one or more child nodes u we select an assignment that is identical to the state of the parent v but is not from *opt(u)*. However, while this solution reaches optimality for *T* (*v*), it will not be optimal for *T* (*u*).

