## Supplementary Materials for "Single-cell lineages reveal the rates, routes, and drivers of metastasis in cancer xenografts"

$$h'_p(a_i,b_i) = \begin{cases} -log(p(a_i)) - log(p(b_i)) & \text{if } a_i \neq b_i \text{ and } a_i, b_i \text{ are mutated} \\ -log(p(a_i)) & \text{if } a_i \text{ mutated, } b_i \text{unmutated} \\ -log(p(b_i)) & \text{if } b_i \text{ mutated, } a_i \text{unmutated} \\ log(p(a_i)) + log(p(b_i)) & \text{if } a_i, b_i \text{ mutated and } a_i == b_i \\ 0 & o.w. \end{cases}$$

Finally, distances $d'(a,b)$were normalized by the number of cut-sites overlapping between cells $a$ and $b$.

***Allelic vs. phylogenetic distance calculations.*** The concordance between allelic and phylogenetic distances were used to assess the agreement between a tree's phylogenetic structure and the observed mutations. To quantify allelic distances, we defined a modified Hamming distance between cells *a* and *b*,

$$d(a,b) = \sum_{i \in M} h'(a_i, b_i)$$

Where $M$ is the number of cut-sites in the clonal population that cells $a$ and $b$ come from, $a_i$ and $b_i$are the states observed at the $i^{th}$character, and *h'* is defined as follows:

$$h'(a_i, b_i) = \begin{cases} 2 & \text{if } a_i \neq b_i \text{ and } a_i \neq 0 \text{ and } b_i \neq 0 \\ 1 & \text{if } (a_i == 0 \text{ or } b_i == 0) \text{ and } a_i \neq b_i \\ 0 & \text{o.w.} \end{cases}$$

Importantly, these values were normalized by $2 * M$, the maximum distance for a pair of cells.

The phylogenetic distance was calculated for all pairs of cells as the number of non-zero length edges that separated the two cells in the tree. The phylogenetic distances were normalized by the "diameter" of the phylogeny, i.e., the maximum distance between any pair of cells.

***Derivation of the AlleleMetRate.*** The AlleleMetRate is provided as a "tree-agnostic" measurement of a clonal population's intrinsic metastatic potential. Intuitively, the rate measures the proportion of cells that do not reside in the same tissue as their closest relative (as determined by the modified Hamming distance between two cells' character-states). Importantly, if a cell has more than one closest relative, each of their votes are normalized by the number of relatives this cell has. More formally, the contribution a cell has to the AlleleMetRate is

$$m(c) = \frac{1}{K} \sum_{i \in Neighbors(c)} I(tissue(i) \neq tissue(c))$$

Where *K* is the number of closest relatives a cell has, and *I(*)* is an indicator function that equals 1 if the tissue of cell *i* is different from the tissue of cell *c*. The AlleleMetRate reported is

$$M_a = \frac{1}{|L|} \sum_{l \in L} m(l)$$

Where *L* is the set of all leaves in the tree.

***Derivation of the TreeMetRate.*** The TreeMetRate is derived from the Fitch-Hartigan maximum parsimony algorithm (Fitch, 1971; Hartigan, 1973). Briefly, the Fitch-Hartigan algorithm takes in a rooted tree with labelings at the leaves (in our case labels indicate which tissue the cell was obtained from) and in our case reports two items: (a) a distribution of labels at the internal nodes that minimize the number of transitions between tissues (i.e., achieves maximum parsimony); and (b) the minimum number of transitions between tissues (referred to as the "parsimony score"). The Fitch-Hartigan algorithm is an efficient (scaling with $n \cdot k$where *n* is the number of cells and *k* is the number of tissues) and common algorithm for solving the "Small Parsimony Problem", as opposed to the "Large Parsimony Problem" which attempts to find the *tree* of maximum parsimony.

The Fitch-Hartigan algorithm operates in two phases: first, by propagating up from the leaves the set of possible labelings at each internal node; second, by traversing in depth-first order from the root of the phylogeny and selecting one state for each internal node from its set of optimal labels. Conveniently, the parsimony score can be obtained from this second phase. To finally report the TreeMetRate, we simply normalize this parsimony score by the number of edges in the tree.

***Derivation of the single-cell MetRate***. The scMetRate for a given cell is defined as the average of all TreeMetRates for the clades that contain that cell. Specifically, we employ the following algorithm:

```
compute_scMetRate(tree, x):
     rate = 0
     num_ancestors = 0
     for n in depth_first_search(tree, x):
          rate += compute_tree_metrate(n)
```

```
            num_ancestors += 1

        rate /= num_ancestors
        return rate
```

Where *compute_tree_metrate(n)* is a function that computes the TreeMetRate for the subtree rooted at an internal node *n* (as described above).

***scRNA-seq preprocessing and normalization.*** RNA-seq libraries were quantified using CellRanger version 2.1.1 with the GRCh38 genome build for M5k and M30k; and CellRanger version 2.0.0 with the hg19 genome build for M10k and M100k. Cells not found in the Target Site Library were filtered out. All RNA-seq datasets were normalized identically – we first normalized the UMI counts in every cell to the median number of UMIs found in each library (referred to hereafter as "UMI-normalized" counts) and then each matrix of UMI-normalized counts was log-transformed (after adding a 1-pseudocount to preserve 0's in the data; hereafter referred to as "log-transformed" counts). In the analysis presented in Figure 5A we applied scVI (version 0.6.5) to the raw counts of the M5k and the pre-implantation cells to obtain a shared, batch-corrected latent space (learning rate of 1e-3, 400 epochs, 10 latent dimensions). This space was used for downstream projections and *Vision* analyses.

$$\widehat{mean}(TMR) = \frac{1}{B}\sum \quad compute\_tree\_metrate(tree)$$

$$\widehat{se}(TMR) = \frac{1}{B}\sum \quad (compute\_tree\_metrate(tree) - \widehat{mean}(TMR))$$

where $B$ is the number of bootstrapped samples for a given metastatic process (here, $B = 100$). The coefficient variation for each metastatic process was defined as the ratio $\widehat{se}/\widehat{mean}$.

**Supplementary Figures and Legends:** (embedded below)

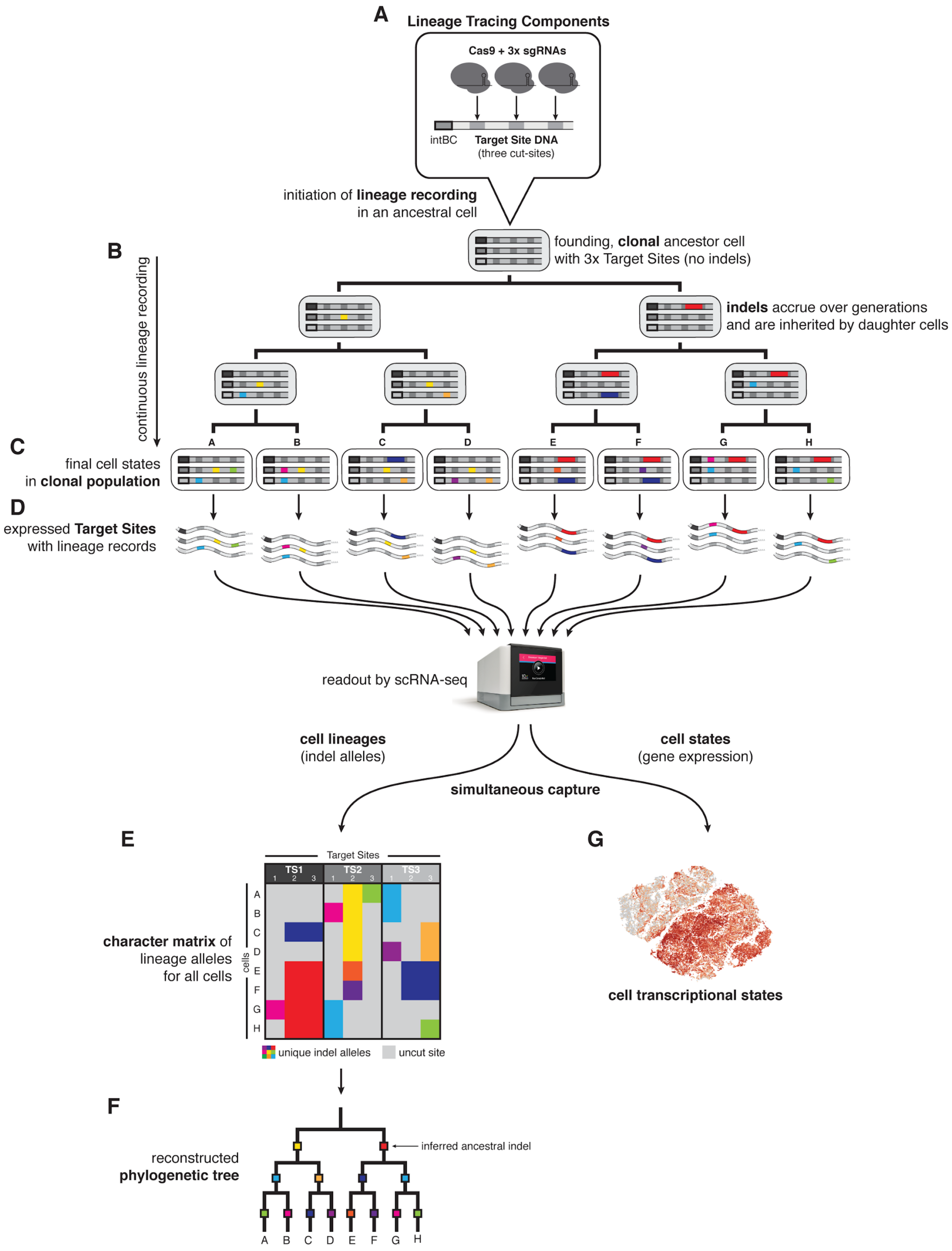
A
Lineage Tracing Components
Cas9 + 3x sgRNAs
intBC
Target Site DNA
(three cut-sites)
initiation of lineage recording in an ancestral cell
B
continuous lineage recording
founding, clonal ancestor cell with 3x Target Sites (no indels)
indels accrue over generations and are inherited by daughter cells
C
final cell states in clonal population
A B C D E F G H
D
expressed Target Sites with lineage records
readout by scRNA-seq
cell lineages (indel alleles)
cell states (gene expression)
simultaneous capture
E
Target Sites
TS1 TS2 TS3
1 2 3
cells
A B C D E F G H
character matrix of lineage alleles for all cells
unique indel alleles
uncut site
G
cell transcriptional states
F
reconstructed phylogenetic tree
inferred ancestral indel
A B C D E F G H

**Fig. S1. Detailed schematic of lineage tracing methodology.** (**A**) Cells are genetically engineered with the lineage tracing components: (i) Cas9, (ii) multiple copies of the Target Site, and finally (iii) three sgRNAs that are complementary to three cut-sites on each Target Site. Multiple copies of the Target Site per cell are distinguished by unique integration barcodes (intBCs). Cas9-induced double-stranded breaks at cut-sites are repaired with high-diversity insertions or deletions (indels), which act as stable, heritable markers of cell lineage. (**B**) Upon initiation of lineage recording in a founding cell (i.e., one "clone"), indels continuously accrue on the Target Sites (colored boxes), which are inherited by descendents over subsequent generations. (**C**) At the end of the lineage recording experiment, the final population of descendent cells (i.e., one "clonal population") are collected and (**D**) their expressed Target Site mRNAs are captured by single-cell RNA-sequencing (e.g., by the 10X Genomics Chromium platform) alongside transcriptome-wide expressed genes. (**E**) The indel allele information is read from the Target Site sequences and summarized in a "character matrix" of indel allele states (values) for each cut-site (columns) in each cell (rows). (**F**) From the pattern of cells' shared and distinguishing indel alleles, a tree reconstruction algorithm builds a phylogenetic model that best captures cell–cell relationships (e.g., by maximizing parsimony), thus producing a detailed map of cell lineage. (**G**) Simultaneously, single-cell RNA-sequencing captures the transcriptional profiles of the observed cells, allowing for direct comparisons between cell lineage and cell state.

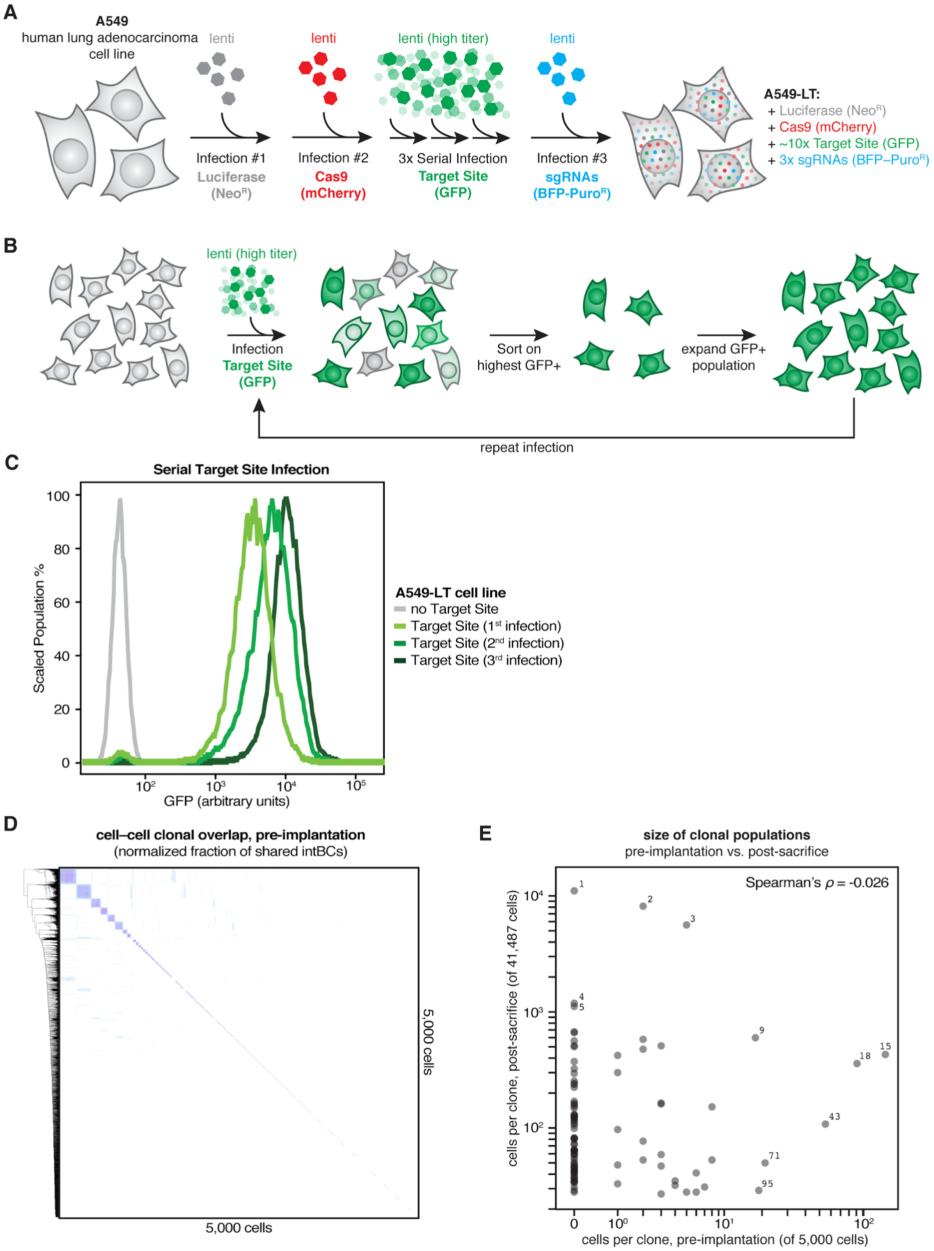


**Fig. S2. Cell line engineering strategy and estimation of clonal diversity.** (**A**) Human lung adenocarcinoma (A549) cells were genetically engineered with the lineage tracing components by lentiviral transduction with (1) Luciferase-Neomycin$^R$ and antibiotic-selected, (2) Cas9-mCherry and fluorescence-sorted, (3) serial, high-titer TargetSite-GFP and fluorescence-sorted, and finally (4) triple-sgRNA BFP-Puromycin$^R$ and fluorescence-sorted, thus producing lineage tracing-competent A549-LT cells. (**B**) Serial, high-titer TargetSite-GFP lentiviral transduction strategy to achieve high copy-number. (**C**) Cells with high-shifted GFP fluorescence after successive TargetSite-GFP infections, indicating increasing copy-number of the Target Site. (**D**) Sample of 5,000 A549-LT cells prior to injection to estimate initial clonal diversity. Shown is a heat-map of the fraction of shared intBCs in all cell–cell comparisons. On average, approximately 22,000 unique, high-quality intBCs were identified per random sample of 5,000 cells; thus, we estimate approximately 2,150 distinct clones per 5,000 cells (assuming 10.3 intBCs per clonal population; **Fig. S6A**) at the beginning of the experiment. (**E**) Comparison of the size of each clonal population observed in mouse M5k in a pre-implantation sample of 5,000 cells (*in vitro*) and post-sacrifice (*in vivo*). There is no correlation between the *in vitro* and *in vivo* population sizes (Spearman's $\rho$=-0.026).

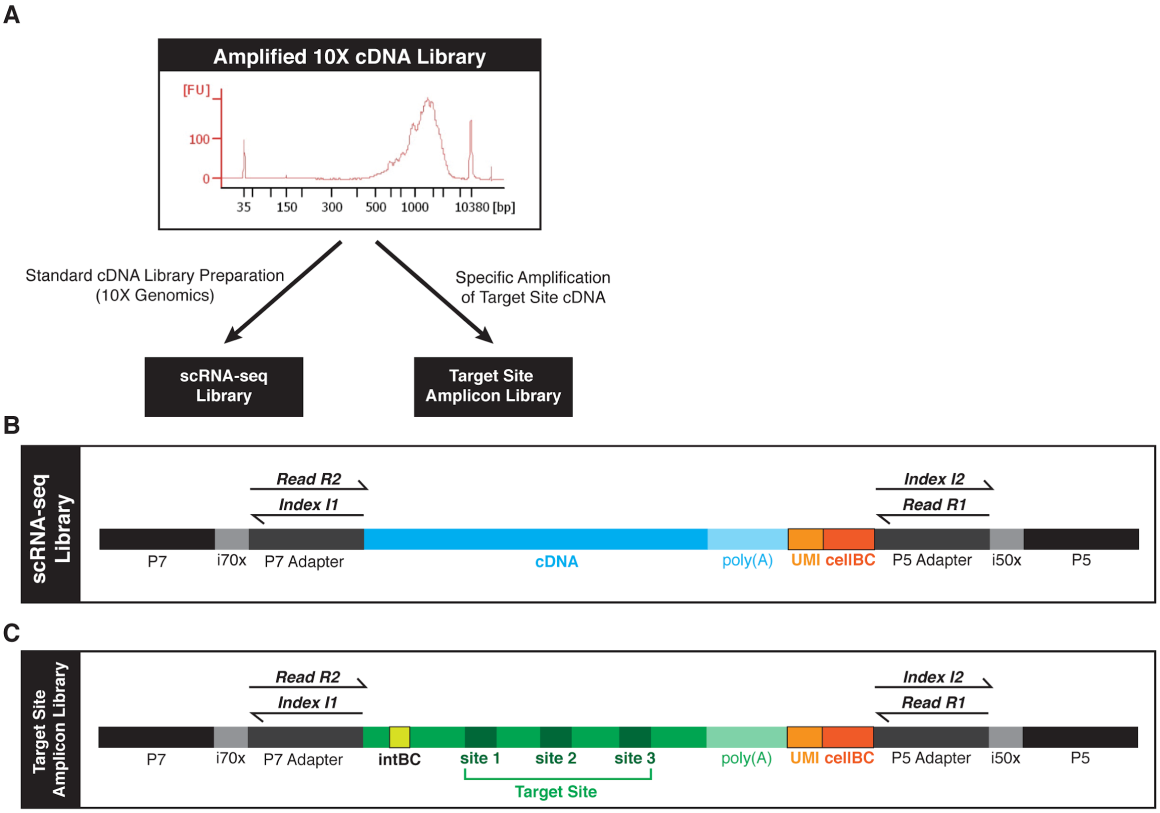


D

| Read | Cycles | Feature(s) |
|---|---|---|
| Read 1 | 26x | cellBC, UMI |
| Index 1 | 8x | sample index (1) |
| Index 2 | 8x | sample index (2) |
| Read 2 | >250x | expressed cDNA; OR intBC, cellBC |

E

| | M5k | M10k | M100k | M30k |
|---|---|---|---|---|
| **Raw reads (RNA)** | 2,630,541,106 | 1,769,228,115 | 1,507,857,428 | 1,100,492,472 |
| **Raw cells (RNA)** | 43,423 | 3,364 | 7,290 | 22,360 |
| **Mean Pre-normalization reads / cell (RNA)** | 60,579 | 525,929 | 206,839 | 49,217 |
| **Mean Post-normalization reads / cell (RNA)** | 52,243 | 303,364 | 170,014 | 40,973 |
| **Median UMIs / cell (RNA)** | 6,076 | 39,010 | 37,559 | 11,533 |
| **Median genes / cell (RNA)** | 1,862 | 5,893 | 5,234 | 2,861 |
| **Processed reads (TargetSite)** | 342,657,918 | 92,686,488 | 39,198,115 | 155,704,017 |
| **Raw cells (TargetSite)** | 45,420 | 3,120 | 7,453 | 22,053 |
| **Mean reads / UMI (TargetSite)** | 80.5 | 287.4 | 28.8 | 77.3 |
| **Mean UMIs / cell (TargetSite)** | 93.8 | 103.3 | 182.7 | 91.4 |

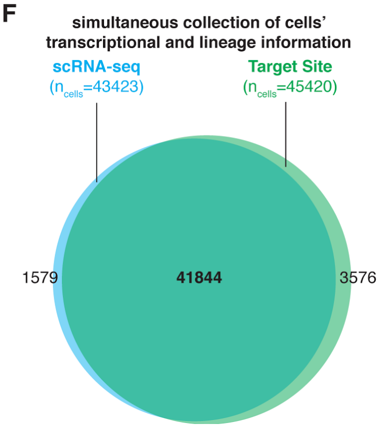


**Fig. S3. Sequencing library construction and metrics.** (**A**) The amplified cDNA (shown here as a BioAnalyzer trace) from the Chromium 3' Single Cell V2 kit (10X Genomics) serves as the template for both (**B**) the single-cell gene expression ("RNA") library and (**C**) the single-cell Target Site amplicon library (Methods). (**D**) The cellBC and unique molecular identifier (UMI) are sequenced from Read R1, the sample identities are sequenced from Indices I1 and I2, and the expressed cDNA or the Target Site amplicon (including the intBC and cut-sites 1, 2, and 3) are sequenced from Read R2. (**E**) Library sequencing metrics for TargetSite and RNA libraries for each mouse. (**F**) There is vast overlap in the cells identified from the gene expression and lineage sequencing datasets, as shown for mouse M5k.

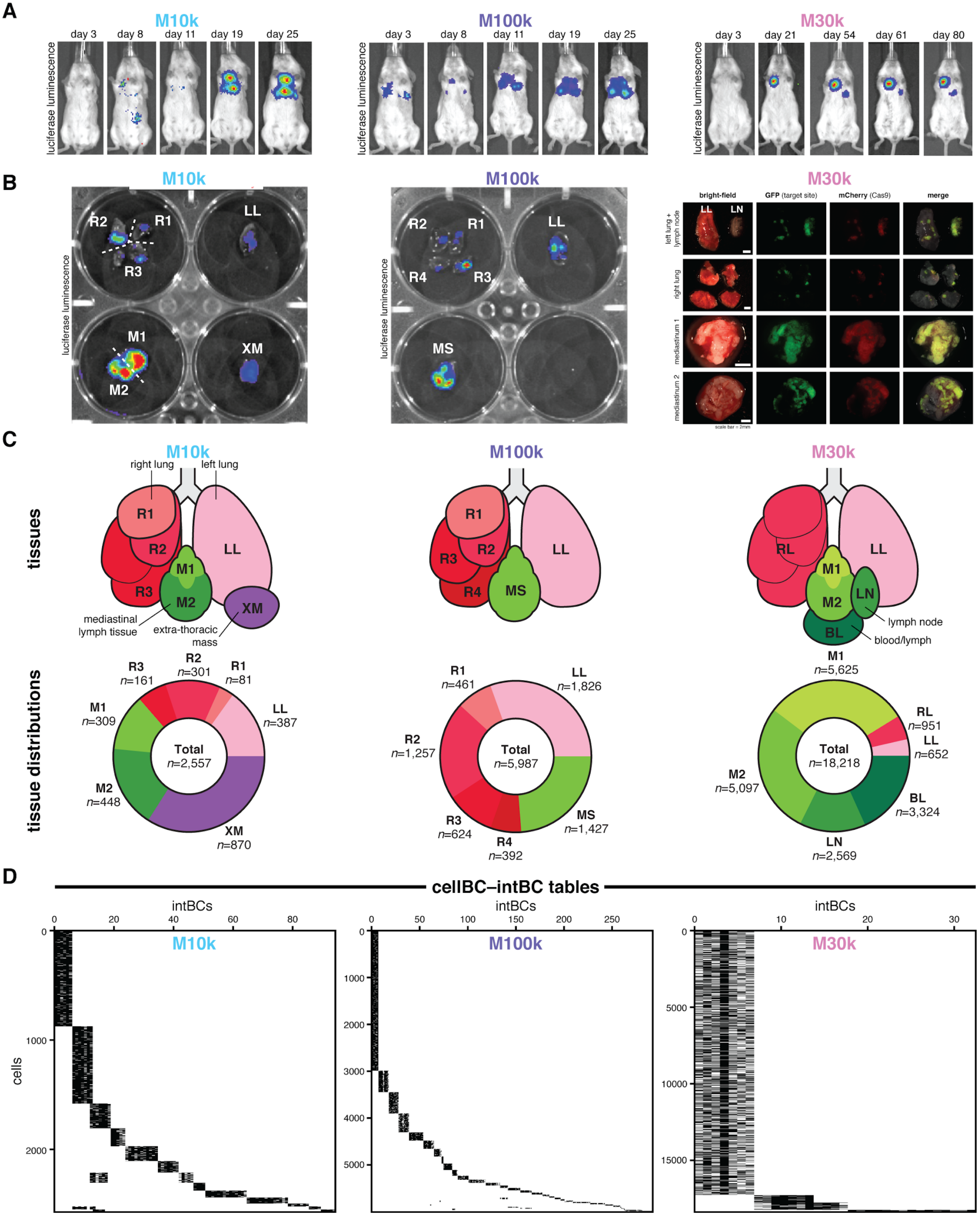
A
M10k
day 3
day 8
day 11
day 19
day 25
luciferase luminescence
M100k
day 3
day 8
day 11
day 19
day 25
M30k
day 3
day 21
day 54
day 61
day 80
B
M10k
R2
R1
R3
LL
M1
M2
XM
M100k
R2
R1
R4
R3
LL
MS
M30k
bright-field
GFP (target site)
mCherry (Cas9)
merge
left lung + lymph node
right lung
mediastinum 1
mediastinum 2
LL
LN
C
tissues
tissue distributions
M10k
right lung
left lung
R1
R2
R3
LL
M1
M2
XM
mediastinal lymph tissue
extra-thoracic mass
R3 n=161
R2 n=301
R1 n=81
LL n=387
M1 n=309
M2 n=448
XM n=870
Total n=2,557
M100k
R1
R2
R3
R4
LL
MS
R1 n=461
LL n=1,826
R2 n=1,257
R3 n=624
R4 n=392
MS n=1,427
Total n=5,987
M30k
RL
LL
M1
M2
LN
BL
lymph node
blood/lymph
M1 n=5,625
RL n=951
LL n=652
M2 n=5,097
BL n=3,324
LN n=2,569
Total n=18,218
D
cellBC–intBC tables
intBCs
cells
M10k
M100k
M30k

**Fig. S4. Tracing the cell lineages of metastatic progression in three additional mice.** In addition to mouse M5k discussed in throughout the main text, we also traced the lineages of metastatic dissemination in three additional mice orthotopically xenografted with 10,000 (M10k, left), 100,000 (M100k, middle), and 30,000 (M30k, right) A549-LT cells in two cohort experiments (first cohort, A549-LT1: M10k and M100k; second cohort, A549-LT2: M5k and M30k). (**A**) *In vivo* bioluminescence imaging of cancer cell engraftment and metastatic spread at indicated times post-implantation. The mice were sacrificed 53, 67, and 80 days post-implantation, respectively. (**B**) *Ex vivo* imaging of tumorous tissues by bioluminescence (left and middle) or fluorescence (right) imaging with the tissue samples indicated. In addition to extensive tumors in the lungs and mediastinum, M10k had one large solid tumor located ventral to the left lung and embedded in the ribcage (called here an "extra-thoracic mass"; XM). (**C**) Anatomical representation of collected tumorous tissues (top) and the number of cells collected for each tissue and each mouse (bottom). Notably, M30k had a large lymph node (LN) on the left lung, as well as diffuse bloody lymph (BL) in the thoracic cavity upon sacrificing. (**D**) CellBC-intBC tables showing clonal populations of cancer cells in each mouse, as in **Fig. S5A,B**. Some intBCs are shared between some clones (most notably in M10k) which likely resulted from cells that were clonally related at the stage of serial Target Site transduction during cell line engineering (Methods).

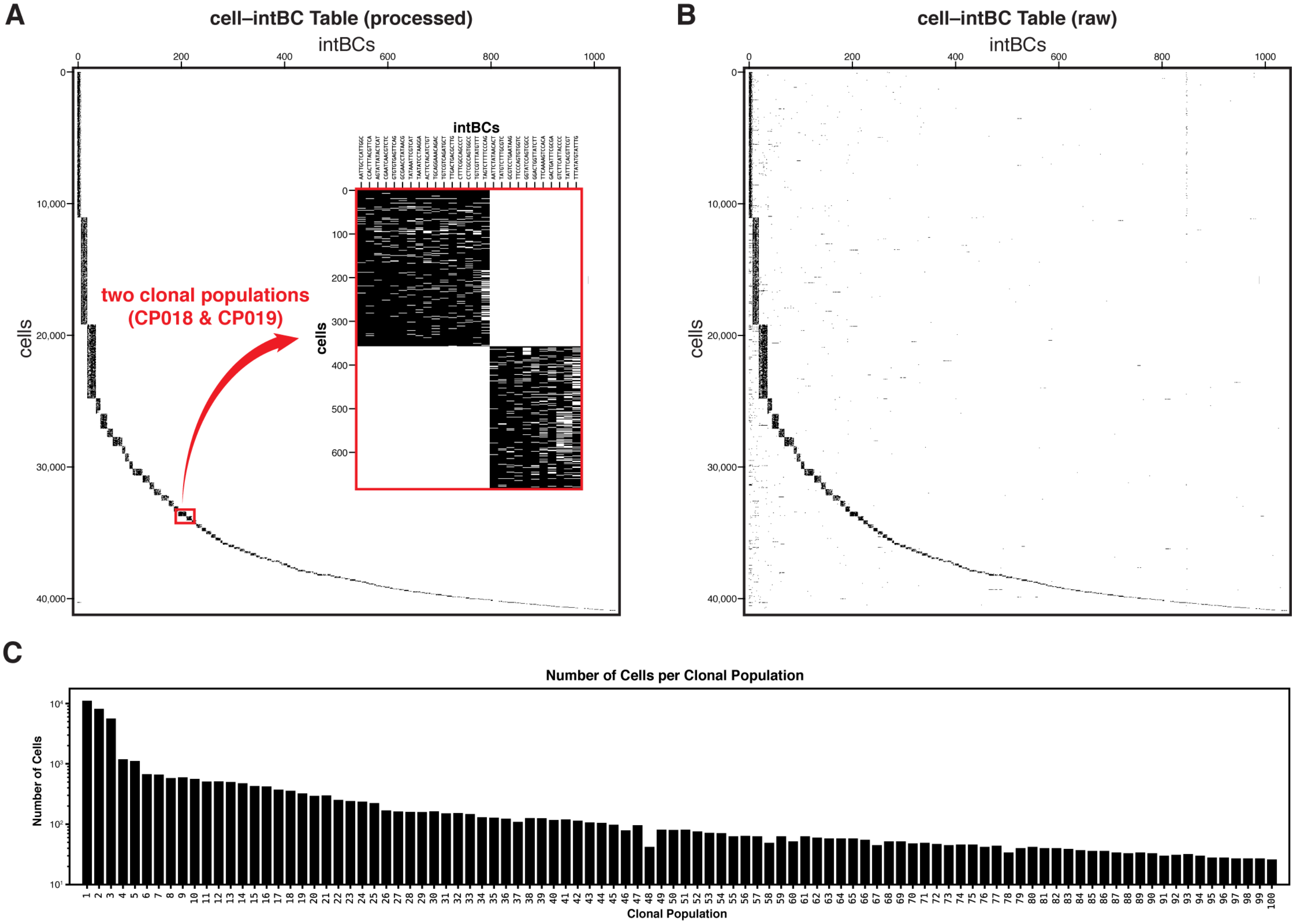


**Fig. S5. Identifying clonal populations by shared integration barcodes (intBCs).** (**A** and **B**) Tables representing the >1,000 unique integration barcodes (intBCs; columns) and >40,000 cells (rows) observed in mouse M5k, after processing (**A**) and before processing (**B**). Because intBCs are clonally inherited, cells that share identical intBCs are grouped into clonal populations (here, black blocks). (**A; inset**) The set of intBCs is generally exclusive to a single clonal population, such as those observed in clonal populations #18 and #19 (CP018 and CP109). (**B**) In the raw, unprocessed cell–intBC table, cells may be associated with intBCs that are not in their defined clonal set (indicated here by density outside of the black blocks). Through the processing pipeline, these conflicting intBCs (and/or the cells to which they pertain) are identified as cell doublets, cell-free transcripts in emulsion droplets, or sequencing artifacts and removed (Methods). (**C**) The number of cells per clonal population, generally numbered by descending population size.

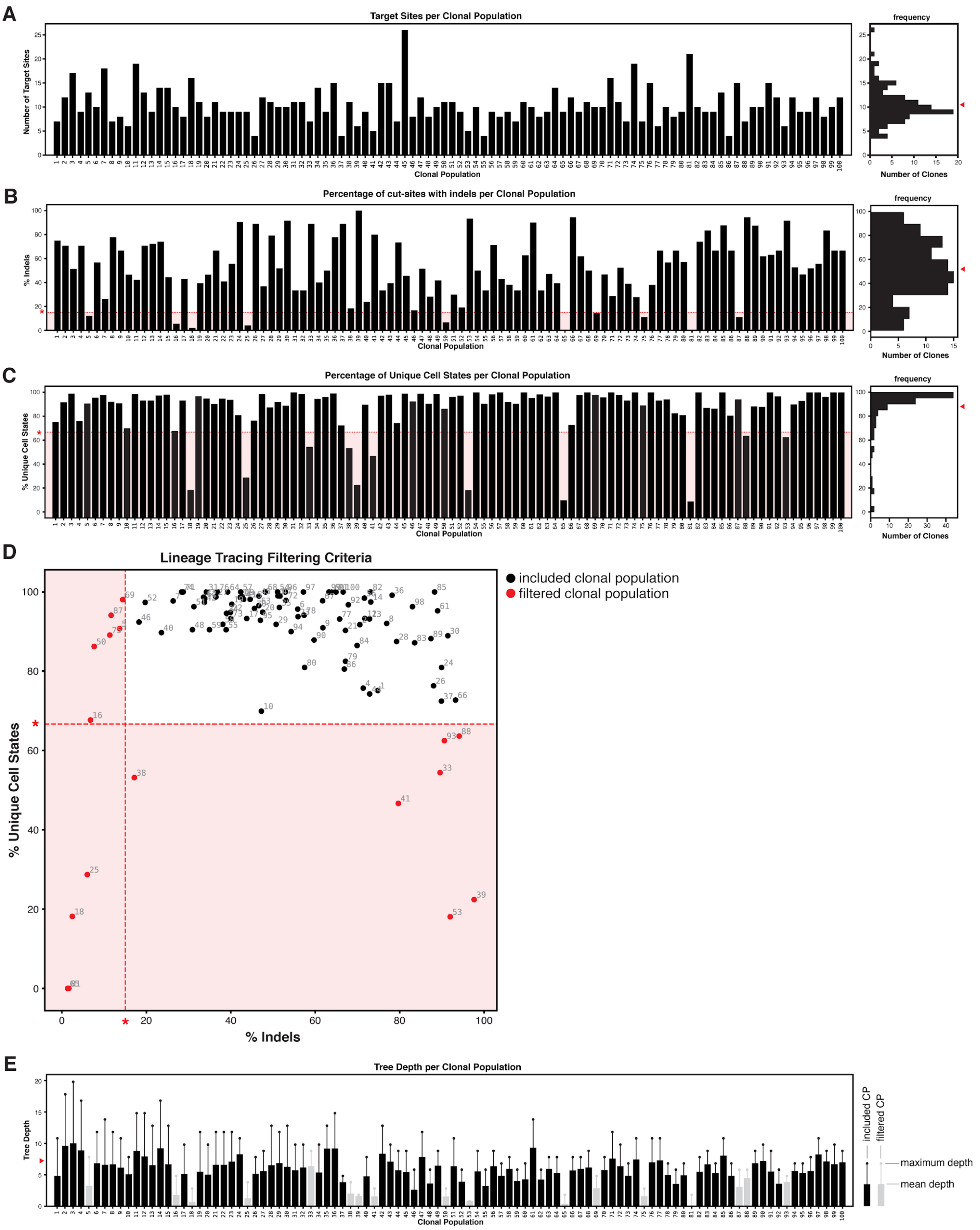
A
Target Sites per Clonal Population
Number of Target Sites
Clonal Population
frequency
Number of Clones
B
Percentage of cut-sites with indels per Clonal Population
% Indels
Clonal Population
frequency
Number of Clones
C
Percentage of Unique Cell States per Clonal Population
% Unique Cell States
Clonal Population
frequency
Number of Clones
D
Lineage Tracing Filtering Criteria
included clonal population
filtered clonal population
% Unique Cell States
% Indels
E
Tree Depth per Clonal Population
Tree Depth
Clonal Population
included CP
filtered CP
maximum depth
mean depth

**Fig. S6. Characteristics of the lineage tracer and quality-control of clonal populations.** (**A**) The copy-number of integrated Target Sites per clonal population, as determined by the number of unique intBCs. (**A, right**) The distribution of Target Site copy-number per clonal population; mean copy-number of Target Sites indicated (red arrowhead). (**B**) The percentage of cut-sites bearing lineage indel alleles per clonal population. Clonal populations with <15% indels are excluded (red asterisk, red underlay). (**B, right**) The distribution of indel-bearing cut-sites per clonal population; mean percentage indicated (red arrowhead). (**C**) The percentage of unique cell lineage states per clonal population (i.e., lineage diversity). Clonal populations with <66.7% diversity were excluded (red asterisk, red underlay). (**C, right**) The distribution of lineage diversity per clonal population; mean percentage indicated (red arrowhead). (**D**) Comparison of lineage tracing characteristics (% indels and % unique cell states) to define quality-control filtering criteria. Clonal populations removed by the filter (red closed circles) and filtering thresholds (red asterisks, red underlay) are indicated. (**E**) The depth of the reconstructed phylogenetic trees for each clonal population. Mean tree depths are shown as closed bars; maximum tree depths are shown as whiskers. Clonal populations that were excluded due to suboptimal lineage tracing characteristics (gray) and the average tree depth across all clonal populations (red arrowhead) are indicated.

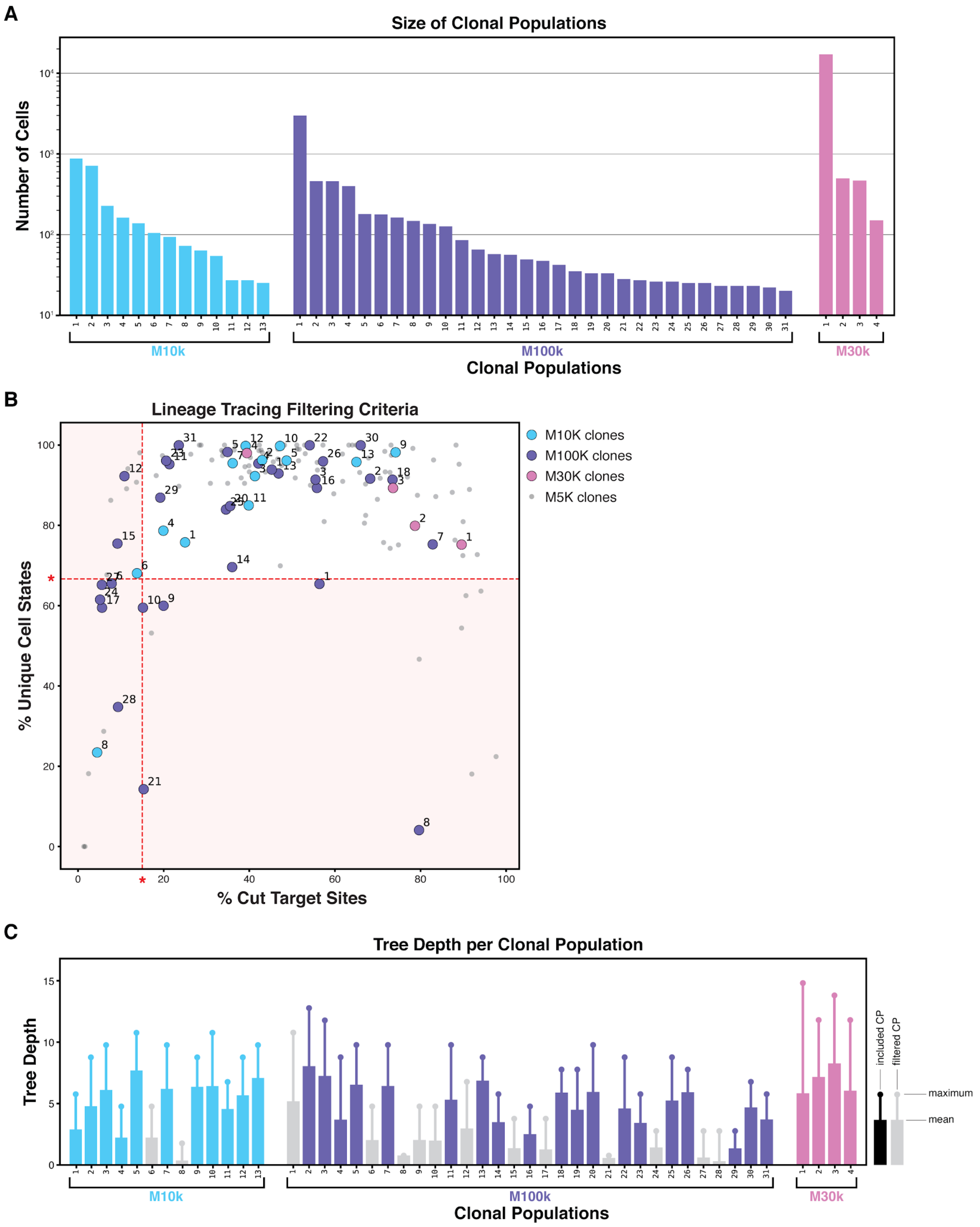
A
Size of Clonal Populations
Number of Cells
M10k
M100k
M30k
Clonal Populations
B
Lineage Tracing Filtering Criteria
% Unique Cell States
% Cut Target Sites
M10K clones
M100K clones
M30K clones
M5K clones
C
Tree Depth per Clonal Population
Tree Depth
included CP
filtered CP
maximum
mean
M10k
M100k
M30k
Clonal Populations

**Fig. S7. Lineage tracing characteristics of clonal populations in additional mice.** (**A**) Number of cells in each clonal population in M10k, M100k, and M30k mice. (**B**) Scatter plot of the percentage of cut-sites bearing indels and the percentage of unique cell states per clonal population, which are characteristics of the lineage tracer that influence tree reconstructability. Some clonal populations exhibited suboptimal parameters (red asterisks and red field) and were excluded from reconstruction and downstream analyses. (**C**) The mean and maximum depths of the reconstructed phylogenetic trees for each clonal population in additional mice, as in **Fig. S6E**.

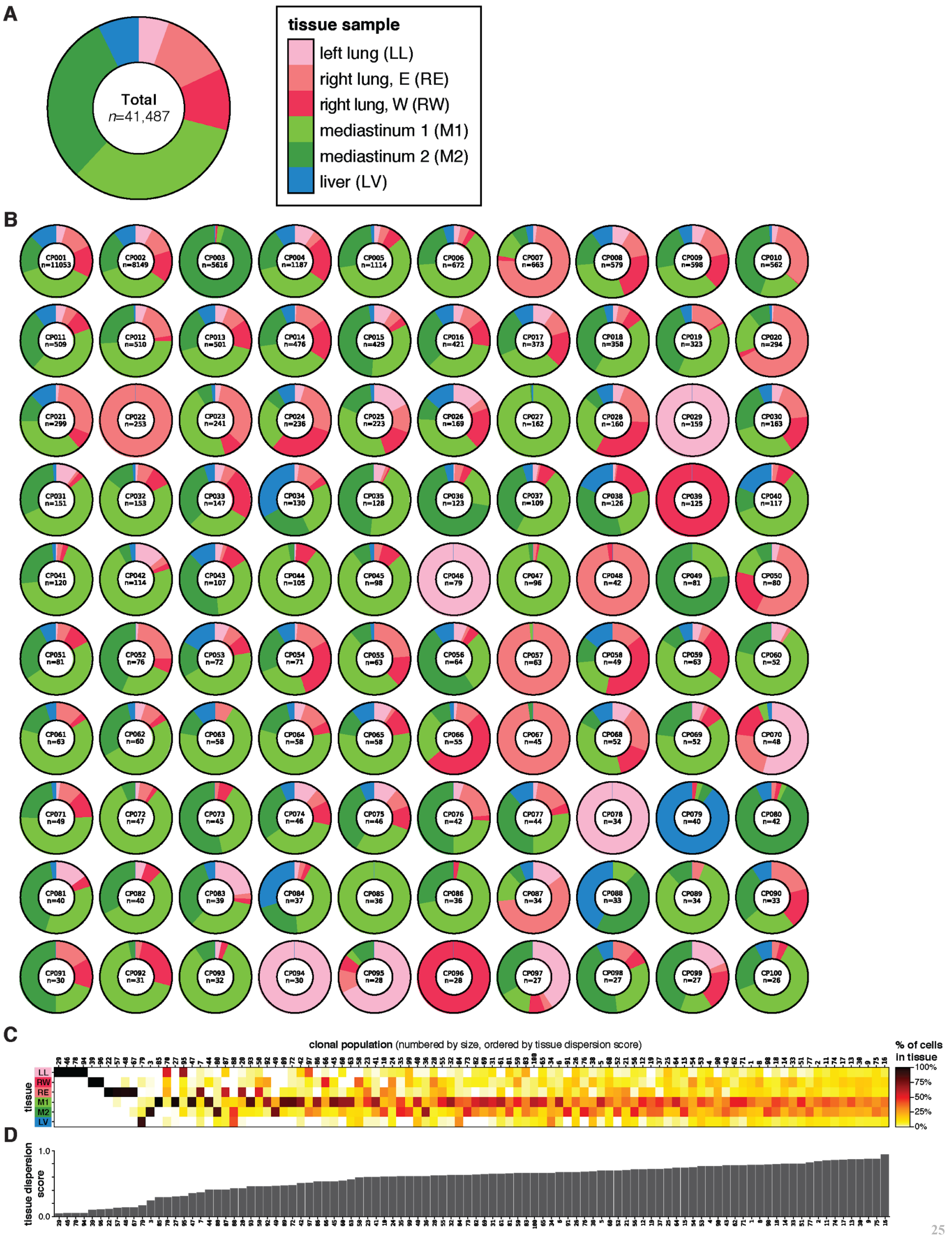

A
tissue sample
left lung (LL)
right lung, E (RE)
right lung, W (RW)
mediastinum 1 (M1)
mediastinum 2 (M2)
liver (LV)
Total
n=41,487
B
CP001 n=11053
CP002 n=8149
CP003 n=5616
CP004 n=1187
CP005 n=1114
CP006 n=672
CP007 n=663
CP008 n=579
CP009 n=598
CP010 n=562
CP011 n=509
CP012 n=510
CP013 n=501
CP014 n=476
CP015 n=429
CP016 n=421
CP017 n=373
CP018 n=358
CP019 n=323
CP020 n=294
CP021 n=299
CP022 n=253
CP023 n=241
CP024 n=236
CP025 n=223
CP026 n=169
CP027 n=162
CP028 n=160
CP029 n=159
CP030 n=163
CP031 n=151
CP032 n=153
CP033 n=147
CP034 n=130
CP035 n=128
CP036 n=123
CP037 n=109
CP038 n=126
CP039 n=125
CP040 n=117
CP041 n=120
CP042 n=114
CP043 n=107
CP044 n=105
CP045 n=98
CP046 n=79
CP047 n=96
CP048 n=42
CP049 n=81
CP050 n=80
CP051 n=81
CP052 n=76
CP053 n=72
CP054 n=71
CP055 n=63
CP056 n=64
CP057 n=63
CP058 n=49
CP059 n=63
CP060 n=52
CP061 n=63
CP062 n=60
CP063 n=58
CP064 n=58
CP065 n=58
CP066 n=55
CP067 n=45
CP068 n=52
CP069 n=52
CP070 n=48
CP071 n=49
CP072 n=47
CP073 n=45
CP074 n=46
CP075 n=46
CP076 n=42
CP077 n=44
CP078 n=34
CP079 n=40
CP080 n=42
CP081 n=40
CP082 n=40
CP083 n=39
CP084 n=37
CP085 n=36
CP086 n=36
CP087 n=34
CP088 n=33
CP089 n=34
CP090 n=33
CP091 n=30
CP092 n=31
CP093 n=32
CP094 n=30
CP095 n=28
CP096 n=28
CP097 n=27
CP098 n=27
CP099 n=27
CP100 n=26
C
clonal population (numbered by size, ordered by tissue dispersion score)
tissue
LL
RW
RE
M1
M2
LV
% of cells in tissue
100%
75%
50%
25%
0%
D
tissue dispersion score
1.0
0.0

**Fig. S8. Clonal populations exhibit distinct tissue distributions.** (**A**) The bulk distribution of all collected cells across the six tissue samples. (**B**) The distributions of cells from each clonal population across the six tissue samples. Some clonal populations were exclusive to the primary tissue (e.g., clonal population CP046), whereas some clones exhibited biased tissue distributions (e.g., CP003) and many others were observed broadly distributed across all tissues (e.g., CP011). (**C**) Tissue distributions of the largest 100 clonal populations. (**D**) The Tissue Dispersion Score is a statistical measurement of the distribution across tissues for each clonal population; *x*-axis shared with (**C**).

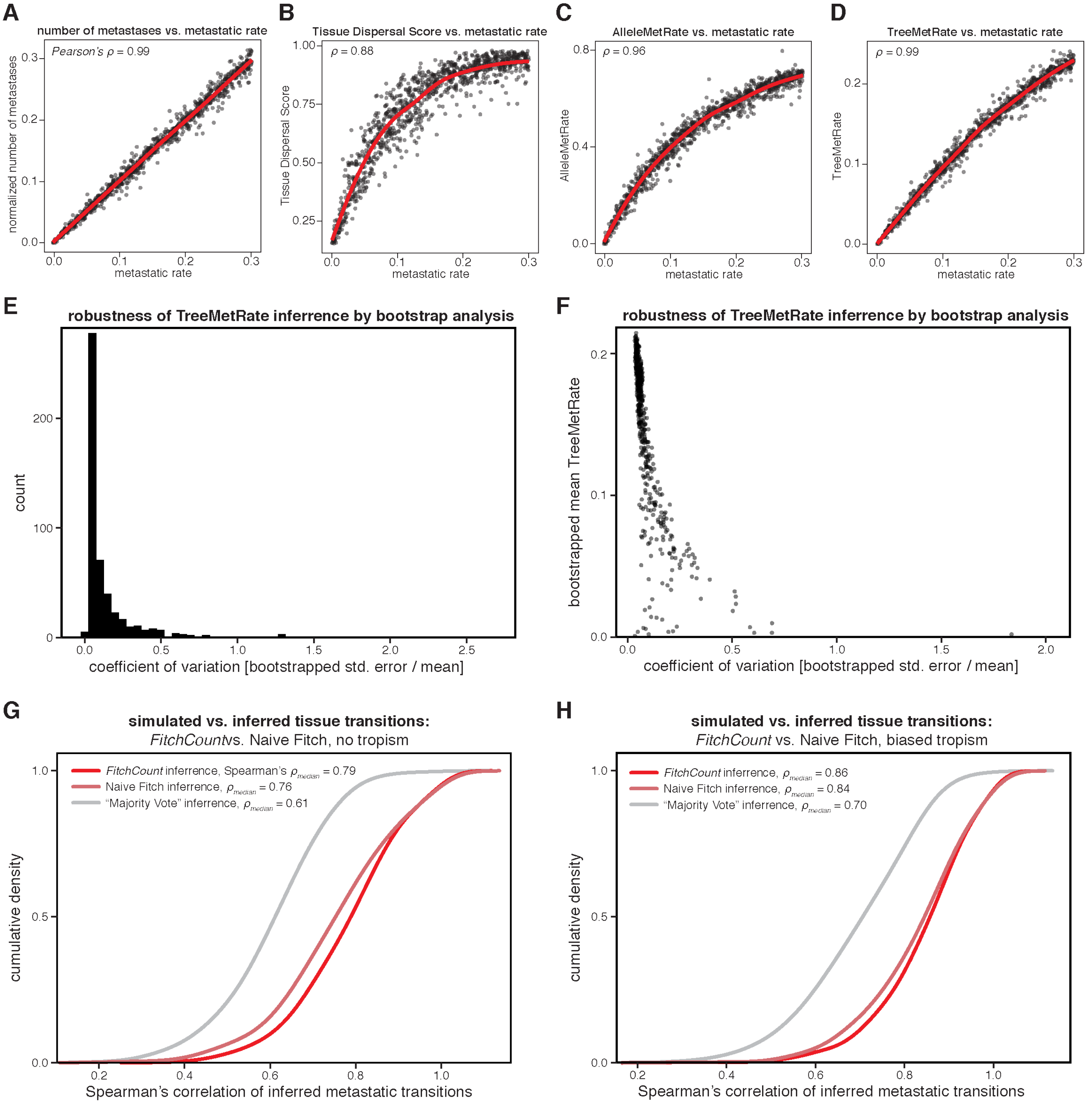


**Fig. S9. Assessing the accuracy of different measurements of metastatic rate and inference of tissue transitions using simulated lineages.** (**A–D**) Comparison between the simulated metastatic rate and various lineage tracer-derived statistics, with the correlation (Pearson's $\rho$) indicated; red lines represent moving average. (**A**) The normalized count of simulated metastatic transitions is very well correlated with the simulated metastatic rates, and serves as a ground-truth benchmark. (**B**) The Tissue Dispersal Score, which is a statistical measure of how closely a clone's tissue distribution matches the background tissue distributions, is correlated with the metastatic rate, but saturates at intermediate metastatic regimes. (**C**) The AlleleMetRate, or the proportion of cells whose closest relative by allele similarity is in a different tissue, is better correlated with metastatic rate. (**D**) The TreeMetRate, or the proportion of inferred metastases in a reconstructed phylogeny, is the best lineage-derived measurement of metastatic phenotype by correlation. (**E** and **F**) Robustness of the TreeMetRate inference by bootstrap analysis. (**E**) The distribution of the coefficients of variation of the TreeMetRate across 50,000 simulated, bootstrapped phylogenies (Methods). The small coefficient of variation indicates that the TreeMetRate is a robust measurement. (**F**) The TreeMetRate coefficient of variation is only relatively large when the mean TreeMetRate is small. (**G** and **H**) Cumulative density plots assessing the accuracy of the *FitchCount* strategy for inferring ancestral tissue transition from simulated phylogenies with or without simulated biased tissue transitions (thus approximating tropism; Methods: "*Assessing accuracy of the tissue transition matrices*"). *FitchCount* outperforms other inference approaches, as measured by the Spearman correlation between the inferred and the ground-truth conditional tissue transition probability matrices. *FitchCount* was benchmarked against two other inference methods: (i) a "naive" single-solution implementation of the Fitch-Hartigan maximum parsimony algorithm or (ii) "Majority Vote", which infers ancestral tissue location as the tissue in which the majority of cells below each clade-level reside. Conditional probabilities for each algorithm were obtained by row-normalizing the count matrix with respect to the non-diagonal counts in each row.

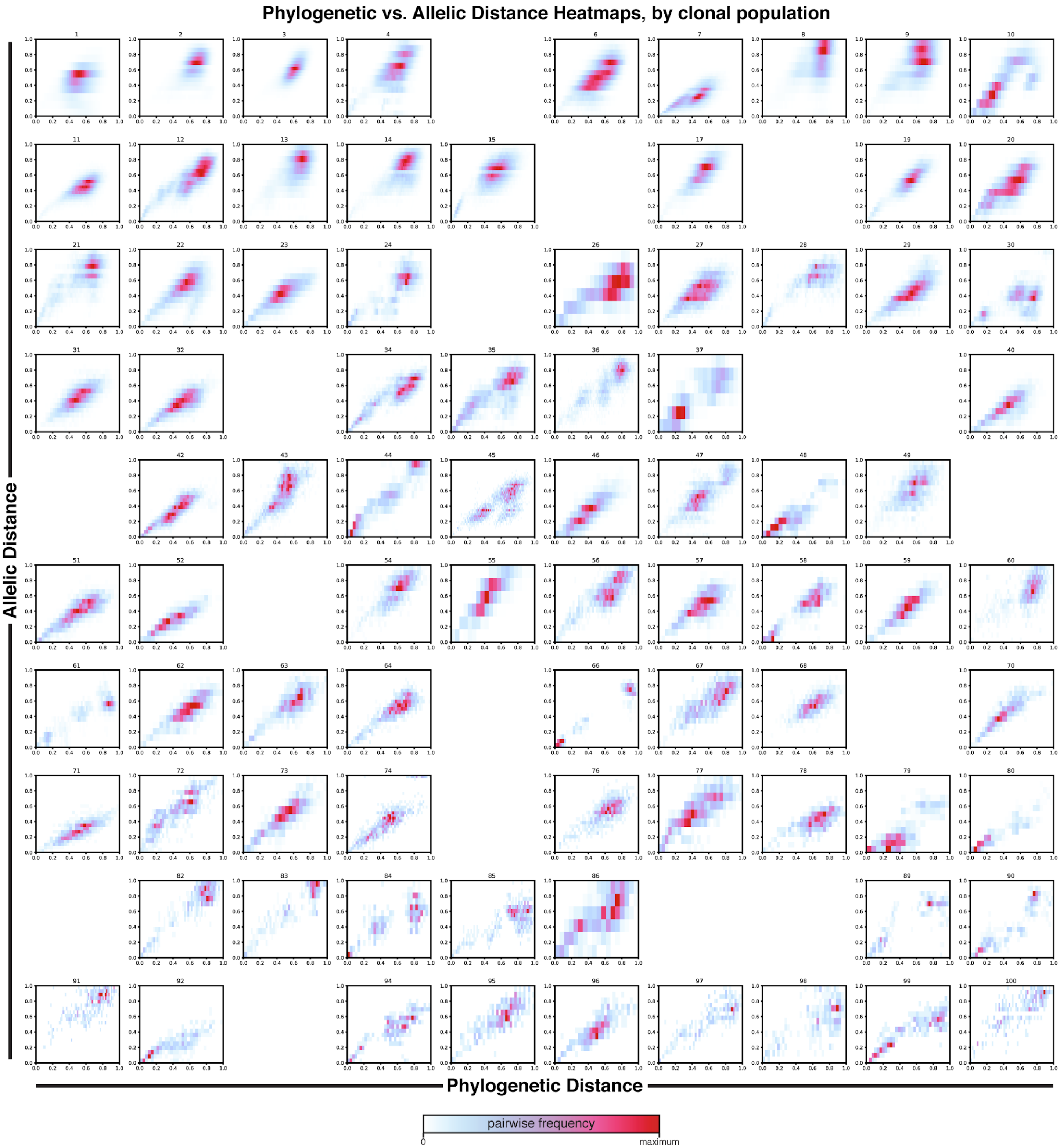


**Fig. S10. Relationship between phylogenetic distance and allelic distance for each clonal population.** Density heat-maps comparing phylogenetic distance (i.e., the normalized tree branch distance between two cells) and allelic distance (i.e., the normalized difference in lineage allele state between two cells) for all pairwise cell–cell relationships and for each clonal population, as in **Fig. 2C**. As expected, phylogenetic and allelic distances are correlated, suggesting that the reconstructed trees are a good phylogenetic model of cell–cell relationships. Excluded clonal populations are not shown.

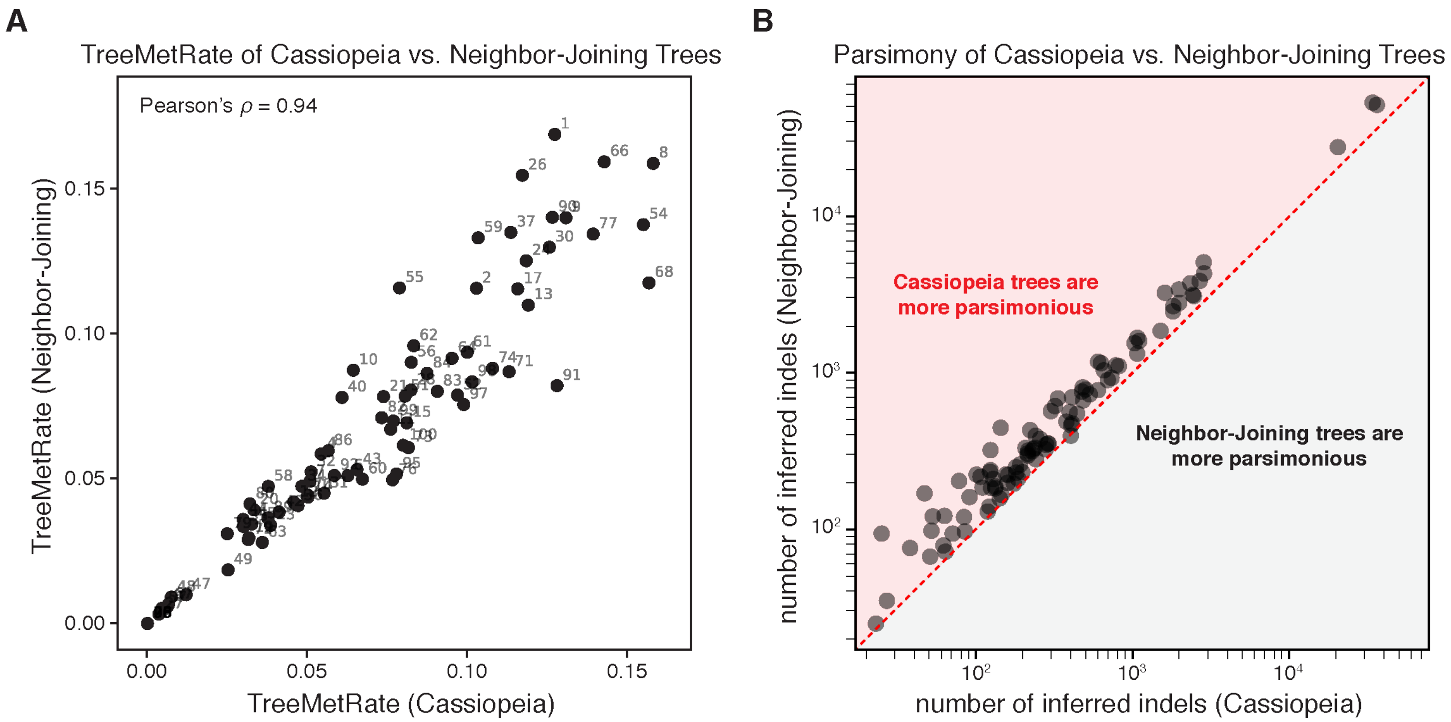


**Fig. S11. The TreeMetRate is stable across tree reconstruction algorithms (Cassiopeia versus Neighbor-Joining).** (**A**) Comparison of the TreeMetRates for each clonal population from Cassiopeia trees and Neighbor-Joining trees. The TreeMetRates are correlated for both the Cassiopeia and Neighbor-Joining trees (Pearson's $\rho$=0.94). (**B**) Comparison of the parsimony of Cassiopeia trees and Neighbor-Joining trees, defined as the number of inferred indels in each tree. Notably, the Cassiopeia trees are more parsimonious than the Neighbor-Joining trees (i.e., they have fewer inferred indels; red overlay).

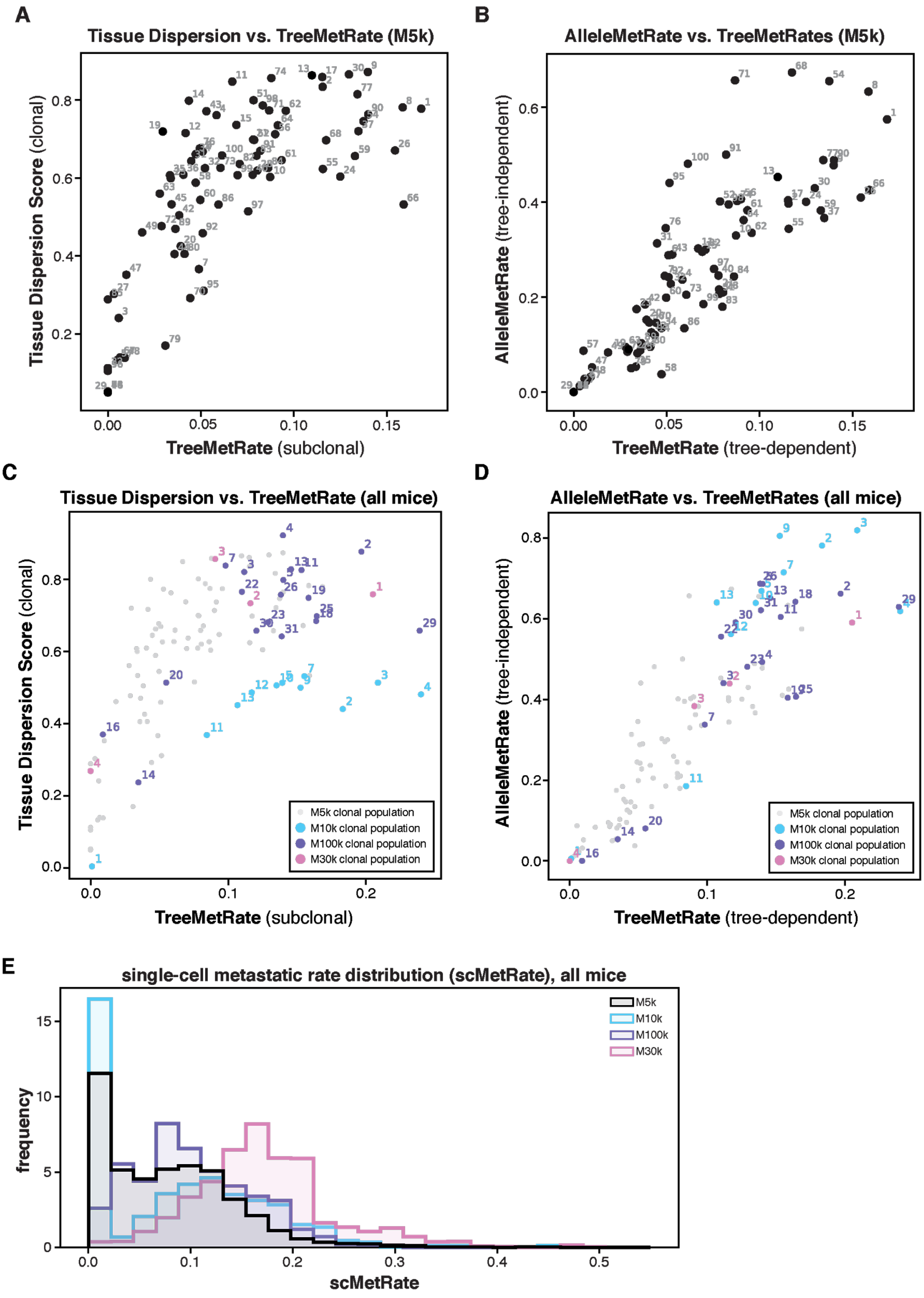

A
Tissue Dispersion vs. TreeMetRate (M5k)
Tissue Dispersion Score (clonal)
TreeMetRate (subclonal)
B
AlleleMetRate vs. TreeMetRates (M5k)
AlleleMetRate (tree-independent)
TreeMetRate (tree-dependent)
C
Tissue Dispersion vs. TreeMetRate (all mice)
Tissue Dispersion Score (clonal)
TreeMetRate (subclonal)
M5k clonal population
M10k clonal population
M100k clonal population
M30k clonal population
D
AlleleMetRate vs. TreeMetRates (all mice)
AlleleMetRate (tree-independent)
TreeMetRate (tree-dependent)
M5k clonal population
M10k clonal population
M100k clonal population
M30k clonal population
E
single-cell metastatic rate distribution (scMetRate), all mice
frequency
scMetRate
M5k
M10k
M100k
M30k

**Fig. S12. Clonal populations exhibit broad metastatic phenotypes, measured by Tissue Dispersal Score, AlleleMetRate, and TreeMetRate.** (**A–D**) We evaluated the metastatic phenotype of each clonal population from mouse M5k (**A, B**) and the additional mice M10k, M100k, and M30k (**C, D**) using our three lineage-derived measurements: Tissue Dispersal Score (as in **Fig. S8D**), AlleleMetRate, and TreeMetRate (as in **Fig. 3C**). These three measurements follow similar relative trends across all four mouse experiments. First, the clonal populations exhibit a broad range of metastatic phenotypes. Second, all three measurements are correlated with one another, though simulations indicate that the TreeMetRate is the most accurate for estimating the underlying metastatic rate (**Fig. S9**). Third, Tissue Dispersal Score saturates at intermediate metastatic regimes (**A** and **C**). (**E**) The distribution of single-cell-resolution metastatic rates (scMetRates) across all cells for each mouse (as in **Fig. 3D**). Though all mice have broad distributions of metastatic phenotypes, mouse M5k (black) is particularly well represented by cells in low-to-intermediate metastatic regimes whereas mouse M30k (pink) has very few cells in the low metastatic regime.

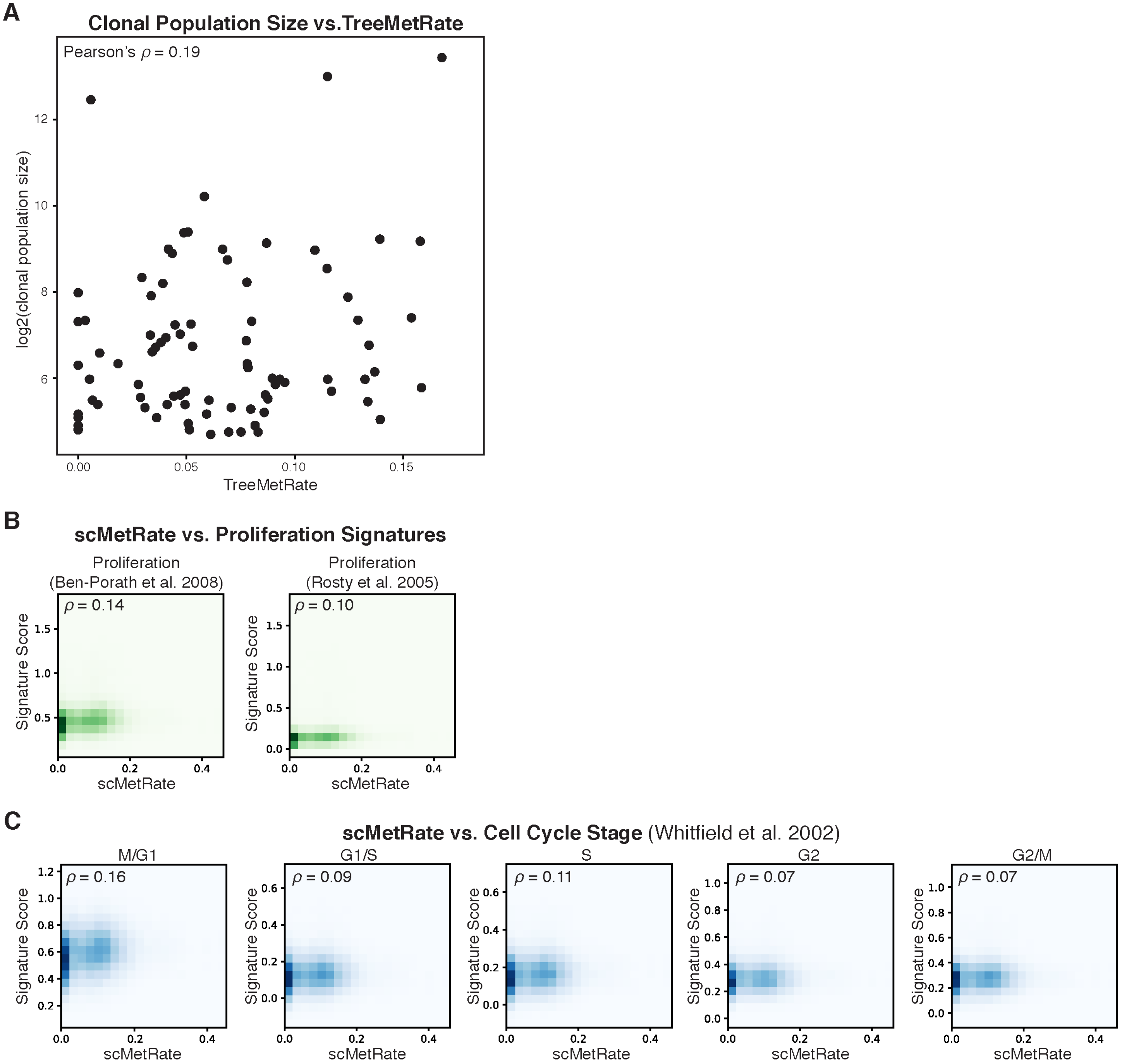


**Fig. S13. The scMetRate measures metastatic potential decoupled from proliferative capacity.** (**A**) There is poor correlation between the scMetRate and the log2-transformed clonal population size, a proxy for clonal fitness. (**B** and **C**) Density heat-maps comparing the scMetRate and various transcriptional signatures. Notably, the scMetRate is poorly correlated with transcriptional signatures of proliferation (**B**) nor stages of the cell cycle (**C**). Pearson's correlations ($\rho$) are indicated for each subplot.

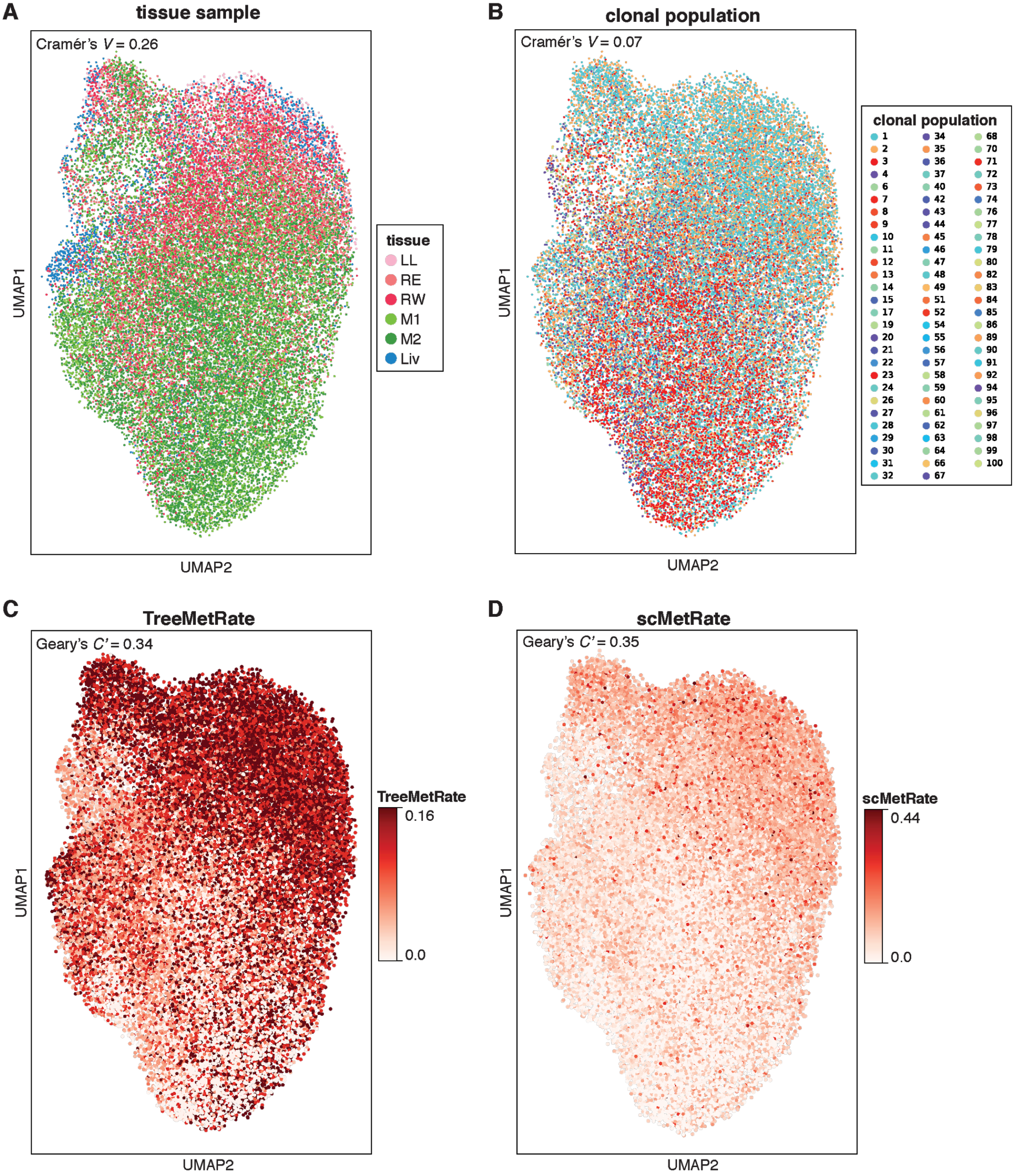


**Fig. S14. Effects of tissue sample, clonal population, and metastatic rate on transcriptional state.** To identify global trends in the gene expression data, we used *Vision* (*52*) to statistically assess the transcriptional effect of four features: (**A**) tissue sample, (**B**) clonal population identity, (**C**) TreeMetRate, and (**D**) scMetRate. The transcriptional states are represented here as a two-dimensional projection using Uniform Manifold

Approximation and Projection (UMAP; B). Distinctions in transcriptional state are not predominantly explained by the clonal population (**B**; by Cramér's *V*), though there is modest association between transcriptional state and both tissue sample and metastatic phenotype (by Cramér's *V* and inverted Geary's *C'*, respectively).

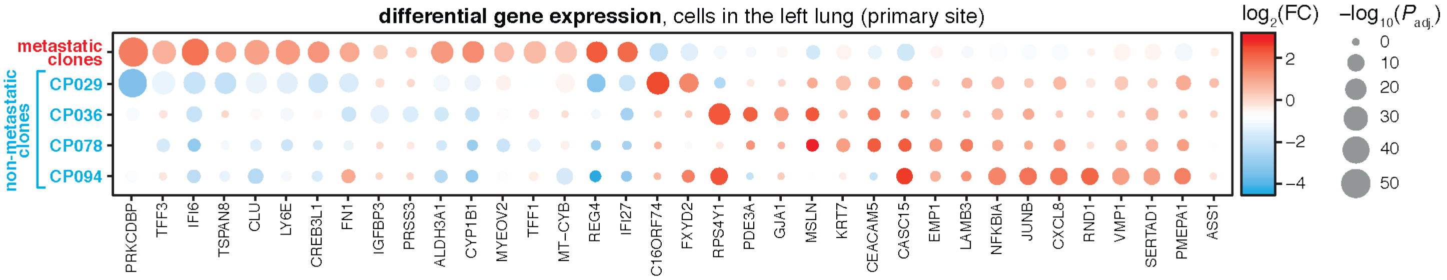


**Fig. S15. Differential expression between non-metastatic and metastatic clonal populations in the primary tissue.** Differential gene expression analysis comparing four non-metastatic clonal populations (CP029, 36, 78, and 94) and all metastatic clonal populations in the primary tumor tissue (i.e., all other cells in the left lung). Significantly differentially expressed genes are colored by the log2-transformed fold-change in gene expression and scaled by the adjusted Wilcoxon rank-sum test *P*-value.

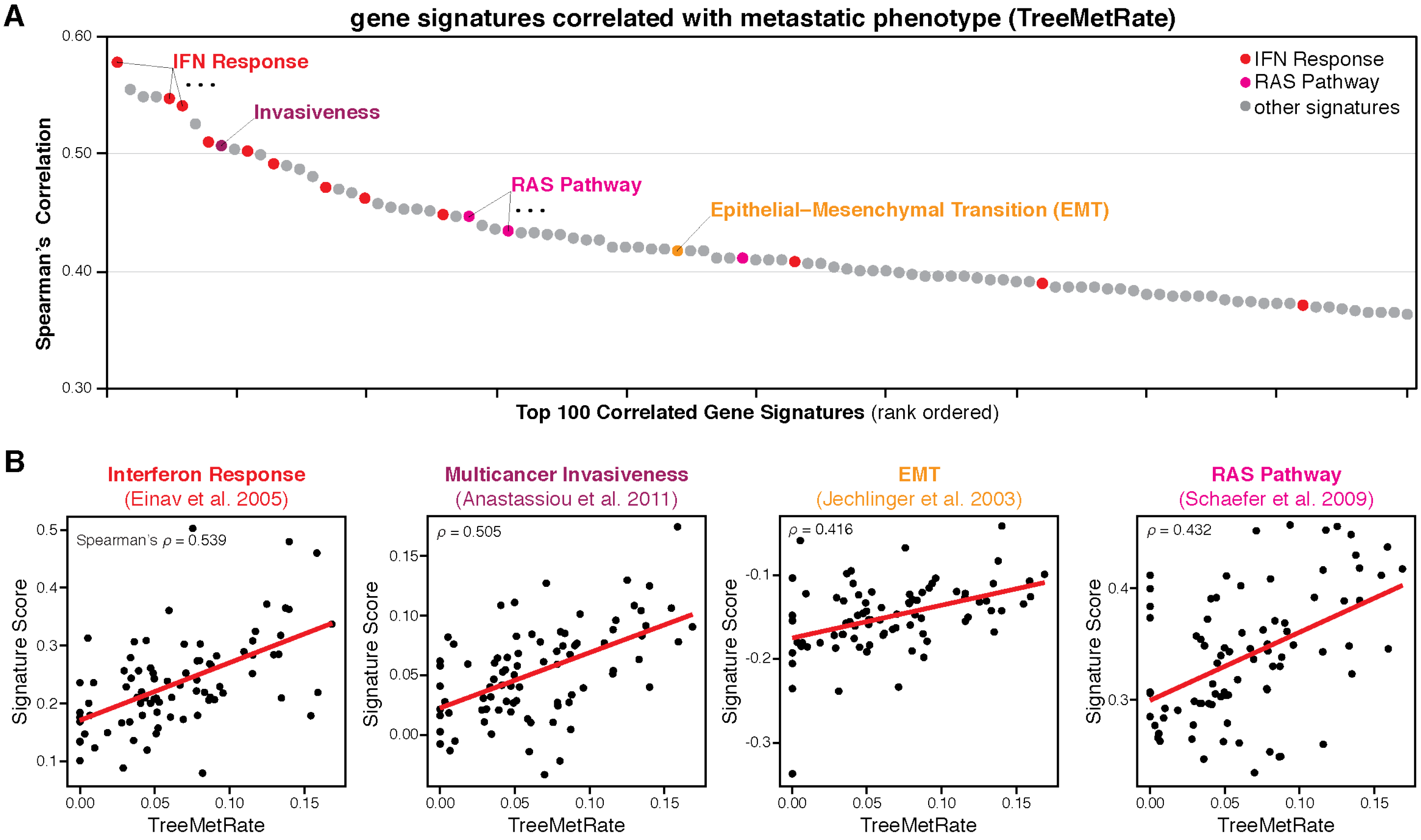


**Fig. S16. Metastasis-related gene signatures are correlated with metastatic potential.** (**A**) Rank-ordered gene signatures that are the most positively correlated with TreeMetRate, including many related to interferon response (red) and RAS pathways (magenta), as well as other metastasis-related signatures. (**B**) Scatter plots showing the correlation between TreeMetRate and noted gene signature scores per clonal population; Spearman's correlation ($\rho$) indicated.

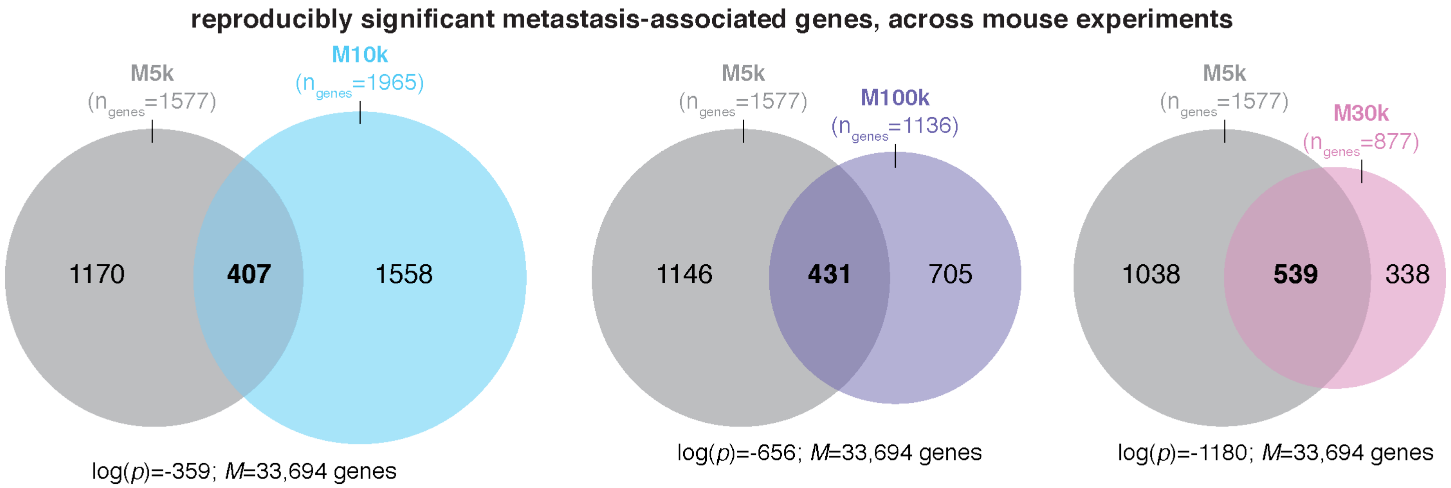


**Fig. S17. Many of the same genes are associated with metastatic phenotype across all mice.** Using the same regression strategy as in the analysis of mouse M5k, we found many genes with expression that is significantly associated with high or low scMetRates. The number of significant genes for each mouse (FDR < 0.01) and their overlap in the same direction with mouse M5k (gray) are shown ($n_{genes}$). In all cases, the overlap between mouse M5k and each additional mouse is significant by hypergeometric test ($p$-value and $M$ indicated).

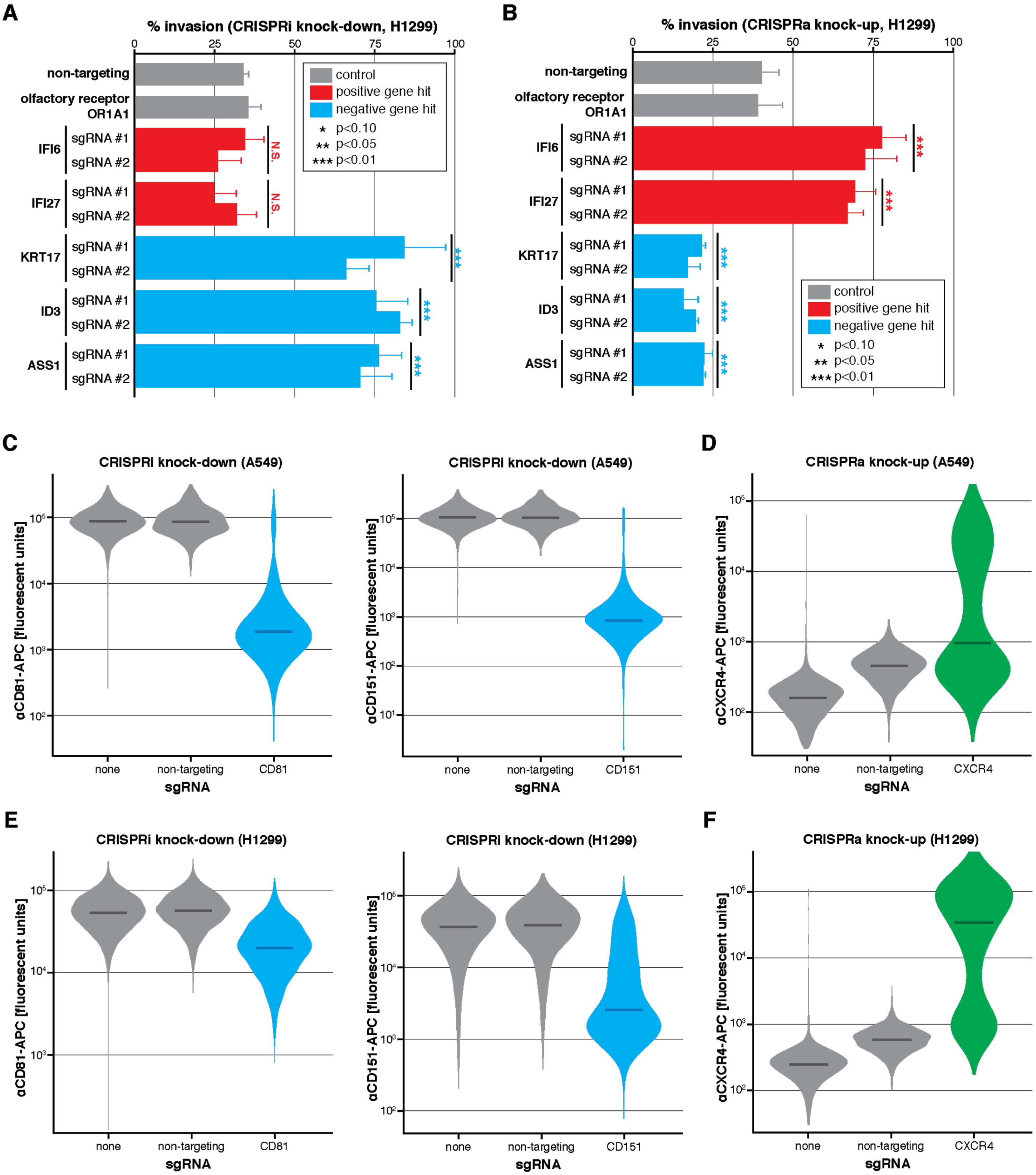


**Fig. S18. Functional validation of five gene candidates in a different cell line (H1299s) and validation of CRISPRi and CRISPRa activity.** (**A** and **B**) *In vitro* transwell invasion assays following CRISPRi or CRISPRa gene perturbation, respectively, in H1299 cells; as in **Fig.4E, F**. Perturbation of positive and negative metastasis-associated gene candidates were performed in triplicate using two independent sgRNAs per gene. Differences in invasion phenotype relative to two negative control guides (non-targeting and olfactory receptor) were significant by two-tailed *t*-test. N.S., not significant; error bars show standard deviation. (**C, E**) A549-CRISPRi cells or H1299-CRISPRi cells, respectively, were treated with no sgRNA, non-targeting sgRNA, or sgRNAs against either CD81 or CD151, two highly expressed cell-surface markers. One week following treatment, the cells were collected, stained with APC-labelled anti-CD81 or anti-CD151 antibodies and their fluorescence was measured by flow cytometry. Shown here is substantial knock-down of CD81 or CD151 gene expression relative to no sgRNA or non-targeting sgRNA controls. (**D, F**) The same validation experiment as conducted in **C** and **E**, but for A549-CRISPRa cells or H1299-CRISPRa cells, respectively, treated with an sgRNA against CXCR4, a lowly expressed cell-surface marker. Shown here is substantially increased CXCR4 gene expression relative to no sgRNA or non-targeting sgRNA controls. Violin plots show distribution of fluorescent signal for each cell identified by flow cytometry; median marked by dark bar.

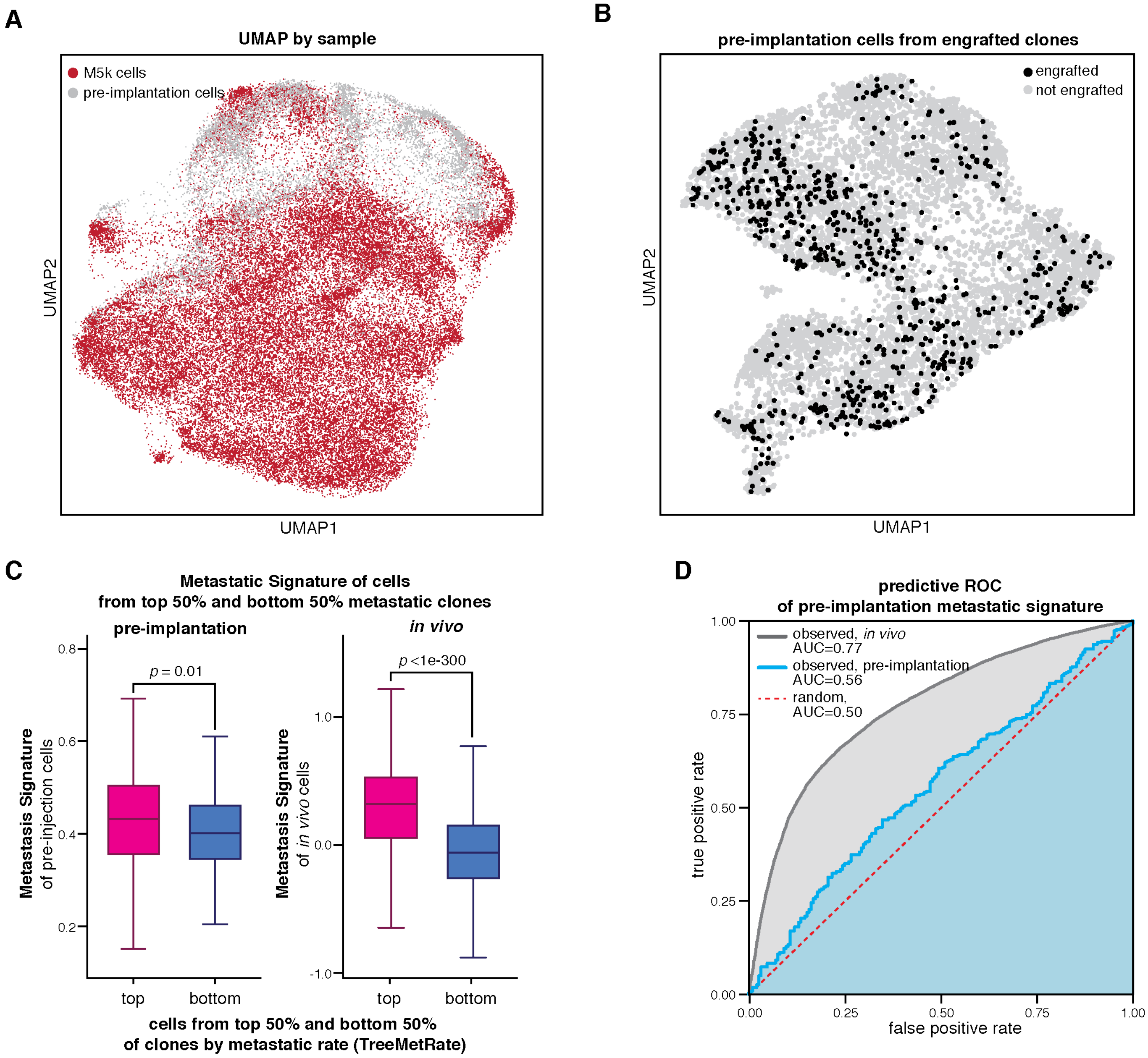


**Fig. S19. The cells in the pre-implantation pool heterogeneously express metastasis-associated genes, which are modestly predictive of their *in vivo* metastatic phenotype.** (**A**) Projection of transcriptional states of M5k and pre-implantation cells, colored by sample, as in **Fig. 5A**. (**B**) Some of the cells from the pre-implantation pool could be assigned to the ~100 clonal populations that engrafted and proliferated in mouse M5k based on their clonal barcodes (i.e., intBCs). Shown are the pre-implantation cells that could be assigned to an engrafted clone (black) on a projection of pre-implantation transcriptional states, as in **Fig. 5B,C**. (**C, left**) Pre-implantation cells from the top 50% (most) metastatic clones *in vivo* have higher Metastatic Signature scores than pre-implantation cells from the bottom 50% (least) metastatic clones *in vivo* (Mann-Whitney $U$ $p$-value=0.01). (**C, right**) The Metastatic Signature of the most and least metastatic clones is more pronounced *in vivo* than in the pre-implantation cells ($p$-value<1e-300). (**D**) For the pre-implantation cells, the difference in Metastatic Signature scores between the most and least metastatic clones is modest, yet significant by ROC (receiver operator characteristic) analysis of false positives vs. true positives (area under the curve, AUC=0.56), indicating that the Metastatic Signature score pre-implantation is a modest predictor of *in vivo* metastatic phenotype. The predictive power for the *in vivo* population of cells is greater (AUC=0.77).

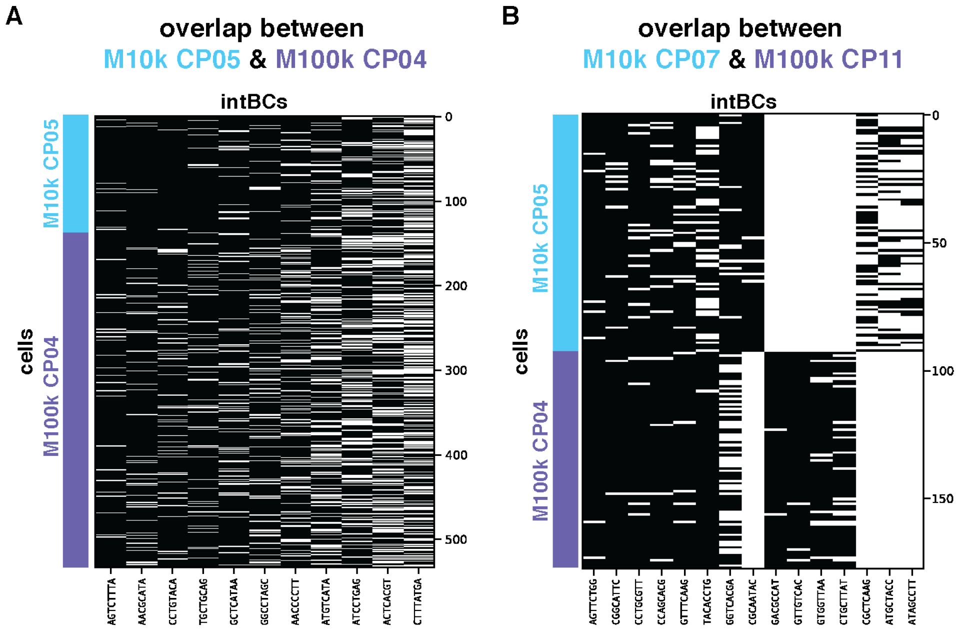


**Fig. S20. Two pairs of clonal populations from mice M10k and M100k are related, enabling an experiment to determine the robustness and reproducibility of metastatic phenotype across independent mouse experiments.** Each intBC (columns) observed for each cell (rows) from (**A**) M10k CP05 and M100k CP04 and (**B**) M10k CP07 and M100k CP11. Cells from M10k are shown in light blue; M100k in purple. The clonal populations in each of these pairs are related to one another based on their shared sets of intBCs, as in **Fig. 5D**. The paired clonal populations here are the most closely related between the two mouse experiments.

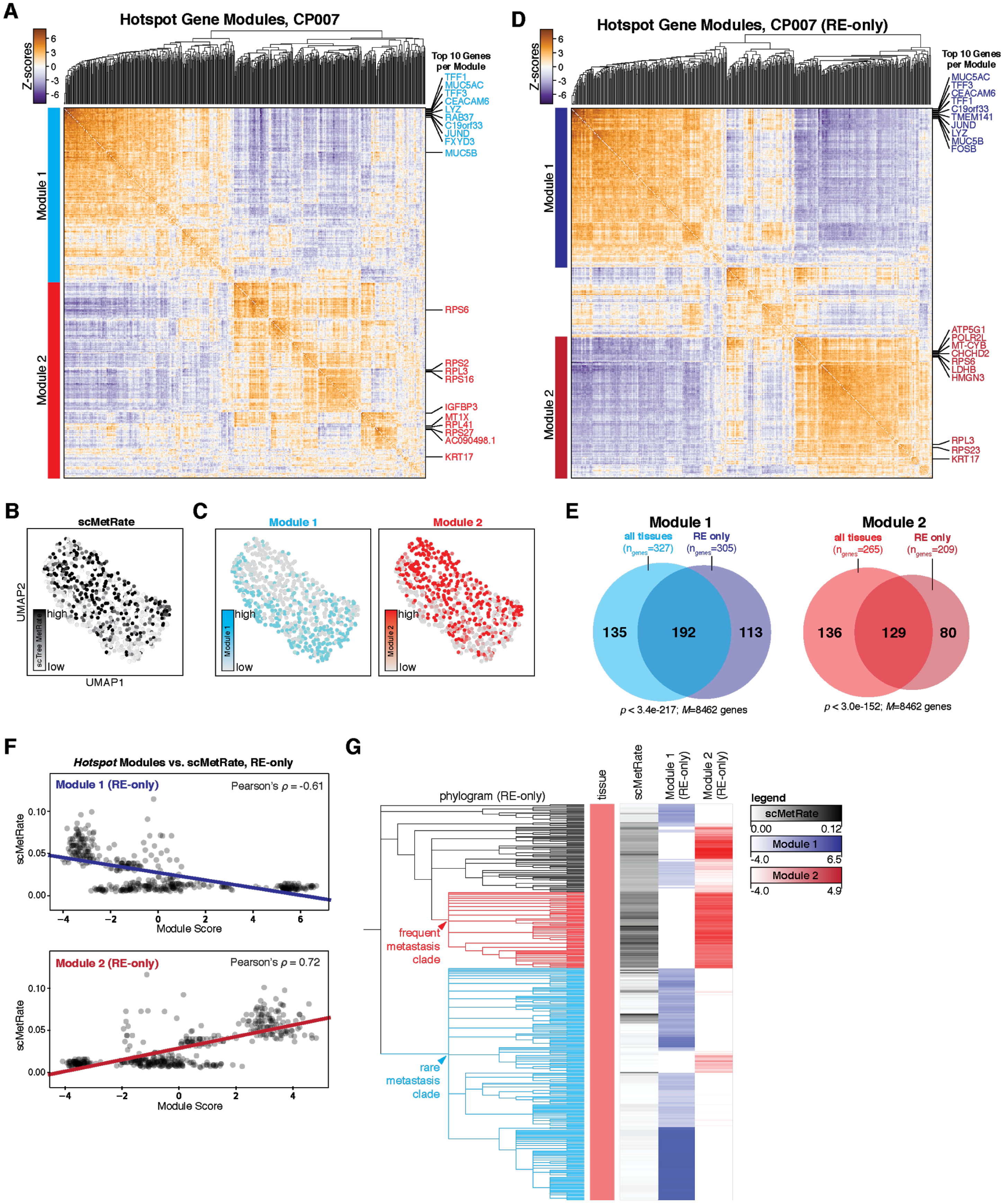


**Fig. S21. Distinct transcriptional modules underlie distinct clade-specific metastatic behaviors in Clone #7.** (**A**) *Hotspot* analysis identifies two gene modules that have heritable expression patterns in CP007 (Modules 1 and 2; indicated in cyan and red, respectively). Pairwise local correlations of genes (with FDR < 0.1) calculated with *Hostpot* are shown by heat-map; the top 10 most significant genes each for Modules 1 and 2 are annotated in cyan and red, respectively. (**B** and **C**) The transcriptional states for all cells in CP007 represented in a two-dimensional projection (UMAP) and colored by scMetRate (**B**) or *Hotspot* Module scores (**C**). (**D**) When restricting *Hotspot* analysis to only cells from the "RE" tissue sample in CP007, two heritable gene modules are identified (dark blue and dark red). As before, the top 10 genes from each module are annotated. (**E**) The gene modules identified from all cells or from only RE-only cells in CP007 overlap significantly by hypergeometric test. Number of genes in each set ($n_{genes}$), number of expressed genes in at least 10% of cells in CP007 ($M$), and $p$-value of the hypergeometric test are indicated. (**F**) RE-only Modules 1 and 2 are negatively and positively associated with the scMetRate, respectively, as in the analysis for all cells from CP007 (**Fig. 5I**). (**G**) Overlay of the phylogram for RE-only cells from CP007, scMetRate, and *Hotspot* module scores (RE-only), showing concordance between the frequently metastasizing clade (red) and Module 2 and the rarely metastasizing clade (cyan) and Module 1, as in **Fig. 5J.**

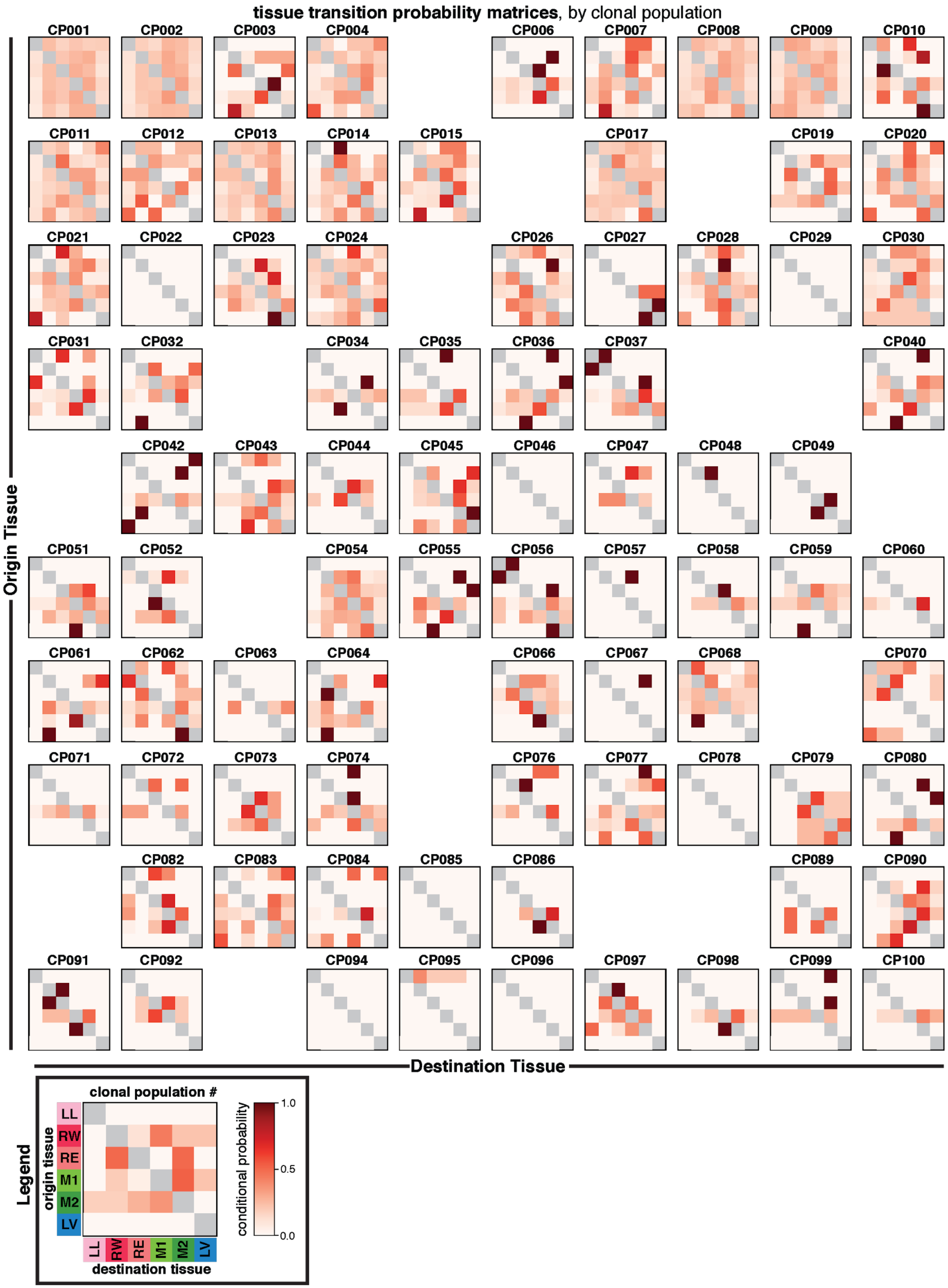
tissue transition probability matrices, by clonal population
CP001
CP002
CP003
CP004
CP006
CP007
CP008
CP009
CP010
CP011
CP012
CP013
CP014
CP015
CP017
CP019
CP020
CP021
CP022
CP023
CP024
CP026
CP027
CP028
CP029
CP030
CP031
CP032
CP034
CP035
CP036
CP037
CP040
CP042
CP043
CP044
CP045
CP046
CP047
CP048
CP049
CP051
CP052
CP054
CP055
CP056
CP057
CP058
CP059
CP060
CP061
CP062
CP063
CP064
CP066
CP067
CP068
CP070
CP071
CP072
CP073
CP074
CP076
CP077
CP078
CP079
CP080
CP082
CP083
CP084
CP085
CP086
CP089
CP090
CP091
CP092
CP094
CP095
CP096
CP097
CP098
CP099
CP100
Origin Tissue
Destination Tissue
Legend
clonal population #
origin tissue
LL
RW
RE
M1
M2
LV
destination tissue
conditional probability
1.0
0.5
0.0

**Fig. S22. Tissue transition probability matrices for each clonal population.** The conditional probability of transition from and to each tissue inferred from the phylogenetic trees (calculated with *FitchCount)* of each clonal population, thus summarizing the most probable tissue routes of metastasis. Legend (lower left) indicates the color bar showing conditional probability and the tissue labels, as in **Fig. 1E**. Notably, the transition matrices are varied and distinct to each clonal population.

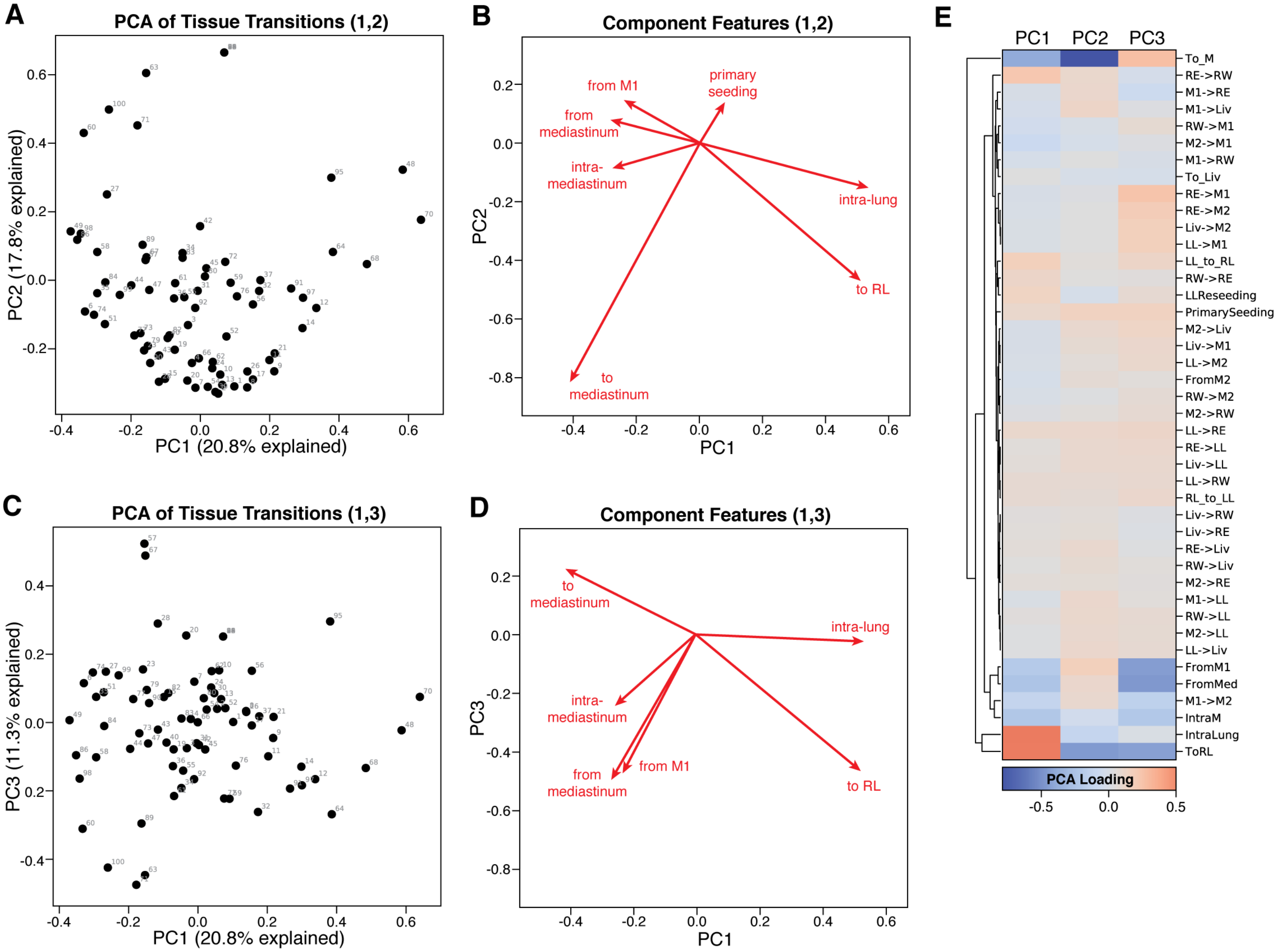


**Fig. S23. Describing the principal features of metastatic seeding routes.** (**A**, **C**) PCA projections of the metastatic tissue transitions for each clonal population (annotated). The percentage of the variance explained by each component is indicated on the axes for the first and second (**A**) or first and third (**C**) components. (**B**, **D**) Biplot vectors representing the most explanatory features of the first, second, and third principal components, annotated by descriptive features of metastatic transitions. The length and angle of the vector describe the scale and direction, respectively, of each descriptive feature. (**F**) The PCA loadings of the metastatic transition features for each principal component.

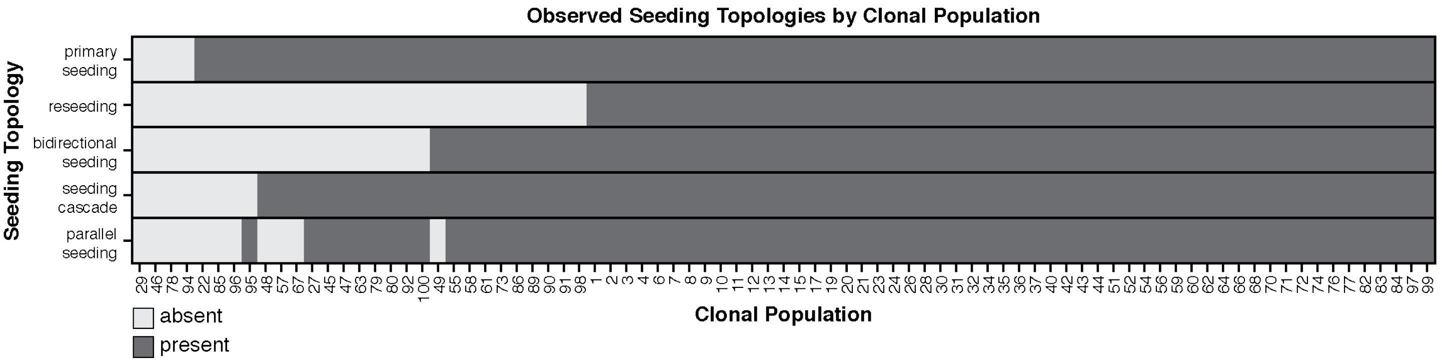


**Fig. S24. Seeding topologies observed in each clonal population.** A table describing classified seeding topologies (rows) that are present or absent (dark or light gray, respectively) in each clonal population (columns). The majority of clonal populations exhibit examples of all seeding topologies.

**Supplementary Text:** (embedded below)

### *FitchCount:* an Efficient Algorithm for Inferring Transition Matrices on Phylogenetic Trees

April 16, 2020

#### Contents


**1 Deriving transition matrices from phylogenetic trees** **1**

**2 Algorithmic strategy** **2**

**3 Finding the minimal number of transitions** **3**

**4 Inferring the frequency of different state transition events** **3**

4.1 The algorithm . . . . . . . . . . . . . . . . . . . . . . . . . . . . . . . . . . 4

4.2 Proof . . . . . . . . . . . . . . . . . . . . . . . . . . . . . . . . . . . . . . . . 5

**5 Appendix** **8**

**References** **10**


#### 1 Deriving transition matrices from phylogenetic trees

---

**function** OPT($node = v$)
    **if** is_leaf(v) **then**
        **return** $\{state(v)\}$ ▷ for leaves return their assignment, which was provided as input
    $\forall s \in \Sigma, c(s) = \#\{u \in child(v) \text{ s.t. } s \in opt(u)\}$
    $k = max_{s \in \Sigma} c(s)$
    $n(v) = \mid child(v) \mid - k + sum_{u \in child(v)} n(u)$
    **return** $\{s \in \Sigma \text{ s.t. } c(s) = k\}$

---

Note that in the original formulation by Hartigan, an additional complication is added to increase the number of optimal assignments that can be retrieved by the algorithm. This is done by accumulation of an additional set of states $opt_2(v)$ for every node $v$ that can be used to derive solutions that are globally optimal, but are not optimal in the sub-tree rooted by $v$. We therefore excluded this part of the algorithm (see appendix for proof [Claim 3]).

#### 4 Inferring the frequency of different state transition events

The second part of the Fitch-Hartigan algorithm is a top-down procedure for finding one (out of potentially many) optimal solution, i.e., a labeling $state : V \to \Sigma$ of each node $v \in V$ in the tree with a state $s \in \Sigma$. It starts by randomly selecting a state for the root node $r$ out of the set $opt(r)$ and then continues to select legal (see Definition 1) states for child nodes, based on the value assigned to their parent.

---

**function** STATE-ASSIGNMENT($node = v$)
    **if** is_root(v) **then**
        $state(v)$ = random selection out of $opt(v)$
    **else**
        **if** $state(parent(v)) \in opt(v)$ **then** $state(v) = state(parent(v))$
        **else**$state(v)$ = random selection out of $opt(v)$

---

The above procedure can face many ties during its execution, and can thus potentially return all optimal solutions (provided that they remain optimal for every sub-tree) if it is applied many times. When we compare Fitch-Hartigan to FitchCount in the main text, we use one top-down round (i.e., consider one optimal solution) for the former.

The *FitchCount* procedure was designed to provide a comprehensive evaluation of state transition frequencies, by basing its estimation on the space of all possible optimal state assignments, instead of only a single or few optimal assignments. Compared to the naive approach to enumerate all possible optimal state assignments given by the Fitch-Hartigan algorithm (which may require exponential number of executions), *FitchCount* performs in $\mathcal{O}(n \cdot k^3)$ time for a tree with $n$ leaf nodes and $k$ possible states (assuming each internal node has at least two child nodes, which is the case in our work).

#### 4.1 The algorithm

We define several arrays for storing necessary information:

1. $opt[v]$: The set of optimal assignments for a node $v$ given by the Fitch-Hartigan bottom up approach procedure (defined in the algorithm *opt*).
2. $N[v, s]$: The number of optimal solutions below the node $v$ given that it takes on the state $s$.
3. $C[v, s, s_i, s_j]$: The number of transitions from state $s_i$ to state $s_j$ in all optimal solutions of the tree rooted at $v$, given that $v$ takes on the state $s$.
4. $M[i, j]$: The number of transitions between $s_i$ and $s_j$ observed across all optimal solutions to the Fitch-Hartigan algorithm.

Our overall objective is to fill in the dynamic programming matrix $M$, which will subsequently require knowledge of the other dynamic programming arrays. Note that in the following we refer to the arrays using either rounded parenthesis or rectangular parenthesis. The former denotes a function call and the latter denotes a retrieval of an already-computed entry (which we assume to get populated automatically after the respective function call and available globally to the algorithms). Our Main function is:

---

**function** MAIN($tree = T$, $states = \Sigma$).
    $r$ = root of $T$
    Call opt(r)    ▷ Note that opt(r) fills out the array opt in post-order from the root
    **for all** $s \in opt[r]$ **do**
        Call $N(r, s)$
    **for all** $s \in opt[r]$ **do**
        **for all** $\{s_i, s_j\} \in \Sigma^2$ **do**
            Call $C(r, s, s_i, s_j)$
    **for all** $\{s_i, s_j\} \in S^2$ **do**
        $M[s_i, s_j] = sum(C[r, :, s_i, s_j])$
    **return** $M$

---

The following algorithms for filling in specific entries to $N[v, s]$, and $C[v, s, s_i, s_j]$:

```
function N(node = v, state = s)
    if is_leaf(v) then
        return 1
    A = []  ▷ an array storing the number of solutions in each subtree below v given its state s
    for all u ∈ child(v) do
        LS = ∅                          ▷ The set of legal states for node u given state(v) = s
        if s ∈ opt[u] then
            LS = {s}
        else
            LS = opt[u]
      A[u] = ∑_{s'∈LS} N(u, s')
    return ∏_{u∈child(v)} A[u]

function C(node = v, state = s, from = s_i, to = s_j)
    if is_leaf(v) then
        return 0
    K = []        ▷ A temporary array to store the number of transitions observed for each child
    for all u ∈ child(v) do
        LS[u] = ∅                       ▷ The set of legal states for node u given state(v) = s
        if s ∈ opt[u] then
            LS[u] = {s}
        else
            LS[u] = opt[u]
        K[u] = ∑_{s'∈LS[u]} C(u, s', s_i, s_j)
        if (s_i == s) ∧ (s_j ∈ LS[u]) then
            K[u]+ = N[u, s_j]
    return ∑_{u∈child(v)} ( K[u] ∏_{u'∈child(v)\{u}} ( ∑_{s'∈LS[u']} N[u', s'] ) )
```

#### 4.2 Proof

We prove correctness of the dynamic programming matrices $N$, and $C$.

**Claim 1.** *For any node $v$ and state $s \in opt[v]$, $N[v, s]$, is precisely the number of optimal solutions in $T^{(v)}$ where $state(v) = s$.*

*Proof.* To prove the correctness of the dynamic programming array $N$, we'll proceed by induction over the height of the tree $height(T) = h$. For convenience, we'll also make use of a temporary dynamic programming array $A$ which stores the number of solutions for each child $c \in child(v)$, aggregated across each possible state of that child given the parent's state $s$.

<u>Base Case #1, $h = 0$</u>. $N[l, state(l)] = 1$. This relation is trivially true for the case where $h = 0$ and there exists a single leaf $l_1$ in which case the only solution consists of $state(l) = s$.

<u>Base Case #2, $h = 1$</u>. Consider a tree $T^{(v)}$ rooted at node $v$, where $child(v) = \{l_1, ..., l_m\}$. We know that each leaf $l_i$ has a single assignment $state(l_i)$ from the definition of the small-parsimony

problem. Thus, the number of possible solutions for this tree with $state(v) = s$ is always one, namely with each leaf taking on their only state. Specifically, in this base case, we observe that $A[u] = 1 \forall u \in \{l_1, .., l_m\}$. To show this, we consider two cases:

- If $s \notin opt(l_i)$: $A[l_i] = \sum_{s' \in opt[l_i]} N[l_i, s'] = N[l_i, state(l_i)] = 1$ since $height(T^{(l_i)}) = 0$.
- If $s \in opt[l_i]$: $s == state(l_i)$ and $A[l_i] = N[l_i, s] = 1$ since $height(T^{(l_i)}) = 0$.

and the relation

$$N[v, s] = \prod_{u \in child(v)} A[u] = A[l_1] * ... * A[l_m] = 1$$

thus is correct.

Inductive Hypothesis. For a tree $T^{(v)}$ of height $h$, and some $s \in opt[v]$, $N[v, s]$ exactly stores the number of optimal solutions in the tree rooted at $v$.

Inductive Step. Consider a tree $T^{(v)}$ of height $h + 1$ and some state $s \in opt[v]$. We will show that both the array $A$ correctly stores the number of solutions for the child $u$ given $state(v) = s$ and that the relation $N[v, s] = \prod_{u \in child(v)} A[u]$ is correct.

First, we note that for the tree to be globally optimal, for each $u \in child(v)$, $state(u)$ must be $s$ if $s \in opt[u]$; else, any state from $opt[u]$ can be assigned to $u$ as each incurs a cost of 1 to the overall parsimony of the tree (see Claim 3). These choices for "optimal" states are stored in the array $LS[u]$.

Second, we know from our inductive hypothesis that $N[u, s']$ is correct for any child $u \in child(v)$ and any state $s' \in opt[u]$ as the tree $T^{(u)}$ has a height $h$. Thus, it is clear that $A[u] = \sum_{s' \in LS[u]} N[u, s']$ correctly returns the number of solutions in the subtree rooted at $u$ over all possible legal states that $u$ can take on.

Finally, we observe that given $state(v) = s$, each child can be treated independently as we consider global solutions that in the tree $T^{(v)}$ with $state(v) = s$. Because of this, the number of such solutions is the size of the permutation of all optimal sub-trees rooted at each $u \in child(v)$ - i.e. the product of all $A[u]$. To show this, consider $v$ has $m$ children. Let the set of optimal internal labellings to $T^{(u_j)}$ be denoted as $\tau_j = \{t_i^{(j)}\}_{i=1}^{A[u_j]}$ where $t_i^{(j)}$ is the $i^{th}$ solution for the tree rooted at $u_j$ given $state(v) = s$. Then, the possible set of solutions is the Cartesian Product between $\tau_1, \tau_2, ..., \tau_m$:

$$\tau_1 \times ... \times \tau_k = \Big\{\{t_1^{(1)}, t_1^{(2)}\}, \{t_2^{(1)}, t_1^{(2)}\}, ..., \{t_{A[u_1]}^{(1)}, t_1^{(2)}\}, ..., \{t_1^{(m-1)}, t_2^{(m)}\}, ..., \{t_{A[u_{m-1}]}^{(m-1)}, t_{[A[u_m]}^{(m)}\}\Big\}$$

This Cartesian product has a size $|\tau_1| \times |\tau_2| ... \times |\tau_m| = A[u_i] \times ... \times A[u_k] = \prod_{u \in child(v)} A[u]$. Thus, this relation holds $T^{(v)}$ where $height(T^{(v)}) = h + 1$. □

**Claim 2.** *For any node $v$ and state $s \in opt[v]$ assigned to $v$ and $\{s_i, s_j\} \in 2^{\Sigma}$, the array $C[v, s, s_i, s_j]$ correctly stores the number of transitions from $s_i \to s_j$ in $T^{(v)}$.*

*Proof.* We will prove by induction over the height of the tree, $h$, that for a node $v$, a state $state(v) = s$, and some $(s_i, s_j) \in 2^{\Sigma}$ both $K[u] \forall u \in child(v)$ and

$$C[v, s, s_i, s_j] = \sum_{u \in child(v)} \left( K[u] \prod_{u' \in child(v) \setminus \{u\}} \Big( \sum_{s' \in LS[u']} N[u', s'] \Big) \right)$$

are correct. Here $K[u]$ is the number of transitions from $s_i$ to $s_j$ that exist by considering child $u$ of node $v$ given $state(v) = s$ and $LS[u]$ is a function that finds the set of legal (Definition 1) assignments to $u$ given the parent's state is $s$.

Base Case #1, $h = 0$. The relation trivially holds for a tree of height 0 as there cannot exist any transitions for a tree without edges. As calculated, $C[v, s, s_i, s_j] = 0$ for all leaves and thus the relation holds.

Base Case #2, $h = 1$. Consider a tree of height 1, $T^{(v)}$, where $child(v) = \{l_1, ..., l_m\}$ and that $N[l_i, state(l_i)] = 1$. We can count the number of transitions by considering for every edge the following:

- $s \neq s_i$ then $C[v, s, s_i, s_j]$ is necessarily 0.
- $s == s_i$, then $C[v, s, s_i, s_j]$ is the number of leaves that have state $s_j$.

By construction, for some child $l_i$, $C[v, s, s_i, s_j]$ must be

$$K[l_i] = 1[s == s_i \wedge s_j \in LS[u]] == 1[s == s_i \wedge s_j == state(l_i)]$$

Then, $C[v, s, s_i, s_j] = \sum_{l \in child(v)} K[l_i]$. We'll prove that this is equal to the relation described above:

$$\begin{aligned} C[v, s, s_i, s_j] &= \sum_{l \in child(v)} \left( K[l] \prod_{l' \in child(v) \setminus \{l\}} \left( \sum_{s' \in LS[u']} N[u', s'] \right) \right) \\ &= \sum_{l \in child(v)} \left( K[l] \prod_{l' \in child(v) \setminus \{l\}} 1 \right) \\ &= \sum_{l \in child(v)} K[l] \end{aligned}$$

Thus, our relation holds for $h = 1$.

Inductive Hypothesis. Assume for a tree $T^{(v)}$ of height $h$ where $state(v) = s$, $C[v, s, s_i, s_j]$ correctly computes the number of transitions from $s_i, s_j \in 2^{\Sigma}$ for the tree.

Induction Step. Now consider a tree of height $h + 1$ rooted at $v$ where $state(v) = s$. We'll show that both $K[u]$ is correct for all $u \in child(v)$ and that the relation for calculating $C[v, s, s_i, s_j]$ holds.

We'll first show that $K[u]$ is correct $\forall u \in child(v)$. As defined above, $K[u]$ is the number of $s_i \to s_j$ transitions that are due to the node $u$ given $state(v) = s$. We know that given our inductive hypothesis, for the subtree rooted at $u$, $T^{(u)}$, $C[u, s', s_i, s_j]$ for any state $s' \in opt[u]$ is correct. Then, the number of $s_i \to s_j$ transitions under $u$, given $state(v) = s$, is equal to the sum of all $C[u, s', s_i, s_j]$ for those $s' \in LS[u]$ as those are the only solutions that would be considered by the Fitch-Hartigan algorithm (note that we are guaranteed to have optimal state assignments to chose from for $LS[u]$ as we prove in Claim 3), plus the transition (if it exists) from $v$ to $u$.

Now, we'll show that the relation for $C[v, s, s_i, s_j]$ holds. For the tree $T^{(v)}$ let's assume that $v$ has $m$ children: $child(v) = \{u_i\}_{i=1}^{m}$. As above, we'll maintain the notation that the set of legal assignments given that $state(v) = s$ for $u_i$ to be $LS[u_i]$. Furthermore, let the set of $s_i \to s_j$ transitions underneath $u_j$ be $\rho_j = \{r_i^{(j)}\}_{i=1}^{K[u_j]}$ and the set of trees that are legal, optimal assignments under $u_j$ be $\tau_j = \{t_i^{(j)}\}_{i=1}^{\Lambda(u_j)}$ where $\Lambda(u_j) = \sum_{s' \in LS[u_j]} N[u_j, s']$, assuming $state(v) = s$. We can see then that the total number of transitions from $s_i \rightarrow s_j$ is

$$\Big\{\{r_1^{(1)}, t_1^{(2)}\}, \{r_2^{(1)}, t_1^{(2)}\}, ..., \{r_{K[u_1]}^{(1)}, t_1^{(2)}\},$$
$$\{r_1^{(1)}, t_2^{(2)}\}, ..., \{r_{K[u_1]}^{(1)}, t_{\Lambda(u_2)}^{(2)}\}, ..., \{r_{K[u_1]}^{(1)}, t_{\Lambda(u_m)}^{(m)}\},$$
$$\{r_1^{(2)}, t_1^{(1)}\}, ..., \{r_{K[u_2]}^{(2)}, t_{\Lambda(u_m)}^{(m)}\},$$
$$\{r_1^{(m)}, t_1^{(1)}\}, ..., \{r_{A[u_m]}^{(1)}, t_1^{(2)}\}, \{r_{A[u_m]}^{(m)}, t_{\Lambda(u_{m-1})}^{(m-1)}\}\Big\}$$

Which is equal to the sum of the following Cartesian Products:

$$\rho_1 \times \{(\tau_2, ..., \tau_m)\} + \rho_2 \times \{(\tau_1, \tau_3, ..., \tau_m)\} + ... + \rho_m \times \{(\tau_1, ..., \tau_{m-1})\}$$

where the cardinality of this set is

$$K[u_1] \prod_{i \in 2..m} \Lambda(u_i) + K[u_2] \prod_{i \in 1,3,..m} \Lambda(u_i) + ... + K[u_m] \prod_{i \in 1,...,m-1} \Lambda(u_i)$$

which can be further simplified to

$$\sum_{u \in child(v)} K[u] \prod_{u' \in child(v) \setminus \{u\}} \sum_{s' \in LS[u']} N(u', s')$$

Thus the relation is correct and $C[v, s, s_i, s_j]$ is correct by induction.

□

### 5 Appendix

**Definition 1.** *(Legal Assignment). An assignment $state(v) = s$ is* ***legal*** *for a node $v$ and given $state(parent(v)) = s'$ if either $s == s'$ or $s' \notin opt(v)$. Observe that only legal assignments are explored in the Fitch-Hartigan algorithm, and are guaranteed to be optimal in the sub-tree rooted at $v$.*

□
